# Overexpression of the WWE domain of RNF146 modulates poly-(ADP)-ribose dynamics at sites of DNA damage

**DOI:** 10.1101/2023.12.29.573650

**Authors:** Rasha Q. Al-Rahahleh, Kate M. Saville, Joel F. Andrews, Zhijin Wu, Christopher A. Koczor, Robert W. Sobol

## Abstract

Protein poly-ADP-ribosylation (PARylation) is a post-translational modification formed by transfer of successive units of ADP-ribose to target proteins to form poly-ADP-ribose (PAR) chains. PAR plays a critical role in the DNA damage response (DDR) by acting as a signaling platform to promote the recruitment of DNA repair factors to the sites of DNA damage that bind via their PAR-binding domains (PBDs). Several classes of PBD families have been recognized, which identify distinct parts of the PAR chain. Proteins encoding PBDs play an essential role in conveying the PAR-mediated signal through their interaction with PAR chains, which mediates many cellular functions, including the DDR. The WWE domain identifies the iso-ADP-ribose moiety of the PAR chain. We recently described the WWE domain of RNF146 as a robust genetically encoded probe, when fused to EGFP, for detection of PAR in live cells. Here, we evaluated other PBD candidates as molecular PAR probes in live cells, including several other WWE domains and an engineered macrodomain. In addition, we demonstrate unique PAR dynamics when tracked by different PAR binding domains, a finding that that can be exploited for modulation of the PAR-dependent DNA damage response.

## 1. Introduction

ADP-ribosylation is a reversible, covalent post-translational modification of proteins [1-4], as well as a modification of DNA [5] and RNA [6, 7], catalyzed by the ADP-ribose polymerase family of enzymes [1]. These modifications are defined as those adding a single ADP-ribose moiety (mono-ADP-ribose polymerases, MARPs) or multiple ADP-ribose moieties (poly-ADP-ribose polymerase, PARPs) and are overall defined as ADP-Ribosyl-Transferase Diphtheria Toxin-Like proteins (ARTDs) [8, 9]. ADP-ribosylation occurs either as: (1) mono-ADP-ribosylation, in which only one ADP-ribose unit is transferred to residues of target proteins, or (2) poly-ADP-ribosylation (PARylation) in which several units of ADP-ribose are successively transferred to specific residues within the target proteins to constitute poly-ADP-ribose (PAR) chains [1, 10]. Side chains of glutamate, aspartate, serine, arginine, cysteine, lysine and asparagine were reported as ADP-ribose acceptors [11]. Recently, ADP-ribosylation (ADPr) of serine residues has been linked with DNA damage-induced modifications catalyzed by PARP1 and/ or PARP2 when in complex with HPF1 [12]. For details on the ADP-ribose modification of DNA and RNA molecules, see [5-7, 13-16].

PARylation regulates several biological processes such as the DNA damage response (DDR), chromatin reorganization, transcription, mitosis, and apoptosis [17, 18]. Of the seventeen proteins of the PARP family of enzymes, PARP1, PARP2 and PARP3 were reported to be involved in the DNA damage response and in DNA repair [2, 19, 20]. PARP1 and PARP2 sense DNA breaks [8, 21], and once a DNA single-strand break or double-strand break occurs, PARP1 and PARP2 are activated and catalyze the successive transfer of ADP-ribose units from nicotinamide adenine dinucleotide (NAD^+^) to the side chains of their substrates, forming PAR chains [22-25]. PARylation is a key player in the DDR in which PAR promotes the recruitment of several DNA repair proteins to the site of DNA damage, as mediated via their PAR-binding domains (PBDs) [9, 18, 26-29]. The resulting DNA repair complex is then disassembled once the damage is resolved, and PAR is degraded [2, 30, 31].

Hence, like other covalent post-translational modifications, PARylation is a dynamic process that initiates within seconds and has been shown to have a half-life of a few minutes or more [32, 33]. Mono- and poly-ADP-ribosylation are reversed by PAR degrading enzymes. Six human dePARylation proteins have been recognized so far, with poly-ADP-ribose glycohydrolase (PARG) shown to be highly correlated to the DDR [34, 35]. First identified in both bovine and human cells [36, 37], PARG has been an enzyme of significant interest, both for biological characterization of mono- and poly-ADP-ribose metabolism and cellular function [38, 39] and the role of the PARP/PAR/PARG axis in cancer [31, 40, 41]. PARG possesses both exo-glycohydrolase and endo-glycohydrolase activities to hydrolyze the ribose-ribose bonds within PAR [42], releasing free PAR chains or mono-ADP-ribose moieties [36, 43].

PBDs have been appreciated as important mediators or readers of PAR to transduce the signal [44]. To date, ten different PBD families have been identified, each displaying different binding affinities as well as recognition dynamics for distinct parts within PAR chains [9, 44, 45]. The first discovered PAR recognition domain is the PAR binding motif (PBM) [46], that has been refined to a consensus eight amino acid motif [47], recognized in almost all characterized PAR-binding proteins [46]. To-date, over 800 human proteins have been shown to contain a PBM [47].

Several classes of PAR binding molecules have been studied such as the WWE domain, a conserved globular domain that was originally detected in several proteins in the E3 ubiquitin ligase family such as Deltex and TRIP12 and in some poly-ADP-ribose polymerase (PARP) homologs [48]. Another is the macrodomain, a large globular PAR-binding domain identified in some PARP family members, in histone variants, and in many of the PAR degrading enzymes [44, 49, 50]. Further, there is the PAR binding Zinc finger (PBZ) domain, a zinc finger domain encoded in APLF and CHFR [28], in addition to other less PAR-specific PBD families [44, 51, 52]. Defining PAR binding characteristics of PBDs has been an attractive field of research, as PBDs were used to elucidate cellular functions associated with ADP-ribosylation via affinity purification of ADP-ribosylated proteins [53, 54] and more recently as molecular probes to track PAR dynamics in live cells [55-59]. However, the binding kinetics of PBDs to PAR is not yet fully explored, as this differential binding may play a role in selectivity of PAR signal transduction as well as in the regulation of the PARylation response.

Fluorescent microscopy has emerged as a valuable tool for visualization of cellular architecture, to determine protein localization, in addition to examining protein interactions [60]. Advances in laser scanning-confocal microscopy, followed by the introduction of laser wavelengths with DNA damage-inducing capability, has enhanced our knowledge of the DDR and the dynamics of DNA repair factor recruitment to DNA damage sites [55, 56, 61, 62]. To evaluate the assembly and dissociation of DNA repair factors at DNA damage sites, imaging can be performed in live cells expressing fluorescently labeled DNA repair factors or in fixed cells using immunofluorescence [2]. In contrast to *in vitro* biochemical techniques, which have been extensively used for the study of proteins in the base excision repair (BER) and single-strand break repair (SSBR) pathways and other DNA repair pathways [63, 64], laser micro- irradiation offers the temporal resolution needed to evaluate the association and dissociation of key repair proteins at DNA damage sites by analyzing the accumulation of the protein of interest or its modification over a selected duration following the induction of DNA damage [65].

In that context, the quantification of PAR formed at DNA damage sites in live cells has been pursued by our lab [55, 56, 66], and many others [55-59]. PBDs fused to EGFP can be effectively used as probes for PAR detection and tracking [55, 56, 58, 59]. We have recently reported a genetically encoded fusion of the WWE domain encoded in the RNF146 gene as a molecular tool to detect PAR in live cells at sites of laser micro-irradiation induced DNA damage [55], following hydrogen peroxide (H_2_O_2_) or methyl methane sulfonate (MMS) treatment [66] and to screen small molecule genotoxins, PARP inhibitors, and PARG inhibitors [66]. However, whether other WWE domains share similar live-cell PAR detection characteristics has not been fully vetted. Here, we evaluated the ability of other WWE domains to detect PAR in live cells. Further, we evaluated a recently reported high affinity, engineered macrodomain [50], as a molecular probe to detect PAR in live cells. Finally, we explored the dynamics of PAR formation and degradation as detected by these different PAR probes.

## 2. Materials and Methods

### 2.1 Reagents and chemicals

All plasmids, chemicals, and other reagents used in this study are listed in **Table S1**.

### 2.2 Cells and cell culture

U2OS and ES-2 cells were obtained from ATCC. LN428 cells were a kind gift from Dr. Ian Pollack (University of Pittsburgh) and have been described by us previously [40, 67]. 293-FT cells were obtained from Thermo Fisher Scientific. U2OS cells and 293-FT cells were cultured in DMEM supplemented with 10% heat-inactivated fetal bovine serum, penicillin/streptomycin, and L-glutamine. ES-2 cells were cultured in McCoy’s 5A medium (1X) supplemented with 10% heat-inactivated fetal bovine serum, and penicillin/streptomycin. LN428 cells were cultured in MEM Alpha medium (1X) supplemented with 10% heat-inactivated fetal bovine serum, L-glutamine, and penicillin/streptomycin/amphotericin. All parental and modified cell lines were grown in tissue culture incubators at 37°C, 5% CO_2_.

### 2.3 Plasmid and vector development

To generate lentiviral vectors encoding poly-(ADP-ribose) (PAR) binding domains (PBD) fused to EGFP, a lentiviral backbone vector was designed in-house and then purchased from VectorBuilder (Chicago, IL). As designed, this backbone vector (pLV-Hygro-EF1A- LivePARBackbone) contains a Gly-Ser linker, an EGFP open reading frame and BamHI and MluI restriction sites used to allow in-frame cloning of Af1521 variants or the WWE domains encoded by RNF146, TRIP12, Deltex2 or others as needed. To generate lentiviral vectors encoding a PBD fused to a myc tag, a lentiviral backbone vector was designed in-house and then purchased from VectorBuilder (Chicago, IL). As designed, this backbone vector, pLV-Hygro-EF1A-WWE-Linker-EGFP, contains BamHI and XpaI restriction sites used to allow in-frame cloning of myc tag-fused WWE domains and Af1521 variant fragments and elimination of the EGFP open reading frame, if needed. DNA fragments of each of the AF1521 variants [50], various WWE domains and myc-tagged PBDs were designed in-house and then purchased from Twist (South San Francisco, CA). Next, each DNA fragment was then ligated into the designated restriction-digested lentiviral backbone vector. Selection of positive clones, and plasmid extraction was followed by purification using the QIAprep Spin Miniprep Kit (Qiagen). Sequences were validated by Sanger sequencing (Eurofins Genomics). The complete list of vectors used and developed for this study is listed in **Table S1**.

### 2.4 Lentivirus production and cell transduction

293-FT cells were used to produce lentiviral particles by co-transfecting four plasmids using the TransIT-X2 transfection reagent composed of: the packaging vectors pMD2.g(VSVG), pVSV-REV and pMDLg/pRRE together with the desired shuttle vectors, as listed in **Table S1**. Supernatant containing lentivirus was collected after forty-eight hours and was passed through 0.45 mM filters to separate the viral particles as described previously [68]. For lentiviral transduction, 1-2 x 10^5^ cells were seeded into each well of a 6-well plate and cultured at 37°C, 5% CO_2_. After 24 hours, lentiviral particles (1ml) were mixed with growth medium (1ml) supplemented with polybrene (2μg/ml) and then added to the cells. Cells were incubated at 32°C, 5% CO_2_ overnight, followed by removal of medium containing the lentiviral particles and replacement with fresh medium. For stable cell lines (e.g., cells expressing a myc-tagged PBD), cells were cultured for 48 hours at 37°C followed by selection with antibiotics (hygromycin) for 1–2 weeks. For the second transduction for the creation of cells expressing a fluorescently tagged fusion protein in addition to expressing a myc-tagged PBD, selection for the first stable cell line was completed and expression validated prior to starting the second transduction. For experiments with short-term expression, cells were cultured for at least 96 hours following transduction at 37°C, and experiments were completed in less than two weeks. All stable cell lines developed and used in this study (along with media formulations) are listed in **Table S1**.

### 2.5 Cell protein extract preparation and immunoblot

Protein extracts (whole cell lysates) were prepared from cells expressing different proteins and/or treated with various chemicals as indicated in the text and/or figure legends. First, cells were cultured in a 60-mm cell culture plate. When the confluency reached 70-80%, cells were then washed with cold PBS, followed by vacuum aspiration of the PBS and lysis with 150μl of 2x clear Laemmli buffer (2%SDS, 20%glycerol, 62.5mmol/l Tris-HCl pH6.8). Cell lysates were heated at 95°C for 10 min and the protein concentration determined using the Nanodrop 2000C spectrophotometer [69].

Whole cell protein lysates (25-40μg protein) were loaded onto precast NuPAGE 4-12% Bis-Tris gels and then separated by electrophoresis for 70 minutes at 130V. Gel-separated proteins were transferred onto a 0.2μM nitrocellulose membrane using a Trans-Blot Turbo (Bio-Rad). Once transfer was completed, the membrane was blocked with B-TBST (TBS buffer with 0.05% Tween-20 and 5% blotting grade non-fat dry milk; Bio-Rad) for 1 hr at room temperature. Subsequently, the membrane was incubated with the primary antibodies in B-TBST overnight at 4°C. The primary antibodies and their dilutions are listed in **Table S1**. After washing with TBST (3x10 minutes), membranes were incubated with secondary antibodies in B-TBST for 1 hr (room temperature). HRP conjugated secondary antibodies used in the study include goat anti-mouse-HRP conjugate (Bio-Rad), goat anti-rabbit-HRP conjugate (Bio-Rad), and HRP-conjugated streptavidin (Thermo Fisher Scientific) (**Table S1**). The membrane was subjected to illumination using Immun-star HRP peroxide buffer with luminol/enhancer (Bio-Rad) and visualized using a Chemi-Doc MP imaging system (Bio-Rad).

### 2.6 Characterization of the cellular distribution of the EGFP-fused PBDs

U2OS or ES-2 cells expressing either the EGFP-fused WWE domains or EGFP-fused Af1521 macrodomain variants were cultured in wells of an 8-chamber glass bottom vessel (Thermo Fisher Scientific) at a density of 7x10^4^ cells per well. After 24 hours, cells were washed twice with 1X PBS. Next, cells were fixed with 4% formaldehyde supplemented with Triton 100-X (Thermo Fisher Scientific) for permeabilization of the cells, and Alexa Fluor 647 phalloidin (Invitrogen) to stain filamentous actin (F-actin). Cells were incubated at 4°C for 20 minutes according to the manufacturer’s protocol. Subsequently, cells were washed once with PBS and then NucBlue Fixed Ready Probes reagent (Invitrogen) was added for 20 minutes at room temperature according to the manufacturer’s protocol to stain the nuclear compartment. Cells were washed three times with 1X PBS, and then were imaged using a Nikon A1rsi laser scanning confocal microscope.

### 2.7 Laser micro-irradiation

For laser micro-irradiation, 7x10^4^ cells were cultured in wells of an 8-chamber glass bottom vessel (Thermo Fisher Scientific), as described [55, 56, 66]. After 24 hours, or after 48 hours if the sensitizing agent BrdU (10μM, 24 hrs) was used, laser micro-irradiation followed by time-lapse imaging were conducted with a Nikon A1rsi laser scanning confocal microscope equipped with 6 visible wavelength lasers (405, 441, 514, 561, 647nm, Coherent), customized to add a UVA 355nm laser (PicoQuant) controlled by a Bruker XY Galvanometer, and equipped with a live-cell incubation chamber (Tokai Hit) maintained at 5% CO_2_ and 37°C, using a 20x (NA=0.8) non-immersion objective for 405nm laser micro-irradiation. When cells were treated with PARPi (ABT-888; 10μM) or PARGi (PDD00017273; 10μM), they were incubated for 1 hour (5% CO_2_ and 37°C) before micro-irradiation. Micro-irradiation was performed on 20-50 individual cells. Each experiment consisted of a set of 10 irradiated cells along with 2 non-irradiated controls. Two sets of experiments were conducted on any single day, as we have described [55]. Experiments were conducted at 100% laser power with a 0.125 s stimulation time per site in a parallel micro-irradiation scheme [55]. Time lapse images were taken every 15 s during a 20-minute interval. Quantification of focal recruitment within images was done using the MIDAS (<u>M</u>odular, Irradiation, <u>D</u>etection, and <u>A</u>nalysis <u>S</u>ystem) software platform to generate recruitment dynamics plots and kinetic parameters of focal recruitment, as we have described [55].

Fluorescence intensity quantification of the recruitment foci was normalized in two ways; either to the first frame of images taken at t_0_ of the time lapse experiment, as a normalization to the base level of fluorescence intensity of the region of interest (ROI) or the normalization was done by subtracting the nucleus background intensity from the ROI fluorescence intensity. Quantitative assessment of recruitment was made through three measurements: the relative peak intensity, time to peak recruitment intensity and half -life of recruitment (dissociation time). Relative peak intensity is defined as the ratio of maximum intensity to the pre-irradiation intensity of that stimulated ROI. Time to peak is defined as the time at which recruitment intensity crosses a threshold set to the 95% confidence interval of the maximum intensity for that stimulated ROI and half-life of recruitment is defined as the time post-peak at which the intensity crosses a lower threshold set to the same confidence interval of 50% of the maximum intensity. At least twenty individual cells from 2-3 fields were analyzed and used to generate recruitment profiles and kinetic parameters for each condition of expressed PBD.

### 2.8 Purification of recombinant proteins expressed in E. coli and the in vitro PAR binding assay

For the *in vitro* PAR binding assay, proteins encoding the RN146 encoded WWE domain, RNF146(100-182) or the macrodomains AF1521(WT) or AF1521(K35E/Y145R), each designed as a glutathione-S-transferase (GST) fusion protein, were expressed in, and purified from, *E. coli* (BL21-CodonPlus-RP) according to the purification scheme shown in **Figure S2A,** and as previously described [70]. In short, BL21-CodonPlus-RP cells, expressing GST- RNF146(100-182), GST-AF1521(WT), or GST-AF1521(K35E/Y145R), were used for GST-tag protein expression. Cells were cultured overnight, and then were added to fresh 1:1000 ampicillin supplemented LB medium in a 1:20 dilution. The cells were cultured until OD_600_ reached 0.6–1, followed by addition of isopropyl β-D-1-thiogalactopyranoside (IPTG, Sigma Aldrich; 0.4mM) and cells were cultured overnight at 16°C. The cell pellets were collected by centrifugation at 5000 rpm for 10 min at 4°C. The cell pellet was then washed with lysis buffer [50 mM 4-(2-hydroxyethyl)-1-piperazineethanesulfonic acid (HEPES) pH 7.4, 500mM NaCl, 1× Sigma protease inhibitor cocktail, 1 mM Dithiothreitol (DTT) and 1mM EDTA], resuspended in lysis buffer and lysed by sonication (20 s, on and 20 s, off, for 5 min). Cell lysates were then centrifuged at 10000 rpm for 1hr at 4°C to collect the cell lysate supernatant. Cell lysate supernatant was mixed with 6ml lysis buffer-washed glutathione Sepharose 4B resin, and the mixture was rotated overnight at 4°C. The resin was washed with 15ml lysis buffer four times and the washes were collected. Then, the resin was washed twice with 15ml low salt buffer (50mM HEPES pH 7.4, 100mM NaCl). GST-tagged protein was eluted with 6ml glutathione (10mg/ml, Takara) and the flow-through was collected. The column was washed again twice with low salt buffer and the flow-through was collected. Samples from each of the collected flow-through aliquots were separated by gel electrophoresis and GST-fused protein band size was confirmed (**Figure S2B**). Protein in the flow-through samples was then concentrated using Amicon Ultra centrifugal filters and the protein concentration was measured using the DC protein assay.

To investigate the binding of GST-fused PBDs to PAR *in vitro*, PAR-enriched lysates of ES-2 cells was used, prepared by treating the cells with a PARGi (PDD00017273, 10μM), for 8 hours followed by treatment with N-Methyl-N’-Nitro-N-Nitrosoguanidine (MNNG) for 30 minutes (10μM). This combined treatment was used to promote the accumulation of PARylated proteins so that the lysate can be used as a PAR source. GST pull-down of PARylated proteins using GST-tagged RNF146(100-182) or GST-tagged AF1521(K35E/Y145R) was done using the Pierce GST protein interaction pull-down kit (Thermo Fisher Scientific). Glutathione agarose resin (50μl) were washed with a 1:1 wash solution of TBS: pull-down lysis buffer (TBS is composed of 25mM Tris HCL, 0.15M NaCL, pH 7.2) according to manufacturer’s instructions. GST-tagged RNF146(100-182) or AF1521(K35E/Y145R) (100μg) were incubated with glutathione agarose resin at 4°C for 1 hour with rotation in a spin column. Resin was then washed five times with washing buffer, followed by the addition of the PAR-enriched ES-2 lysates in TBS with either 150mM or 300mM NaCl, at 4°C overnight with rotation. Next, the resin was washed five times with wash solution. GST-tagged proteins bound to PARylated proteins were eluted using 10mM Glutathione Elution Buffer (250μl) and were blotted onto nitrocellulose membrane using a slot blot (MiniFold II, Schleicher & Schuell, Inc.; Keene, NH). The membranes were then blocked with B-TBST (TBS buffer with 0.05% Tween-20 and supplemented with 5% blotting grade non-fat dry milk; Bio-Rad) for 1 hr at room temperature. Subsequently, membranes were then blotted with the primary PAR antibody in B-TBST for 2 hours at room temperature. After washing (3x10 minutes) with TBST, membranes were then incubated with the secondary antibody (HRP-conjugated goat anti-mouse secondary antibodies, BioRAD) in B-TBST for 1 hr at room temperature. After washing, the membrane was illuminated with Immun-star HRP peroxide buffer with luminol/enhancer (Bio-Rad) and imaged using a Chemi-Doc MP imaging system (Bio-Rad).

An *In vitro* PAR binding assay (Sandwich ELISA) was performed using Glutathione coated plates that were used to capture GST-tagged RNF146(100-182) or GST-tagged AF1521(K35E/Y145R) in an increasing concentration. For each well, 100μl of the assigned protein in PBS-T (Phosphate-buffered saline (PBS) supplemented with 0.05% Tween-20) was used and incubated at room temperature for 1 hour, as indicated in the figure. The wells were then washed four times with PBS-T. PAR (R&D systems; 100nM) was added to each well, followed by incubation of the plate overnight at 4°C. The next day, the plate was washed four times with PBS-T and anti-PAR primary antibody (1:200) diluted in antibody diluent solution (PBS-T, BSA) was added (50μl/well) and incubated for 2 hours at room temperature. Wells were then washed four times with PBS-T and 50μl/well of secondary antibody HRP conjugated goat anti-mouse (1:200) and the plate was incubated for 1 hour at room temperature. The wells were then washed with PBST four times and HRP substrate was added (100μl/well) and luminescence was measured with a SpectraMax M2^e^ plate reader (Molecular Devices, CA).

The scheme for the *in vitro* nitrocellulose PAR binding assay is described in **Figure S2C**. Briefly, cell lysates were prepared from ES-2 cells treated with a PARGi (PDD00017273) for 8 hours followed by treatment with N-Methyl-N’-Nitro-N-Nitrosoguanidine (MNNG) for 30 minutes (10μM), to promote the accumulation of PARylated proteins so that the lysate can be used as a PAR source. Cell lysates were collected as described above (Section 2.5). Protein concentration was measured using the DC protein assay (BioRAD). To determine the relative binding capacity of the PBD of RNF146(100-182), AF1521(WT), and AF1521(K35E/Y145R) to PAR, PAR-enriched cell lysates were blotted onto nitrocellulose using a slot blot (MiniFold II, Schleicher & Schuell, Inc, Keene, NH) and then each purified PBD GST-fusion protein was used to probe the membrane for capacity to bind to PAR. In short, PAR-enriched cell lysates (100μl) were bound, in triplicate, on a nitrocellulose membrane using vacuum suction in a decreasing concentration (**Figure S2D-F**). The membranes were then blocked with B-TBST (TBS buffer with 0.05% Tween-20 and supplemented with 5% blotting grade non-fat dry milk; Bio-Rad) for 1 hr at room temperature. Subsequently, membranes were incubated with similar amounts (20μg) of the PBD fusion-proteins in 5ml incubation buffer (50 mM HEPES pH 7.4, 100 mM NaCl) overnight at 4°C. After washing (3x10 minutes), membranes were then blotted with the primary GST-biotin antibody in B-TBST for 2 hours at room temperature. After washing (3x 10 minutes) with TBST, membranes were then incubated with the secondary antibody (HRP-conjugated streptavidin, Thermo Fisher Scientific) in B-TBST for 1 hr at room temperature. After washing, the membrane was illuminated with Immun-star HRP peroxide buffer with luminol/enhancer (Bio-Rad) and imaged using a Chemi-Doc MP imaging system (Bio-Rad).

### 2.9 Statistical analysis

For most analyses, data is shown as the mean ± standard error of the mean (SEM). Means were calculated from multiple independent experiments (n = number of independent experiments as indicated in the figure legends), unless stated otherwise. Student’s t-test, ANOVA (with a Tukey *post-hoc* test) or non-parametric tests were used to test for significant differences as appropriate, with results compared to controls and as indicated in the figure legends. For PBDs with a positive recruitment profile, exclusion criteria were set for focal recruitment quantification for PBDs that demonstrated foci recruitment as follows: foci with a relative peak intensity below 1.15 of the first frame were excluded from the experiment and from statistical analysis. Statistical analyses were performed using GraphPad Prism 8 with P-values indicated by asterisks (*p<0.05, **p<0.01, ***p<0.001, ****p<0.0001) or as stated in the figure legends.

*2.10 Software*

The software used for these studies are listed in **Table S1**.

## 3. Results

### 3.1 Variable capacity among WWE domains to detect PARylation levels and PAR dynamics in live cells

WWE domains are named after the three conserved residues tryptophan (W), tryptophan (W) and glutamate (E) encoded in this PBD, which function in stabilizing the WWE domain [71]. To date, 12 human proteins have been identified to encode a WWE domain (**Table S2).** The RNF146 WWE domain is the most studied of the WWE domains, while little is known about the other 11 WWE domain containing proteins. Structural resemblance is a characteristic that is shared by many members of the WWE domain family [72], in addition to affinity for iso-ADP-ribose, which is the smallest internal structural unit within PAR chains that contains the unique glycosidic bond [48, 71]. Structural and mutagenesis studies have demonstrated that four residues are essential for iso-ADP-ribose binding in the RNF146 WWE domain, which corresponds to Tyr107, Tyr144, Gln153, and Arg163 [71]. Despite the lack of sequence homology between the WWE domains (**Figure 1A**), these iso-ADP-ribose binding residues are conserved in most WWE domains including those encoded by TRIP12 (also named as TRIPC), Deltex1, Deltex2, Deltex4, HUWE1 and PARP11, which have been demonstrated to bind PAR chains *in vitro* [71].

**Figure 1.**
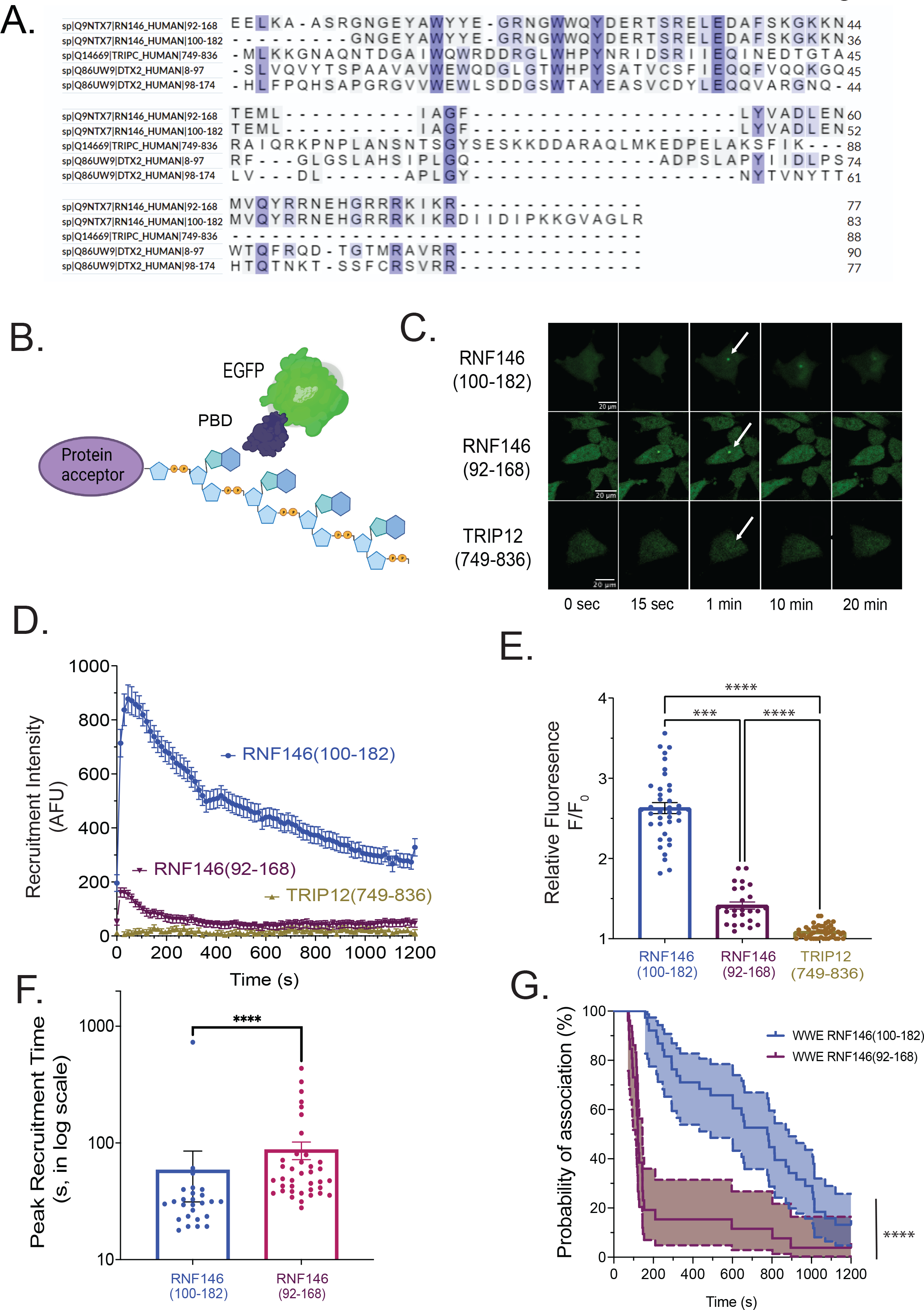
WWE domain mediated detection of PARylation levels and temporal dynamics in live cells. (**A**) Alignment of the amino acid sequences of the WWE domains from RNF146(92-168), RNF146(100-182), TRIP12(749-836), Deltex2(8-97) and Deltex2(98-174); (**B**) Graphic depicting a PAR binding domain (PBD) fused to EGFP that is used to detect levels of and temporal dynamics of PAR chains in live cells; (**C**) Confocal micrograph images of cells expressing WWE-EGFP fusions in ES-2 cells following laser micro-irradiation at multiple time points, white scale bar denotes 20µm; (**D**) Plot depicting the recruitment dynamics of WWE domains RNF146(92-168), RNF146(100-182) and TRIP12(749-836) in ES-2 cells to sites of laser micro-irradiation (405nm laser), N ≥ 28 cells, recruitment intensity normalized to nucleus fluorescent intensity background; (**E**) Relative recruitment peak intensity of the RNF146(100- 182), RNF146(92-168), and TRIP12(749-836) WWE domains expressed in ES-2 cells. Each point represents a single cell recruitment focus, graph shows mean ± SEM; (**F**) Peak recruitment time of the RNF146(100-182) and RNF146(92-168) WWE domains expressed in ES-2 cells. Each point represents a single cell recruitment focus, graph shows mean ± SEM; (**G**) Plot depicting the dissociation dynamics of the RNF146(100-182) and RNF146(92-168) WWE domains foci during 20 minutes time span following laser-induced DNA damage in ES-2 cells. N ≥ 36 cells. Exclusion percentages were 0% for RNF146(100-182) and 33.3% (12 foci) for RNF146(92-168). No exclusion for TRIP12(749-836) as it did not show any recruitment. NS: no significance, *p<0.05, **p<0.01, ***p<0.001, ****p<0.0001; a Kruskal-Wallis test was used for panel **E**, a Mann-Whitney test for panel **F** and a Kaplan-Meier test for panel **G**.

We sought to evaluate the PAR binding kinetics of some of these WWE domains (**Table 1**) in live cells following micro-irradiation-induced DNA damage. We used a fusion of each of the WWE domains encoding a C-terminal EGFP moiety (**Figure 1B**), expressed in target cells to track PAR formation at sites of DNA damage, using a laser-induced DNA damage analysis methodology, as we have described [55, 56]. LivePAR is a molecular tool for tracking PAR formation and degradation in live cells that we have previously reported [55], which is composed of a fragment of RNF146 that includes the WWE domain, corresponding to amino acid residues 100-182. This will be referred to as RNF146(100-182) from now on unless otherwise stated, as compared to the Uniprot-defined WWE domain encoded in RNF146, corresponding to amino acid residues 92-168 [73]. We then evaluated and compared the recruitment dynamics of several WWE domains, including RNF146(100-182), RNF146(92- 168), TRIP12(749-836), DELTEX2(8-97), DELTEX2(98-174), and DELTEX2(8-174), the latter encoding tandem WWE domains. In the initial screen, using a 405nm laser w/o a photosensitizer, RNF146(92-168), TRIP12(749-836), DELTEX2(8-97), DELTEX2(98-174), and DELTEX2(8-174) did not show any recruitment to the PAR formed at sites of laser micro- irradiation (**Figure S1A**). To enhance the analysis, we then used BrdU pretreatment to photosensitize cells prior to laser micro-irradiation [74] that significantly increases the level of PAR and will allow better recruitment kinetic comparisons, especially for weak binders [55]. Conversely, BrdU photosensitization promoted faint recruitment for RNF146(92-168) as compared to the strong signal for RNF146(100-182) (**Figures 1C,D**). However, we did not observe any recruitment to sites of laser-induced DNA damage for TRIP12(749-836) (**Figures 1C,D**) nor for DELTEX2(8-97), DELTEX2(98-174), or DELTEX2(7-174) (**Figure S1B**).

**Table 1:**
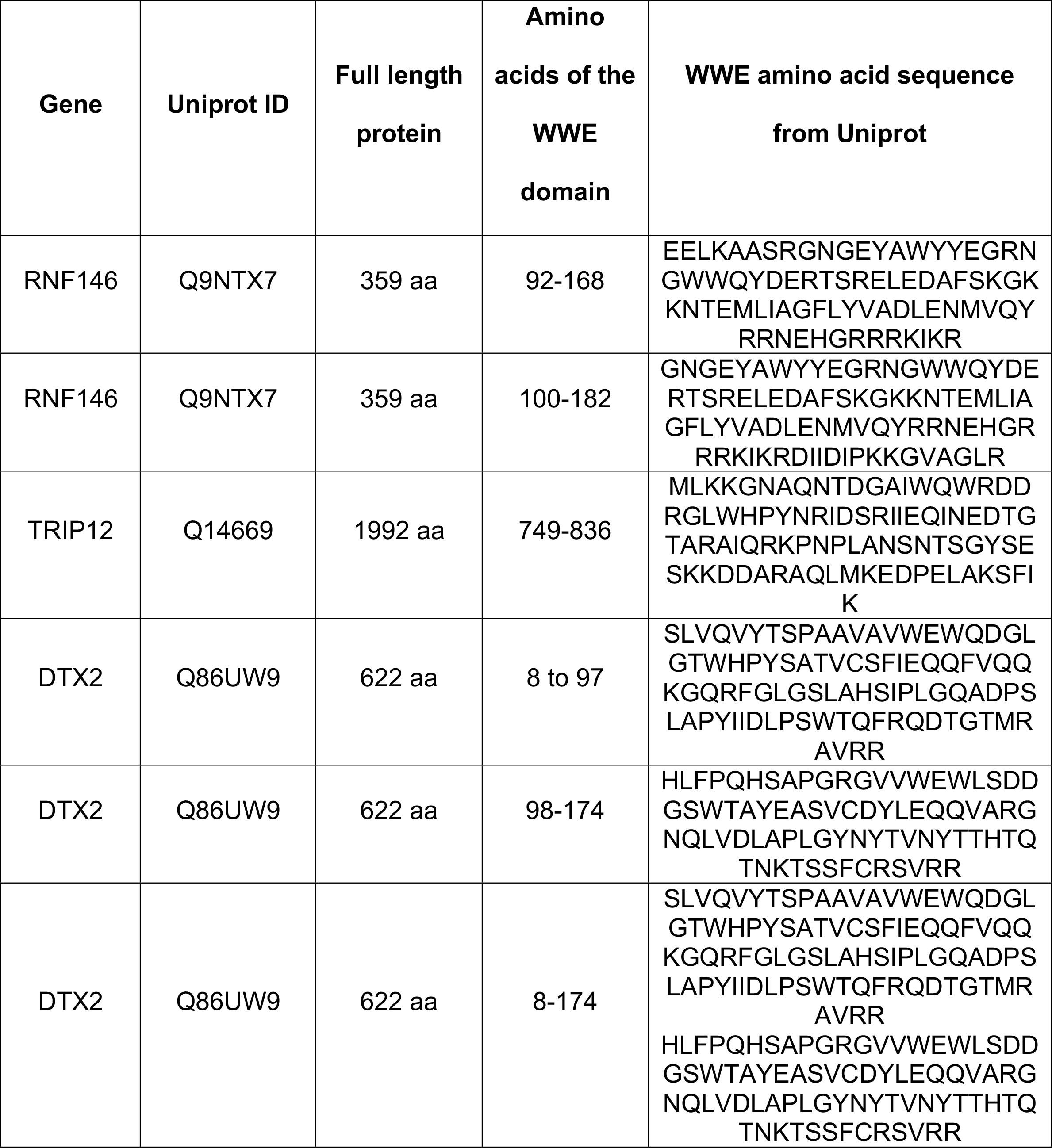
Amino acid residues encoded by the WWE domains evaluated in this study.

Statistically, the relative recruitment intensity is significantly lower for RNF146(92-168) and TRIP12(749-836), as compared to RNF146(100-182) (**Figure 1E**). In terms of peak recruitment time, RNF146(92-168) was significantly suppressed in reaching the peak level of recruitment (**Figure 1F**) but was significantly faster in its dissociation from PAR at sites of laser induced DNA damage (**Figure 1G**), as compared to RNF146(100-182). All together, these data suggest that the surrounding amino acid residues of the RNF146-encoded WWE domain affects the level of recruitment and the binding kinetics to sites of DNA damage following laser micro- irradiation, as we showed for the BRCT domain from XRCC1 [55]. This may explain the lack of recruitment seen for TRIP12(749-836), DELTEX2(8-97), DELTEX2(98-174), or DELTEX2(7-174), as these EGFP fusions proteins encode only the specific WWE domain without the surrounding amino acid residues of the parent protein.

### 3.2 Both RNF146(100-182) and Af1521(K35E/Y145R) are effective tools for live-cell tracking of PARylation dynamics

Our previous investigation of several PAR binding domains, conducted to find a molecular probe for live-cell PAR analysis, showed that the best performance was from a fragment of the RNF146 protein that includes the WWE domain corresponding to amino acid residues 100- 182: RNF146(100-182) [55]. Different PAR binding domains recognize different parts of the PAR chain. The WWE domain identifies the iso-ADP moiety of the PAR chain whereas the macrodomain binds to the terminal residue of the PAR chain [44] (**Figure 2A**). Previously, we found that the macrodomain from H2A1.1 demonstrated faint recruitment but only when cells were photosensitized with BrdU [55]. In this new study, we tested two related macrodomains, the Af1521 macrodomain from the bacterium *Archaeoglobus fulgidus*, Af1521(WT) [75], and an engineered version of Af1521 with two critical amino acid changes, Af1521(K35E/Y145R), reported to significantly increase its binding affinity to ADP-ribose, when evaluated using *in vitro* ADP-ribose binding assays [50]. To investigate the ability of the engineered macrodomain Af1521(K35E/Y145R) to detect PAR levels and PAR dynamics at DNA damage sites following laser micro-irradiation in live cells, the engineered and the wild type Af1521 macrodomains were cloned as a fusion with EGFP, expressed in cells, and imaged to ensure expression and distribution throughout the nucleus and the cytoplasm of U2OS cells (**Figure 2B**) and ES-2 cells (**Figures S2A, S2B**).

**Figure 2.**
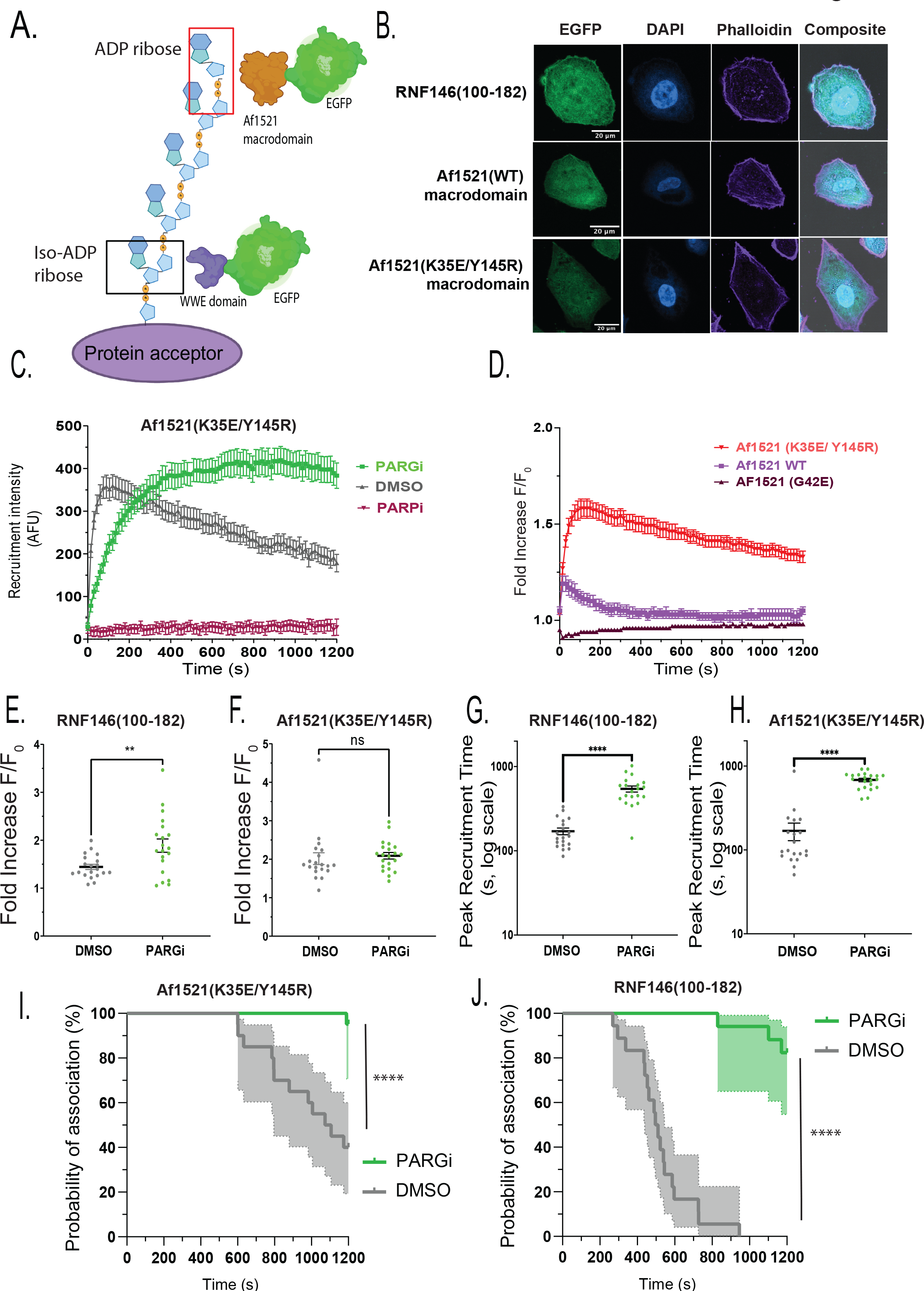
WWE domain of RNF146(100-182) and the engineered Af1521(K35E/Y145R) macrodomain as molecular probes to track PARylation dynamics in live cells. **(A)** Schematic representation of the RNF146(100-182) WWE domain (fused to EGFP) or of the Af1521 macrodomain (fused to EGFP) bound to PAR chains at sites of DNA damage; (**B**) Confocal micrograph images of EGFP-fused RNF146(100-182), Af1521(WT) or Af1521(K35E/Y145R) expressed in U2OS cells, white scale bar denotes 20µm; (**C**) Recruitment of Af1521(K35E/Y145R)-EGFP in U2OS cells following pre-treatment with vehicle (DMSO, 0.1%, 60 minutes), the PARPi ABT-888 (10μM, 60 minutes) or the PARGi PDD00017273(10μM, 60 minutes), to sites of laser micro-irradiation (405nm laser), following BrdU sensitization (10μM, 24 hours), N≥20 cells, recruitment intensity normalized to nucleus fluorescent intensity background; (**D**) Recruitment of Af1521(WT), Af1521(K35E/Y145R) and Af1521(G42E) expressed in U2OS cells to sites of laser micro-irradiation (405nm) following BrdU sensitization (10μM, 24 hours), N≥25 cells, recruitment intensity normalized to first frame; (**E**) Relative peak intensity for recruitment of Af1521(K35E/Y145R)-EGFP expressed in U2OS cells following pre- treatment with vehicle (DMSO, 0.1%, 60 minutes) or the PARG_i_ PDD00017273 (10μM, 30 minutes). Each point represents a single cell recruitment focus, graph shows mean ± SEM; (**F**) Relative peak intensity for recruitment of RNF146(100-182)-EGFP expressed in LN428 cells following pre-treatment with vehicle (DMSO, 0.1%, 60 minutes) or the PARG_i_ PDD00017273 (10μM, 30 minutes). Each point represents a single cell recruitment focus, graph shows mean ± SEM; (**G**) Peak recruitment time of Af1521(K35E/Y145R)-EGFP foci, in U2OS cells Each point represents a single cell recruitment focus, graph shows mean ± SEM; (**H**) Peak recruitment time of RNF146(100-182)-EGFP foci in LN428 cells. Each point represents a single cell recruitment focus, graph shows mean ± SEM; (**I**) Plot depicting the dissociation dynamics of Af1521(K35E/Y145R) foci during 20 minutes time span following laser-induced DNA damage in U2OS cells. N ≥ 20 cells. ; (**J**) Plot depicting the dissociation dynamics of RNF146(100-182) foci in LN428 cells during 20 minutes time span following laser-induced DNA damage in LN428 cells. N ≥ 20 cells. Recruitment foci having a relative peak intensity below 1.15 of first frame were excluded from the experiment and from statistical analysis in graphs (**E-J**). Exclusion percentages were 10% (2 foci) for DMSO treated RNF146(100-182) and 15% for (3 foci) PARGi treated RNF146(100-182). No exclusion was done in DMSO and PARGi- treated Af1521(K35E/Y145R). NS: no significance, *p<0.05, **p<0.01, ***p<0.001, ****p<0.0001; a t- test was used for panel E, a Mann-Whitney test was used for Panels **F**, **G**, and **H** and a Kaplan- Meier test for panels **I** and **J**.

The specificity of RNF146(100-182) to detect PAR in live cells was investigated previously [55, 66] by demonstrating that pretreatment of cells expressing RNF146(100-182), with a PARP1 inhibitor (PARPi; ABT-888) [76], prevented the recruitment of RNF146(100-182) by inhibiting PAR formation [55]. Conversely, pretreatment with a PARG inhibitor (PARGi; PDD00017273) [77] prevented the dissociation of RNF146(100-182) foci by inhibiting PAR degradation [55]. To demonstrate the specificity of recruitment for Af1521(K35E/Y145R) to sites of PAR formation, we showed that pretreatment of cells with a PARPi (ABT-888) inhibited recruitment whereas pretreatment with a PARGi (PDD00017273) prevented the dissociation (**Figure 2C**), like what we found for RNF146(100-182) [55].

To further evaluate the ability of the engineered macrodomain Af1521(K35E/Y145R) to detect PAR at sites of laser micro-irradiation induced DNA damage, we compared recruitment of Af1521(WT), Af1521(K35E/Y145R), and a nonbinding mutant Af1521(G42E) [50], following micro-irradiation in live cells. The initial screen, using a 405nm laser w/o a photosensitizer, showed barely detectable recruitment for Af1521(K35E/Y145R) and no recruitment for Af1521(WT) or the nonbinder mutant, Af1521(G42E) (**Figure S2C**). However, when cells were photosensitized by pre-treatment with BrdU (10μM, 24 hrs), we observed increased recruitment of Af1521(K35E/Y145R) as compared to Af1521(WT), whereas the G42E mutation, Af1521(G42E), eliminated the recruitment of Af1521 to PAR at sites of DNA damage (**Figure 2D**).

Tracking and quantitation of recruitment kinetics provides temporal distinction between the ability of different PAR binding domains to detect PAR and to reflect PAR dynamics in live cells. RNF146(100-182) had a significantly higher peak recruitment intensity when cells were pretreated with a PARGi (PDD00017273) (**Figure 2E**) but was not seen with Af1521(K35E/Y145R) (**Figure 2F**). As expected, PARGi pretreatment increased the peak recruitment time for both RNF146(100-182) (**Figure 2G**) and Af1521(K35E/Y145R) (**Figure 2H**) and decreased the dissociation of both RNF146(100-182) and Af1521(K35E/Y145R) foci at sites of DNA damage (**Figures 2I,J**). Collectively, these data show that both RNF146(100- 182) and Af1521(K35E/Y145R) are specific molecular probes with distinct capacities for detection of PAR levels and PAR dynamics at sites of micro-irradiation induced DNA damage.

### 3.3 Greater recruitment of RNF146(100-182) to sites of DNA damage following micro- irradiation as compared to Af1521(K35E/Y145R)

Despite the well-documented difference in the capacity and specificity of PAR binding for WWE domains and macrodomains [44], the context of detecting PARylation in live cells may include many variables that affect the binding kinetics of different PAR binding domains. To explore this further, we compared the recruitment kinetics of RNF146(100-182) and Af1521(K35E/Y145R) to PAR chains at sites of DNA damage following laser micro-irradiation. Our preliminary analysis showed that RNF146(100-182) recruitment was significantly higher than Af1521(K35E/Y145R) whereas we did not see any recruitment for Af1521(WT) (**Figure S2C**). However, BrdU photosensitization, prior to laser micro-irradiation, revealed minimal recruitment for Af1521(WT) with distinct recruitment kinetics as compared to Af1521(K35E/Y145R) and RNF146(100-182) (**Figures 3A,B**).

**Figure 3.**
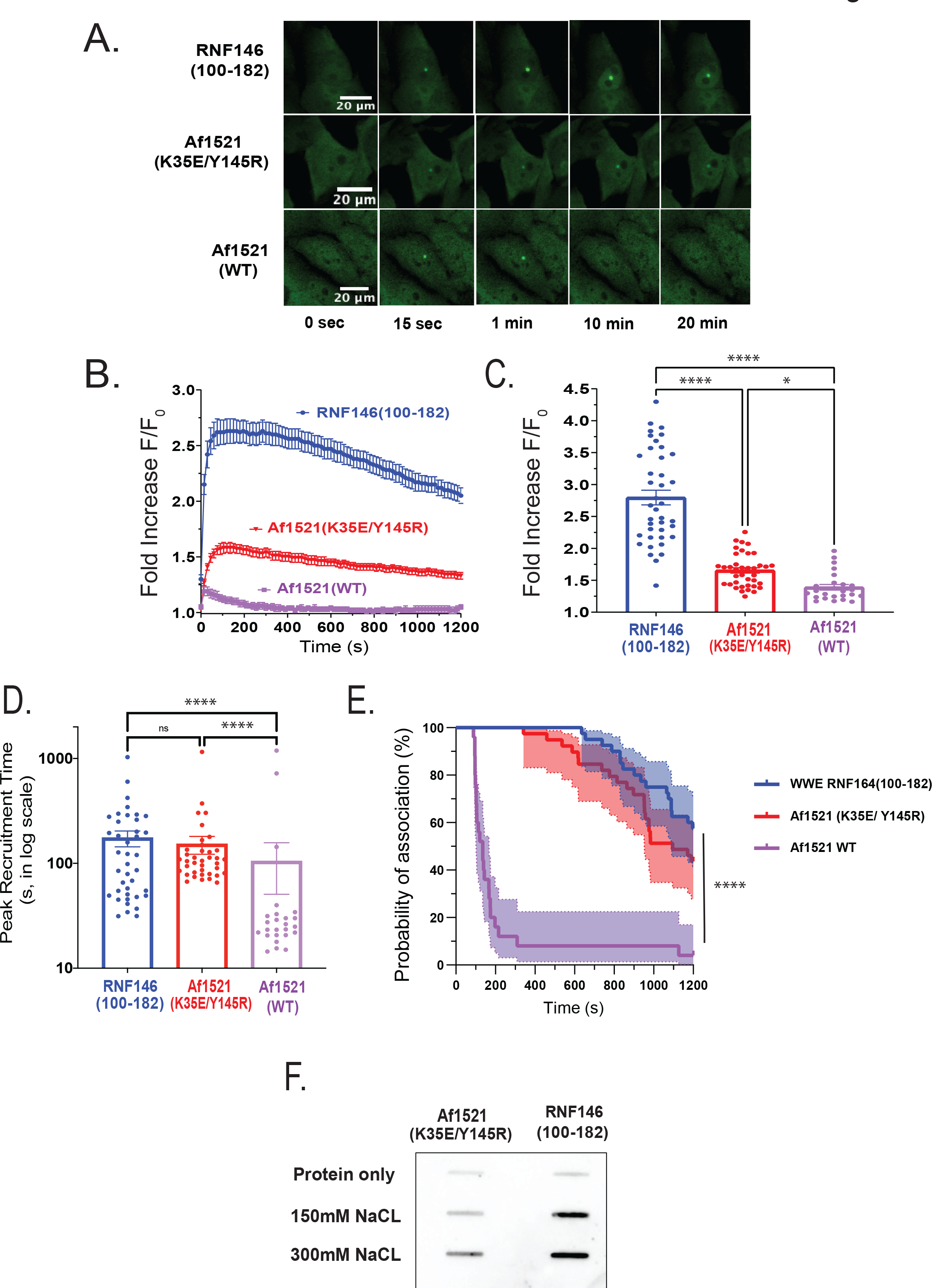
Recruitment dynamics of RNF146(100-182)-EGFP and of Af1521(K35E/Y145R)- EGFP to sites of laser micro-irradiation. (**A**) Confocal micrograph images of cells expressing EGFP fusions with RNF146(100-182), Af1521(K35E/Y145R), and Af1521(WT) in U2OS cells, following laser micro-irradiation; (**B**) Recruitment of RNF146(100-182), Af1521(WT), and Af1521(K35E/Y145R), to sites of laser micro-irradiation (405nm) following BrdU sensitization (10μM, 24 hours), N≥25 cells, recruitment intensity normalized to first frame (F/F_0_: Maximum Fluorescence intensity / Fluorescence intensity at t_0_); (**C**) Relative peak intensity of recruitment for RNF146(100-182), Af1521(WT), and Af1521(K35E/Y145R), in U2OS cells. Each point represents a single cell recruitment focus, graph shows mean ± SEM. (F/F_0_: Maximum Fluorescence intensity / Fluorescence intensity at t_0_); (**D**) Peak recruitment time for RNF146(100-182), Af1521(WT), and Af1521(K35E/Y145R) expressed in U2OS cells. Each point represents a single cell recruitment focus, graph shows mean ± SEM; (**E**) Plot depicting the dissociation dynamics of RNF146(100-182), Af1521(K35E/Y145R) and Af1521(WT) foci in U2OS cells during 20 minutes following laser-induced DNA damage, N ≥ 40 cells. Recruitment foci having a relative peak intensity below 1.15 of the first frame were excluded from the experiment and from statistical analysis in graphs (**C-E**). Exclusion percentages were 5% (2 foci) for Af1521(K35E/Y145R) and 37.5% (15 foci) for Af1521(WT). No exclusion was made in RNF146(100-182). (**F**) Immunoblots of pull-down experiments using purified GST-tagged Af1521(K35E/Y145R) (100µg) or GST-tagged RNF146(100-182) (100µg), bound to glutathione agarose beads of PAR-containing ES-2 cell lysate under increasing salt concentration, as indicated on the left side of the blot (4°C, overnight). Eluates were transferred to nitrocellulose membranes and probed by PAR primary antibody . Graph shows mean ± SEM. NS: no significance, *p<0.05, **p<0.01, ***p<0.001, ****p<0.0001; a Kruskal-Wallis test was used for panels **C** and **D** and a Kaplan-Meier test for panel **E**.

Statistical analysis of recruitment kinetics indicated that the relative peak intensity of RNF146(100-182) recruitment was significantly higher as compared to recruitment of Af1521(K35E/Y145R) or Af1521(WT) (**Figures 3C, S2D**). In terms of peak recruitment time, there was no significant difference between RNF146(100-182) or Af1521(K35E/Y145R), whereas Af1521(WT) reached its peak recruitment, after BrdU sensitization, in significantly less time as compared to both RNF146(100-182) and Af1521(K35E/Y145R) (**Figures 3D, S2E**). In addition, there was a significant difference in the dissociation of RNF146(100-182), Af1521(K35E/Y145R) and Af1521(WT) foci at sites of laser-induced DNA damage; after BrdU sensitization(**Figures 3E, S2F**).

Despite the rapid and extensive level of PARylation at sites of DNA damage, degradation of PAR is tightly controlled and occurs with a half-life of a few minutes [32, 33]. Several PAR degrading proteins have been identified, including ARH3 [78], TARG [79] and PARG [34, 35]. PARG was reported to be the most potent of the PAR hydrolases in the context of the DDR, by facilitating the resolution of the PAR signal [34, 35, 43], thereby complicating the study of protein binding to PAR in live cells. To define the binding affinity of RNF146(100- 182) and Af1521(K35E/Y145R) to PAR in the absence of the dynamic nature of PARylation in live cells, GST-tagged RNF146(100-182) and GST-tagged Af1521(K35E/Y145R) were expressed in *E. coli* and purified as GST tagged proteins (**Figures S3A, S3B**). The ability of the purified GST-fusion proteins to bind PAR was then compared using GST pull-down of PAR (**Figure 3F**), an *in vitro* PAR binding assay (**Figures S3C, S3D**) and a nitrocellulose PAR binding assay (**Figures S3E-M**). We found that the binding affinity of purified RNF146(100- 182) to PARylated proteins isolated from ES-2 cells or to purified PAR (100nM) was higher than that of the macrodomain proteins Af1521(K35E/Y145R) or Af1521(WT). All these data together demonstrate that RNF146(100-182) has a higher binding affinity to sites of PARylation following laser-induced DNA damage in live cells and to PARylated proteins in cell lysates, as compared to both macrodomain containing proteins studied herein.

*3.4 RNF146(100-182) overexpression modulates PAR levels and PAR dynamics by stabilizing PAR chains following laser micro-irradiation-induced DNA damage*

PAR chains show variable length and branching patterns [80], which can influence the number of PAR binding domains that can bind to each PAR chain. Recent findings indicate that linear PAR chains may extend to as much as 200 ADP-ribose units [21]. Conceptually, the higher binding capacity of RNF146(100-182) to PAR chains in live cells following micro-irradiation induced DNA damage and to PARylated proteins *in vitro* can be explained by 2 possible scenarios. In the first scenario, depending on the variability of the length and branching pattern of PARylation at sites of micro-irradiation induced DNA damage, WWE domain type proteins such as RNF146(100-182) may have more available binding sites on such PAR chains than macrodomain type proteins such as Af1521(K35E/Y145R). In the second scenario, RNF146(100-182) may be interfering with the dynamics of PARylation at sites of DNA damage. To delineate the latter option and to determine the ability of RNF146(100-182) and Af1521(K35E/Y145R) to interfere with PARylation dynamics, we expressed both RNF146(100- 182) and Af1521(K35E/Y145R) in U2OS and ES-2 cells, where one is fused to EGFP to allow visualization and analysis of PAR dynamics and the other is fused to a myc-tag for expression validation.

To investigate the ability of RNF146(100-184) to affect PARylation dynamics at the site of DNA damage, we compared PAR formation dynamics using the EGFP-tagged Af1521(K35E/Y145R) protein, as the visual probe, in the presence or absence of myc-tagged RNF146(100-182) (**Figure 4A**), after verifying the expression of both domains in the target cells (**Figure S4A**). Overexpression of RNF146(100-182)-myc resulted in a noticeable change in the dynamics of PAR formation and degradation at sites of DNA damage, demonstrated by higher and more prolonged recruitment of AF1521(K35E/Y145R)-EGFP (**Figures 4B, S4B**).

**Figure 4.**
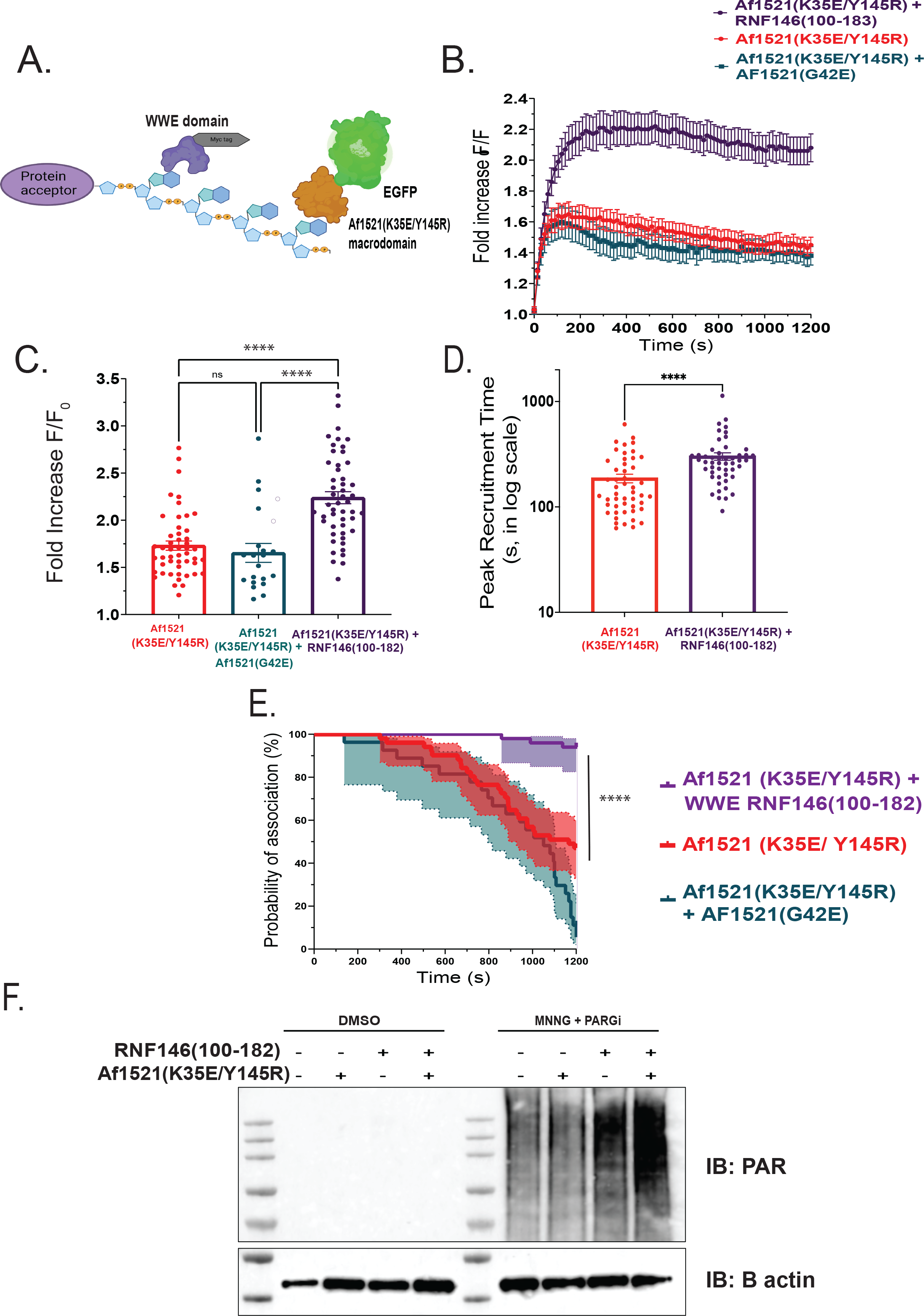
Overexpression of the RNF146(100-182) WWE domain modulates PAR levels and dynamics at sites of laser micro-irradiation. (**A**) Schematic representation of the RNF146(100- 182) WWE domain (fused to a myc-tag) and the engineered macrodomain Af1521(K35E/Y145R) (fused to EGFP). Af1521(K35E/Y145R)-EGFP is used for tracking PAR levels, and the modulation of PAR dynamics, impacted by the expression of the RNF146(100- 182)-myc; (**B**) Recruitment of Af1521(K35E/Y145R)-EGFP in U2OS cells, after overexpression of RNF146(100-182)-myc, to sites of laser micro-irradiation (405nm) following BrdU sensitization (10μM, 24 hours), N≥49 cells, recruitment intensity normalized to first frame, (F/F_0_: Maximum Fluorescence intensity / Fluorescence intensity at t_0_); (**C**) Relative peak intensity of recruitment for Af1521(K35E/Y145R)-EGFP, after overexpression of RNF146(100-182)-myc, in U2OS cells. Each point represents a single cell recruitment focus. Graph shows mean ± SEM (F/F_0_: Maximum Fluorescence intensity/ Fluorescence intensity at t_0_); (**D**) Peak recruitment time for Af1521(K35E/Y145R)-EGFP, after overexpression of RNF146(100-182)- myc, in U2OS cells. Each point represents a single cell recruitment focus, graph shows mean ± SEM; (**E**) Plot depicting the dissociation dynamics of Af1521(K35E/Y145R) foci after overexpression of RNF146(100-182) in U2OS cells during 20 minutes following laser-induced DNA damage. N ≥ 48 cells. Af1521(K35E/Y145R) EGFP foci having a relative peak intensity below 1.15 of first frame were excluded from the experiment and from statistical analysis in graphs (**C-E**). Exclusion percentages were 4% (2 foci) for Af1521(K35E/Y145R) and no exclusion for Af1521(K35E/Y145R) foci after overexpression of RNF146(100-182). (**F**) Immunoblot probing PAR levels in cells expressing Af1521(K35E/Y145R) and/or the RNF146(100-182) and after treatment with the PARG_i_ PDD00017273 (10μM, 8 hours) and MNNG (20μM, 1 hour); graph shows mean ± SEM. Af1521(K35E/Y145R)-EGFP foci having a relative peak intensity below 1.15 of first frame were excluded from the experiment and from statistical analysis in graphs (C-E). NS: no significance, *p<0.05, **p<0.01, ***p<0.001, ****p<0.0001; a Kruskal-Wallis test was used for panel **C**, a Mann-Whitney test for panel **D** and a Kaplan-Meier test for panel **E**.

Recruitment kinetic analysis revealed that the relative peak intensity of Af1521(K35E/Y145R)-EGFP was significantly increased in the presence of RNF146(100-182)- myc (**Figures 4C, S4C**). Overexpression of RNF146(100-182)-myc significantly increased the peak recruitment time of Af1521(K35E/Y145R)-EGFP (**Figures 4D, S4D**) and significantly decreased the dissociation of Af1521(K35E/Y145R)-EGFP foci in U2OS cells (**Figure 4E**) and in ES-2 cells (**Figure S4E**).

To investigate whether expressing RNF146(100-182)-myc affects PAR levels when DNA damage is induced by treatment with MNNG (10μM, 30 min), we probed PAR levels by immunoblot from RNF146(100-182)-myc expressing U2OS cells. Here, we found an increased level of PAR in cells expressing RNF146(100-182), as compared to wild type cells, or to cells expressing Af1521(K35E/Y145R) (**Figure 4F**), suggesting that over-expression of RNF146(100-182) may stabilize PAR chains.

We then investigated whether Af1521(K35E/Y145R) similarly influences PARylation dynamics at sites of DNA damage (following micro-irradiation) by tracking PAR formation and degradation by expressing RNF146(100-182)-EGFP in cells following overexpression of Af1521(K35E/Y145R)-myc (**Figures 5A, S5A**). However, overexpression of Af1521(K35E/Y145R)-myc did not promote any change in PAR levels or PAR dynamics at sites of DNA damage, as shown by the similar recruitment profile of RNF146(100-182)-EGFP in the presence or absence of Af1521(K35E/Y145R)-myc (**Figures 5B, S5B**). The relative peak intensity of RNF146(100-182)-EGFP recruitment increased slightly in the presence of Af1521(K35E/Y145R)-myc in U2OS cells (**Figures 5C**) but not in ES-2 cells (**Figure S5C**). Further, overexpression of Af1521(K35E/Y145R)-myc did not affect the peak recruitment time of RNF146(100-182)-EGFP in U2OS cells (**Figure 5D**) whereas it did increase slightly in ES-2 cells (**Figure S5D**). However, the dissociation of RNF146(100-182)-EGFP foci did not change significantly in the presence of Af1521(K35E/Y145R)-myc (**Figures 5E, S5E**). All together these data suggest that expression of the WWE domain encoded by RNF146(100-182) uniquely modulates PAR dynamics and stabilizes PAR chains at sites of laser-induced DNA damage, a phenotype not observed with expression of the engineered macrodomain encoded by Af1521(K35E/Y145R).

**Figure 5.**
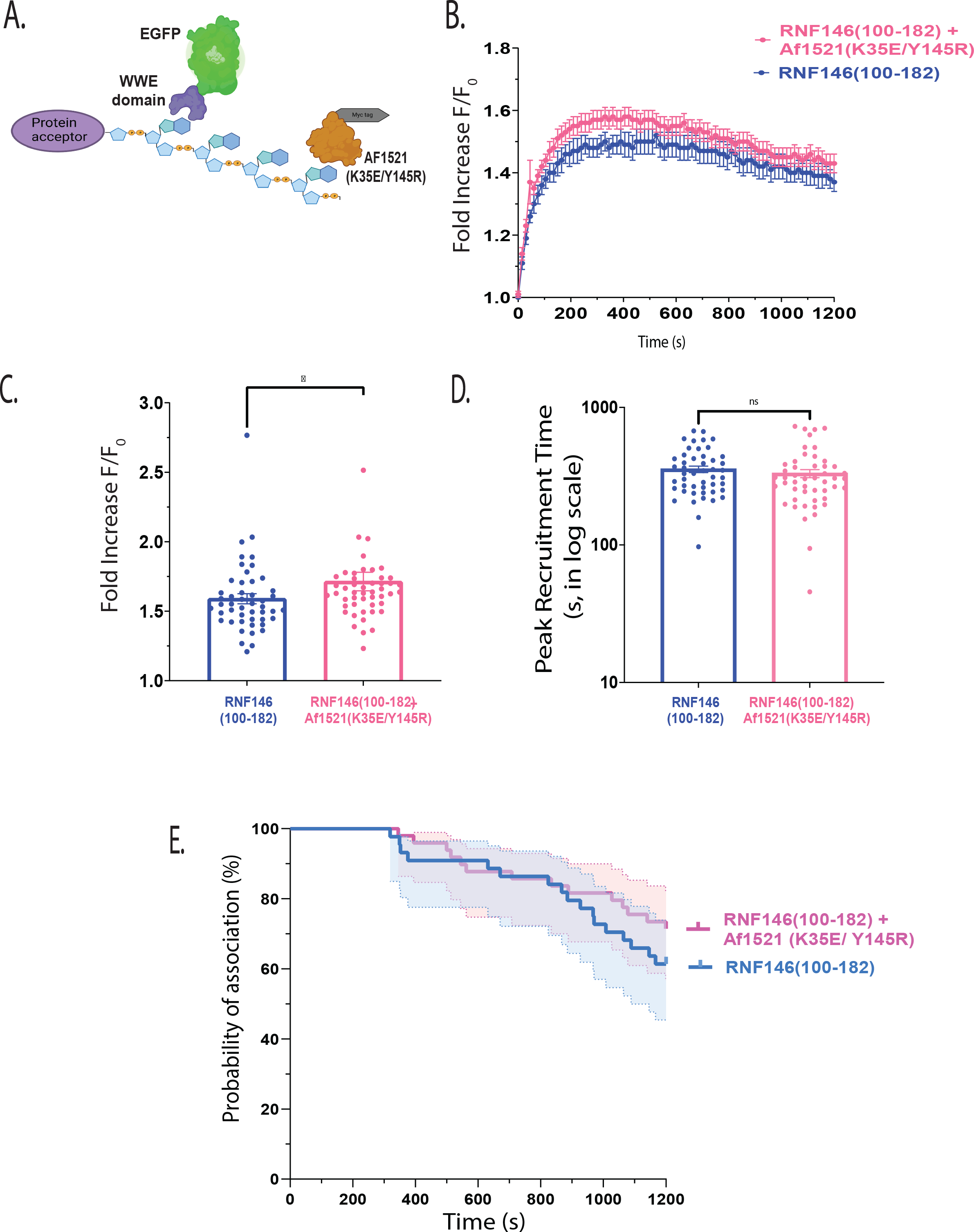
Overexpression of the engineered macrodomain Af1521(K35E/Y145R) does not modulate PAR levels or dynamics at sites of laser micro-irradiation. (**A**) Schematic representation of the engineered macrodomain Af1521(K35E/Y145R) (fused to a myc-tag) and the RNF146(100-182) WWE domain (fused to EGFP). RNF146(100-182)-EGFP is used for tracking PAR levels, and the modulation of PAR dynamics, impacted by the expression of Af1521(K35E/Y145R)-myc; (**B**) Recruitment of RNF146(100-182)-EGFP in U2OS cells, after overexpression of Af1521(K35E/Y145R)-myc, to sites of laser micro-irradiation (405nm), N≥48 cells, recruitment intensity normalized to first frame, (F/F_0_: Maximum Fluorescence intensity / Fluorescence intensity at t_0_); (**C**) Relative peak intensity of recruitment for RNF146(100-182)- EGFP, after overexpression of Af1521(K35E/Y145R)-myc, in U2OS cells at sites of laser micro- irradiation (405nm). Each point represents a single cell recruitment focus, graph shows mean ± SEM (F/F_0_: Maximum Fluorescence intensity / Fluorescence intensity at t_0_); (**D**) Peak recruitment time RNF146(100-182)-EGFP, after overexpression of Af1521(K35E/Y145R)-myc, in U2OS cells at sites of laser micro-irradiation (405nm). Each point represents a single cell recruitment focus, graph shows mean ± SEM; (**E**) Plot depicting the dissociation dynamics of RNF146(100-182) foci after overexpression of Af1521(K35E/Y145R) in U2OS cells during 20 minutes following laser-induced DNA damage; N ≥ 48 cells. RNF146(100-182)-EGFP foci having a relative peak intensity below 1.15 of first frame were excluded from the experiment and from statistical analysis in graphs (**C-E**). Exclusion percentages were 4% (2 foci) for RNF146(100-182) and 2% (1 foci) for RNF146(100-182) foci after overexpression of Af1521(K35E/Y145R). NS: no significance, *p<0.05, **p<0.01, ***p<0.001, ****p<0.0001; a Mann-Whitney test was used for panels **C** and **D** and a Kaplan-Meier test for panel **E**.

### 3.5 WWE domain binding to PAR at sites of laser micro-irradiation is not influenced by overexpression of RNF146(100-182)

It is well documented that PAR is a heterogenic structure with variable length and branching patterns [80]. As discussed above, the length of the PAR chain and the branching pattern can alter the extent of PAR binding proteins (the PAR readers) that can be recruited. In essence, this suggests that the PAR chain may be saturable and there may be direct competition between PAR binding domains that recognize similar binding sites. This may in-turn influence binding of PBD-encoded proteins. To explore this possibility, we overexpressed RNF146(100- 182)-myc and RNF146(100-182)-EGFP (**Figure 6A**) in both U2OS and ES-2 cells (**Figure S6A**). RNF146(100-182) fusion proteins should compete for the same binding sites on PAR chains in our micro-irradiation-induced DNA damage model. Interestingly, we found that the overexpression of RNF146(100-182)-myc did not affect PAR levels or PAR dynamics at sites of DNA damage, as tracked by recruitment of RNF146(100-182)-EGFP (**Figures 6B, S6B**). The relative peak intensity of RNF146(100-182)-EGFP recruitment did not change significantly in the presence of RNF146(100-182)-myc (**Figures 6C, S6C**). However, the competition of both WWE domains was revealed in the analysis of peak recruitment time, where the overexpression of RNF146(100-182)-myc significantly increased the peak recruitment time of the competing RNF146(100-182)-EGFP (**Figures 6D, S6D**). The dissociation of RNF146(100- 182)-EGFP foci significantly decreased in the presence of RNF146(100-182)-myc in U2OS cells (**Figure 6E**) but was not affected in ES-2 cells (**Figure S6E**). The similar recruitment dynamics observed for RNF146(100-182)-EGFP, in the presence of the non-fluorescent and identical PAR binding protein RNF146(100-182)-myc, suggests that PAR binding sites for RNF146(100-182)-EGFP, after the overexpression of RNF146(100-182)-myc, are readily available at sites of micro-irradiation-induced DNA damage. This conclusion also validates the results of the previous experiments in terms of possible interference of our PBDs with the endogenous PAR binding domain-encoded proteins.

**Figure 6.**
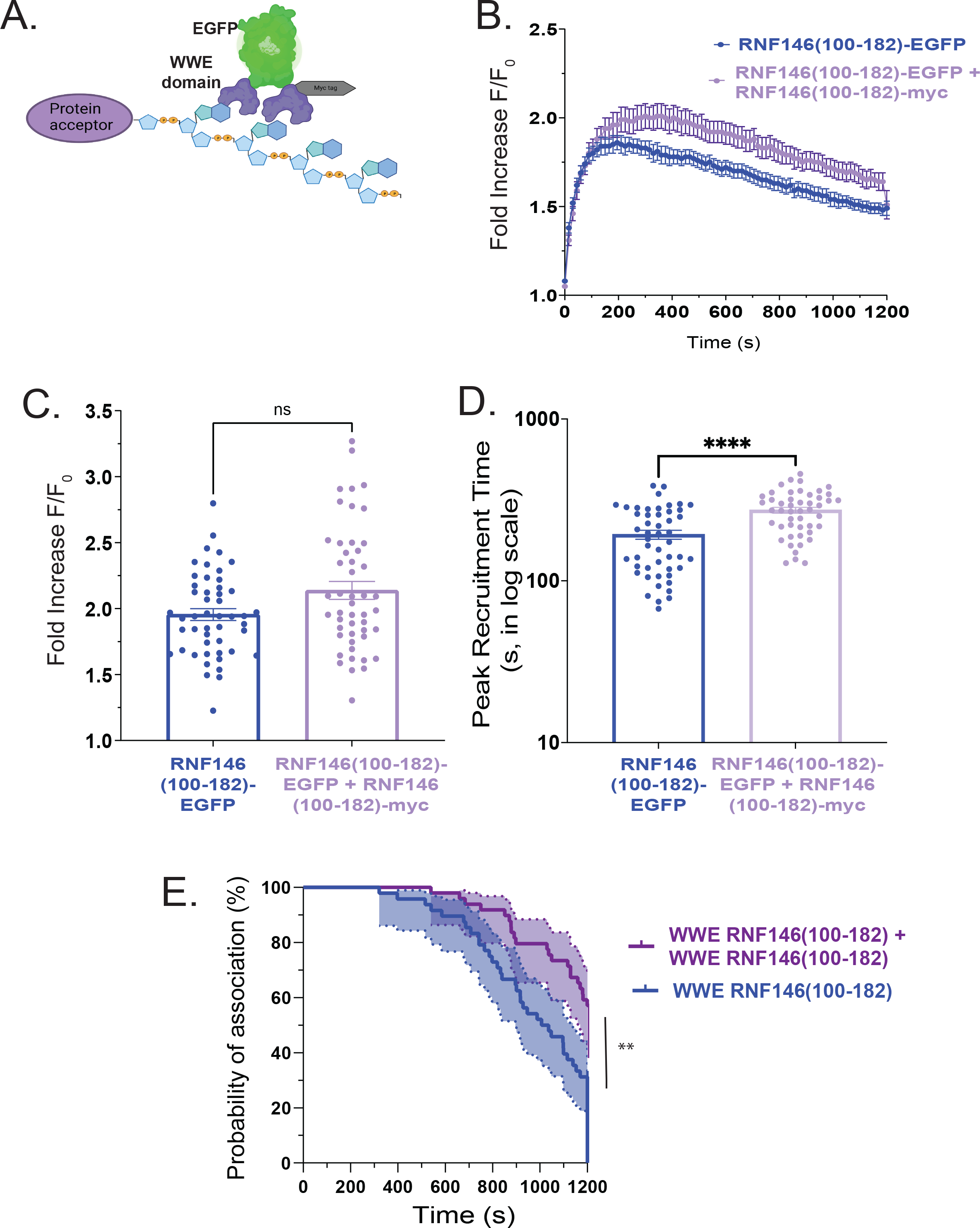
WWE domain binding to sites of laser micro-irradiation is not influenced by the overexpression of RNF146(100-182)-myc. (**A**) Schematic representation of the RNF146(100- 182) WWE domain (fused to a myc tag) and the RNF146(100-182) WWE domain (fused to EGFP). RNF146(100-182)-EGFP is used for tracking PAR levels, and the modulation of PAR dynamics, while competing for available binding sites with RNF146(100-182)-myc; (**B**) Recruitment of RNF146(100-182)-EGFP in U2OS cells, after overexpression of RNF146(100- 182)-myc, to sites of laser micro-irradiation (405nm, N≥48 cells), recruitment intensity normalized to the first frame, (F/F_0_: Maximum Fluorescence intensity / Fluorescence intensity at t_0_); (**C**) Relative peak intensity of recruitment of RNF146(100-182)-EGFP, after overexpression of RNF146(100-182)-myc, in U2OS cells at sites of laser micro-irradiation (405nm). Each point represents a single cell recruitment focus, graph shows mean ± SEM (F/F_0_: Maximum Fluorescence intensity / Fluorescence intensity at t_0_); (**D**) Peak recruitment time for RNF146(100-182)-EGFP, after overexpression of RNF146(100-182)-myc, in U2OS cells at sites of laser micro-irradiation (405nm). Each point represents a single cell recruitment focus, graph shows mean ± SEM; (**E**) Plot depicting the dissociation dynamics of RNF146(100- 182) foci after overexpression of RNF146(100-182)-myc in U2OS cells during 20 minutes following laser-induced DNA damage, N ≥ 48 cells. RNF146(100-182)-EGFP foci having a relative peak intensity below 1.15 of first frame were excluded from the experiment and from statistical analysis in graphs (C-E). Exclusion percentages were 2% (1 foci) for RNF146(100- 182) and 4% (2 foci) for RNF146(100-182) foci after overexpression of RNF146(100-182)-myc. NS: no significance, *p<0.05, **p<0.01, ***p<0.001, ****p<0.0001; a Mann-Whitney test was used for panels **C** and **D** and a Kaplan-Meier test for panel **E**.

## 4. Discussion

Poly-ADP-ribose (PAR) plays an essential role in the DNA damage response in many DNA repair pathways including BER [2, 81], by promoting the recruitment of DNA repair proteins to sites of DNA damage. PAR structure may influence biochemical and cellular outcomes in various ways. The heterogeneity of PAR chains in terms of length and branching pattern is gaining increased interest [80]. Recent findings suggest that PAR structural diversity affects the binding dynamics of its readers [82, 83], which can lead to selectivity of downstream binding partners. In addition, the length of the PAR chain and its branching pattern can influence the availability of binding sites in any PAR chain, whereas the available space for PAR binding domains of reader proteins that identify linear regions of the polymer can be limited by a highly branched PAR chain [80, 83]. Whether a PARylation-mediated signal is conveyed with high specificity by PAR chains of unique structural patterns is still to be explored and thus, PAR structural heterogeneity influence and biological importance on the binding of its readers remains elusive. Hence, various biosensors that reflect the different aspects of PAR dynamics would have added benefit to our understanding of this complicated process.

PARylation plays a role in various cellular processes which is conveyed via the dynamic interactions with PAR-binding proteins. These readers have become the effectors of PAR signaling to mediate certain cellular functions. The extensive nature of PAR-protein interactions displays high specificity and regulation, which is emphasized by the variable structural characteristics of PAR, the structurally different PAR binding domains with various PAR binding characteristics in addition to the presence of these PAR binding domains in many proteins. The binding kinetics of these PBDs is still unclear despite the significant impact of these binding properties on the regulation of PARylation signaling.

PAR binding, mediated by the identification of iso-ADP-ribose, is a common function shared between the WWE domain family members, as shown by surface plasma resonance and isothermal calorimetry [71]. However, despite the conserved iso-ADP-ribose binding residues in the WWE domains of TRIP12 and DELTEX2, we found that the WWE domain of RNF146 was the only WWE domain among the WWE domains that we tested which recruited to PAR at sites of micro-irradiation induced DNA damage. In addition, we found that the RNF146 fragment RNF146(100-182), that encodes the WWE domain, demonstrated more robust PAR binding in live cells as compared to the RNF146 WWE domain defined by amino acid residues 92-168. This is in-line with a previous report whereby the C-terminal region (169- 183) of the RNF146-encoded WWE domain underwent noticeable conformational changes upon iso-ADP-ribose binding to support binding to the distal ribose-phosphate group of iso-ADP-ribose [71]. Consequently, it remains to be seen if the addition of N-terminal or C-terminal residues outside of the WWE domains of TRIP12 or DELTEX2 (or the expression of the full-length protein) may enhance recruitment, as we have seen previously with the BRCT domain of XRCC1 [55].

While studying the binding kinetics of PBDs to PAR, many variables should be taken into consideration, such as the following: (i) while *in vitro* binding kinetic studies can show the actual binding affinities between PAR and the individual PBD, *in vivo* studies are influenced by the presence of other endogenous PBD encoded proteins with a possible synergistic or competitive effect; (ii) the presence of the PBD within the context of a part of or the whole of their respective proteins, as we have reported previously [55], where the binding kinetics of the BRCT domain to PAR was dependent on the presence of the adjacent XL1 linker. In the same fashion, some proteins encode two or more similar or different PBDs - whether these multiple PBDs would affect their binding kinetics to PAR is not yet known; and (iii) the dynamic nature of PAR *in vivo*, and whether this dynamic is influenced or regulated by other available PBD encoded proteins. All these factors indicate there is much to explore in terms of the role of the binding of PBDs (readers) to PAR in live cells, to be able to transduce the PAR signal.

Several probes have been reported to detect PAR in live cells, with differences in capacity to track PAR dynamics, such as the PBZ domain and the FHA domain, in addition to the WWE domain [55, 56, 58, 59, 84, 85]. In this study, we are reporting the engineered macrodomain Af1521(K35E/Y145R) as an additional molecular probe that can specifically detect PAR at sites of DNA damage in cells, as shown by the failure of recruitment when a PAR binding site mutation was introduced, Af1521(G42E), or after treatment with a PARP inhibitor and with enhanced binding in the presence of a PARG inhibitor. However, the engineered macrodomain Af1521(K35E/Y145R) showed a less dynamic range as compared to the WWE domain encoded by RNF146(100-182) in live cells and less PAR binding affinity *in vitro*. This can be explained by the abundant iso-ADP-ribose sites within PAR chains (**Figure 6**) as compared to the limited number of terminal ADP-ribose units recognized by macrodomains. Although fluorescence microscopy is a valuable technique that allows one to study the temporal resolution for the assembly and disassembly of key repair proteins at DNA damage sites, some limitations for this real-time approach includes the possibility of PBD-containing proteins to impede the recruitment of endogenous PAR binding proteins and depending on the level of expression of these proteins, DNA repair could be altered in cells expressing these PBDs. In addition, in the case of low PAR formation at the sites of DNA damage, or low recruitment of some of these PBD domains, the signal could be masked by the background noise with a limitation of detecting low PAR formation or low-affinity binders.

Therefore, it is possible that the other WWE domains we have tested are recruiting weakly yet cannot be detected with this technique. In addition, the possibility that these PAR binding domains may require a structural conformation that may present within the whole protein context and cannot bind in the absence of this conformation. Evaluating the recruitment of the whole protein would be required before a final statement can be made about the ability of the other WWE domains to recognize PAR formed at sites of DNA damage.

As with other PTMs, reversibility of ADP-ribosylation is key to the dynamic regulation of this signaling event. The timely degradation of PAR is an equally essential process in the DNA damage response as it is believed to prevent trapping of repair factors contributing to the initial wave of the DNA damage response and allows downstream repair factors to access the site of DNA damage [11]. A key finding of this study is the influence of RNF146(100-182) on the degradation and dynamics of PAR, as represented by the increased half-life and retained foci at sites of DNA damage. This is in-line with a previous finding in which the overexpression of the mouse Rnf146 encoded WWE domain resulted in an increased level of cellular PAR, which was explained by the increased steady-state level of the PARylated proteins PARP5A and PARP5B [86]. Similarly, it is possible that RNF146(100-182) is stabilizing PAR at the sites of micro-irradiation and interfering with PAR turnover.

PARylation dynamics seems to be associated with cancer [87, 88]. Inhibition of PAR formation, using PARP inhibitors, demonstrated high efficiency in cancer therapy [89]. PARylation is a major component in the DDR that is tightly controlled to maintain cell homeostasis, especially in cancer cells which are known for their increased level of replication stress [88]. Identification of modulators of PAR production and degradation that can modulate PAR dynamics can open new possibilities for selective cancer treatment. Hence, development of RNF146(100-182) into a genetically encoded inhibitor of dePARylation may have benefit for emerging PARG inhibitor-resistant cancer genotypes.

Finally, another aspect worth considering, despite being outside the scope of this study, is the protein RNF146, an E3 ubiquitin ligase that is believed to be involved in the coupling of PARylation and ubiquitination pathways in the context of the DDR [90]. The robust recruitment of RNF146(100-182) to sites of micro-irradiation induced DNA damage, as compared to other PBDs that we tested in this study and in previous studies [55], supports a role for RNF146 in the DDR. Although some previous studies suggested a role of RNF146 in the DDR and protection of the cell following DNA damage [91], the specific role that RNF146 plays in the BER pathway or other DDR pathways and whether it recruits to sites of DNA damage is still unknown and remains to be explored.

## Supporting information

Supplement and Full blots

## Acknowledgements

We would like to thank Aishwarya Prakash and Marlo Thompson (University of South Alabama) for their valuable help in the layout of the protein purification scheme. We also would like to thank Jianfeng Li (Emory University) and MD Maruf Khan (Brown University) for critical feedback during experimental planning and data analysis.

## Appendix A. Supplementary Data

Supplementary material related to this article can be found, in the online version, at xxxx

## References

[1] P.O. Hassa, M.O. Hottiger, The diverse biological roles of mammalian PARPS, a small but powerful family of poly-ADP-ribose polymerases, Front Biosci, 13 (2008) 3046–3082.

[2] K.M. Saville, J. Clark, A. Wilk, G.D. Rogers, J.F. Andrews, C.A. Koczor, R.W. Sobol, NAD(+)-mediated regulation of mammalian base excision repair, DNA Repair (Amst), 93 (2020) 102930.

[3] K. Ueda, O. Hayaishi, ADP-ribosylation, Annu Rev Biochem, 54 (1985) 73–100.

[4] D. Corda, M. Di Girolamo, Functional aspects of protein mono-ADP-ribosylation, EMBO J, 22 (2003) 1953–1958.

[5] M.U. Musheev, L. Schomacher, A. Basu, D. Han, L. Krebs, C. Scholz, C. Niehrs, Mammalian N1-adenosine PARylation is a reversible DNA modification, Nature communications, 13 (2022) 6138.

[6] L. Weixler, K. Scharinger, J. Momoh, B. Luscher, K.L.H. Feijs, R. Zaja, ADP-ribosylation of RNA and DNA: from in vitro characterization to in vivo function, Nucleic Acids Res, 49 (2021) 3634–3650.

[7] L. Weixler, K.L.H. Feijs, R. Zaja, ADP-ribosylation of RNA in mammalian cells is mediated by TRPT1 and multiple PARPs, Nucleic Acids Res, 50 (2022) 9426–9441.

[8] M.O. Hottiger, P.O. Hassa, B. Luscher, H. Schuler, F. Koch-Nolte, Toward a unified nomenclature for mammalian ADP-ribosyltransferases, Trends Biochem Sci, 35 (2010) 208–219.

[9] C. Liu, A. Vyas, M.A. Kassab, A.K. Singh, X. Yu, The role of poly ADP-ribosylation in the first wave of DNA damage response, Nucleic Acids Res, 45 (2017) 8129–8141.

[10] V. Schreiber, F. Dantzer, J.C. Ame, G. de Murcia, Poly(ADP-ribose): novel functions for an old molecule, Nat Rev Mol Cell Biol, 7 (2006) 517–528.

[11] C. Liu, X. Yu, ADP-ribosyltransferases and poly ADP-ribosylation, Curr Protein Pept Sci, 16 (2015) 491–501.

[12] J.J. Bonfiglio, P. Fontana, Q. Zhang, T. Colby, I. Gibbs-Seymour, I. Atanassov, E. Bartlett, R. Zaja, I. Ahel, I. Matic, Serine ADP-Ribosylation Depends on HPF1, Mol Cell, 65 (2017) 932–940 e936.

[13] G. Jankevicius, A. Ariza, M. Ahel, I. Ahel, The Toxin-Antitoxin System DarTG Catalyzes Reversible ADP-Ribosylation of DNA, Mol Cell, 64 (2016) 1109–1116.

[14] T. Nakano, Y. Matsushima-Hibiya, M. Yamamoto, S. Enomoto, Y. Matsumoto, Y. Totsuka, M. Watanabe, T. Sugimura, K. Wakabayashi, Purification and molecular cloning of a DNA ADP-ribosylating protein, CARP-1, from the edible clam Meretrix lamarckii, Proc Natl Acad Sci U S A, 103 (2006) 13652–13657.

[15] T. Nakano, Y. Matsushima-Hibiya, M. Yamamoto, A. Takahashi-Nakaguchi, H. Fukuda, M. Ono, T. Takamura-Enya, H. Kinashi, Y. Totsuka, ADP-ribosylation of guanosine by SCO5461 protein secreted from Streptomyces coelicolor, Toxicon, 63 (2013) 55–63.

[16] T. Nakano, A. Takahashi-Nakaguchi, M. Yamamoto, M. Watanabe, Pierisins and CARP- 1: ADP-ribosylation of DNA by ARTCs in butterflies and shellfish, Curr Top Microbiol Immunol, 384 (2015) 127–149.

[17] D. Perina, A. Mikoc, J. Ahel, H. Cetkovic, R. Zaja, I. Ahel, Distribution of protein poly(ADP-ribosyl)ation systems across all domains of life, DNA Repair (Amst), 23 (2014) 4–16.

[18] A. Kratz, M. Kim, M.R. Kelly, F. Zheng, C.A. Koczor, J. Li, K. Ono, Y. Qin, C. Churas, J. Chen, R.T. Pillich, J. Park, M. Modak, R. Collier, K. Licon, D. Pratt, R.W. Sobol, N.J. Krogan, T. Ideker, A multi-scale map of protein assemblies in the DNA damage response, Cell Syst, 14 (2023) 447–463 e448.

[19] K.H. Almeida, R.W. Sobol, A unified view of base excision repair: lesion-dependent protein complexes regulated by post-translational modification, DNA Repair (Amst), 6 (2007) 695–711.

[20] L. van Beek, E. McClay, S. Patel, M. Schimpl, L. Spagnolo, T. Maia de Oliveira, PARP Power: A Structural Perspective on PARP1, PARP2, and PARP3 in DNA Damage Repair and Nucleosome Remodelling, International journal of molecular sciences, 22 (2021).

[21] S. Shall, G. de Murcia, Poly(ADP-ribose) polymerase-1: what have we learned from the deficient mouse model?, Mutation Research, 460 (2000) 1–15.

[22] C.M. Daniels, S.E. Ong, A.K. Leung, Phosphoproteomic approach to characterize protein mono- and poly(ADP-ribosyl)ation sites from cells, J Proteome Res, 13 (2014) 3510–3522.

[23] S.C. Larsen, I.A. Hendriks, D. Lyon, L.J. Jensen, M.L. Nielsen, Systems-wide Analysis of Serine ADP-Ribosylation Reveals Widespread Occurrence and Site-Specific Overlap with Phosphorylation, Cell reports, 24 (2018) 2493–2505 e2494.

[24] M. Altmeyer, S. Messner, P.O. Hassa, M. Fey, M.O. Hottiger, Molecular mechanism of poly(ADP-ribosyl)ation by PARP1 and identification of lysine residues as ADP-ribose acceptor sites, Nucleic Acids Res, 37 (2009) 3723–3738.

[25] P. Li, Y. Zhen, Y. Yu, Site-specific analysis of the Asp- and Glu-ADP-ribosylated proteome by quantitative mass spectrometry, Methods Enzymol, 626 (2019) 301–321.

[26] M. Li, L.Y. Lu, C.Y. Yang, S. Wang, X. Yu, The FHA and BRCT domains recognize ADP-ribosylation during DNA damage response, Genes Dev, 27 (2013) 1752–1768.

[27] M. Li, X. Yu, Function of BRCA1 in the DNA damage response is mediated by ADP-ribosylation, Cancer Cell, 23 (2013) 693–704.

[28] I. Ahel, D. Ahel, T. Matsusaka, A.J. Clark, J. Pines, S.J. Boulton, S.C. West, Poly(ADP-ribose)-binding zinc finger motifs in DNA repair/checkpoint proteins, Nature, 451 (2008) 81–85.

[29] G.Y. Li, R.D. McCulloch, A.L. Fenton, M. Cheung, L. Meng, M. Ikura, C.A. Koch, Structure and identification of ADP-ribose recognition motifs of APLF and role in the DNA damage response, Proc Natl Acad Sci U S A, 107 (2010) 9129–9134.

[30] L. Wei, S. Nakajima, C.L. Hsieh, S. Kanno, M. Masutani, A.S. Levine, A. Yasui, L. Lan, Damage response of XRCC1 at sites of DNA single strand breaks is regulated by phosphorylation and ubiquitylation after degradation of poly(ADP-ribose), J Cell Sci, 126 (2013) 4414–4423.

[31] D. Harrision, P. Gravells, R. Thompson, H.E. Bryant, Poly(ADP-Ribose) Glycohydrolase (PARG) vs. Poly(ADP-Ribose) Polymerase (PARP) - Function in Genome Maintenance and Relevance of Inhibitors for Anti-cancer Therapy, Front Mol Biosci, 7 (2020) 191.

[32] R. Bernardi, L. Rossi, G.G. Poirier, A.I. Scovassi, Analysis of poly(ADP-ribose) glycohydrolase activity in nuclear extracts from mammalian cells, Biochim Biophys Acta, 1338 (1997) 60–68.

[33] R. Alvarez-Gonzalez, F.R. Althaus, Poly(ADP-ribose) catabolism in mammalian cells exposed to DNA-damaging agents, Mutat Res, 218 (1989) 67–74.

[34] J.P. Gagne, M.J. Hendzel, A. Droit, G.G. Poirier, The expanding role of poly(ADP-ribose) metabolism: current challenges and new perspectives, Curr Opin Cell Biol, 18 (2006) 145–151.

[35] M.E. Bonicalzi, J.F. Haince, A. Droit, G.G. Poirier, Regulation of poly(ADP-ribose) metabolism by poly(ADP-ribose) glycohydrolase: where and when?, Cell Mol Life Sci, 62 (2005) 739–750.

[36] W. Lin, J.C. Ame, N. Aboul-Ela, E.L. Jacobson, M.K. Jacobson, Isolation and characterization of the cDNA encoding bovine poly(ADP-ribose) glycohydrolase, J Biol Chem, 272 (1997) 11895–11901.

[37] J.C. Ame, F. Apiou, E.L. Jacobson, M.K. Jacobson, Assignment of the poly(ADP-ribose) glycohydrolase gene (PARG) to human chromosome 10q11.23 and mouse chromosome 14B by in situ hybridization, Cytogenet Cell Genet, 85 (1999) 269–270.

[38] S.M. Hoang, N. Kaminski, R. Bhargava, J. Barroso-Gonzalez, M.L. Lynskey, L. Garcia-Exposito, J.L. Roncaioli, A.R. Wondisford, C.T. Wallace, S.C. Watkins, D.I. James, I.D. Waddell, D. Ogilvie, K.M. Smith, F. da Veiga Leprevost, D. Mellacharevu, A.I. Nesvizhskii, J. Li, D. Ray-Gallet, R.W. Sobol, G. Almouzni, R.J. O’Sullivan, Regulation of ALT-associated homology-directed repair by polyADP-ribosylation, Nat Struct Mol Biol, 27 (2020) 1152–1164.

[39] L. Davidovic, M. Vodenicharov, E.B. Affar, G.G. Poirier, Importance of poly(ADP-ribose) glycohydrolase in the control of poly(ADP-ribose) metabolism, Exp Cell Res, 268 (2001) 7–13.

[40] J. Li, C.A. Koczor, K.M. Saville, F. Hayat, A. Beiser, S. McClellan, M.E. Migaud, R.W. Sobol, Overcoming Temozolomide Resistance in Glioblastoma via Enhanced NAD(+) Bioavailability and Inhibition of Poly-ADP-Ribose Glycohydrolase, Cancers (Basel), 14 (2022).

[41] J. Li, M.S. K M. Ibrahim, X. Zeng, S. McClellan, A. Angajala, A. Beiser, J.F. Andrews, M. Sun, C.A. Koczor, J. Clark, F. Hayat, M.V. Makarov, A. Wilk, N.A. Yates, M.E. Migaud, R.W. Sobol, NAD(+) bioavailability mediates PARG inhibition-induced replication arrest, intra S-phase checkpoint and apoptosis in glioma stem cells, NAR Cancer, 3 (2021) zcab044.

[42] M. Ikejima, D.M. Gill, Poly(ADP-ribose) degradation by glycohydrolase starts with an endonucleolytic incision, J Biol Chem, 263 (1988) 11037–11040.

[43] D. Slade, M.S. Dunstan, E. Barkauskaite, R. Weston, P. Lafite, N. Dixon, M. Ahel, D. Leys, I. Ahel, The structure and catalytic mechanism of a poly(ADP-ribose) glycohydrolase, Nature, 477 (2011) 616–620.

[44] F. Teloni, M. Altmeyer, Readers of poly(ADP-ribose): designed to be fit for purpose, Nucleic Acids Res, 44 (2016) 993–1006.

[45] H. Wei, X. Yu, Functions of PARylation in DNA Damage Repair Pathways, Genomics Proteomics Bioinformatics, 14 (2016) 131–139.

[46] J.M. Pleschke, H.E. Kleczkowska, M. Strohm, F.R. Althaus, Poly(ADP-ribose) binds to specific domains in DNA damage checkpoint proteins, J Biol Chem, 275 (2000) 40974–40980.

[47] J.P. Gagne, M. Isabelle, K.S. Lo, S. Bourassa, M.J. Hendzel, V.L. Dawson, T.M. Dawson, G.G. Poirier, Proteome-wide identification of poly(ADP-ribose) binding proteins and poly(ADP-ribose)-associated protein complexes, Nucleic Acids Res, 36 (2008) 6959–6976.

[48] L. Aravind, The WWE domain: a common interaction module in protein ubiquitination and ADP ribosylation, Trends Biochem Sci, 26 (2001) 273–275.

[49] E. Barkauskaite, G. Jankevicius, I. Ahel, Structures and Mechanisms of Enzymes Employed in the Synthesis and Degradation of PARP-Dependent Protein ADP-Ribosylation, Mol Cell, 58 (2015) 935–946.

[50] K. Nowak, F. Rosenthal, T. Karlberg, M. Butepage, A.G. Thorsell, B. Dreier, J. Grossmann, J. Sobek, R. Imhof, B. Luscher, H. Schuler, A. Pluckthun, D.M. Leslie Pedrioli, M.O. Hottiger, Engineering Af improves ADP-ribose binding and identification of ADP-ribosylated proteins, Nature communications, 11 (2020) 5199.

[51] M. Altmeyer, K.J. Neelsen, F. Teloni, I. Pozdnyakova, S. Pellegrino, M. Grofte, M.D. Rask, W. Streicher, S. Jungmichel, M.L. Nielsen, J. Lukas, Liquid demixing of intrinsically disordered proteins is seeded by poly(ADP-ribose), Nature communications, 6 (2015) 8088.

[52] T. Karlberg, M.F. Langelier, J.M. Pascal, H. Schuler, Structural biology of the writers, readers, and erasers in mono- and poly(ADP-ribose) mediated signaling, Mol Aspects Med, 34 (2013) 1088–1108.

[53] R. Martello, M. Leutert, S. Jungmichel, V. Bilan, S.C. Larsen, C. Young, M.O. Hottiger, M.L. Nielsen, Proteome-wide identification of the endogenous ADP-ribosylome of mammalian cells and tissue, Nature communications, 7 (2016) 12917.

[54] N. Dani, A. Stilla, A. Marchegiani, A. Tamburro, S. Till, A.G. Ladurner, D. Corda, M. Di Girolamo, Combining affinity purification by ADP-ribose-binding macro domains with mass spectrometry to define the mammalian ADP-ribosyl proteome, Proc Natl Acad Sci U S A, 106 (2009) 4243–4248.

[55] C.A. Koczor, K.M. Saville, J.F. Andrews, J. Clark, Q. Fang, J. Li, R.Q. Al-Rahahleh, M. Ibrahim, S. McClellan, M.V. Makarov, M.E. Migaud, R.W. Sobol, Temporal dynamics of base excision/single-strand break repair protein complex assembly/disassembly are modulated by the PARP/NAD(+)/SIRT6 axis, Cell reports, 37 (2021) 109917.

[56] C.A. Koczor, K.M. Saville, R.Q. Al-Rahahleh, J.F. Andrews, J. Li, R.W. Sobol, Quantitative Analysis of Nuclear Poly(ADP-Ribose) Dynamics in Response to Laser-Induced DNA Damage, Methods Mol Biol, 2609 (2023) 43–59.

[57] J.L. Furman, P.W. Mok, S. Shen, C.I. Stains, I. Ghosh, A turn-on split-luciferase sensor for the direct detection of poly(ADP-ribose) as a marker for DNA repair and cell death, Chem Commun (Camb), 47 (2011) 397–399.

[58] D.B. Krastev, S.J. Pettitt, J. Campbell, F. Song, B.E. Tanos, S.S. Stoynov, A. Ashworth, C.J. Lord, Coupling bimolecular PARylation biosensors with genetic screens to identify PARylation targets, Nature communications, 9 (2018) 2016.

[59] E.O. Serebrovskaya, N.M. Podvalnaya, V.V. Dudenkova, A.S. Efremova, N.G. Gurskaya, D.A. Gorbachev, A.V. Luzhin, O.L. Kantidze, E.V. Zagaynova, S.I. Shram, K.A. Lukyanov, Genetically Encoded Fluorescent Sensor for Poly-ADP-Ribose, International journal of molecular sciences, 21 (2020).

[60] E.C. Greenwald, S. Mehta, J. Zhang, Genetically Encoded Fluorescent Biosensors Illuminate the Spatiotemporal Regulation of Signaling Networks, Chemical reviews, 118 (2018) 11707–11794.

[61] N.R. Gassman, S.H. Wilson, Micro-irradiation tools to visualize base excision repair and single-strand break repair, DNA Repair (Amst), 31 (2015) 52–63.

[62] S. Zentout, R. Smith, M. Jacquier, S. Huet, New Methodologies to Study DNA Repair Processes in Space and Time Within Living Cells, Front Cell Dev Biol, 9 (2021) 730998.

[63] H. Kawamitsu, H. Hoshino, H. Okada, M. Miwa, H. Momoi, T. Sugimura, Monoclonal antibodies to poly(adenosine diphosphate ribose) recognize different structures, Biochemistry, 23 (1984) 3771–3777.

[64] B.A. Gibson, L.B. Conrad, D. Huang, W.L. Kraus, Generation and Characterization of Recombinant Antibody-like ADP-Ribose Binding Proteins, Biochemistry, 56 (2017) 6305–6316.

[65] S. Oeck, N.M. Malewicz, A. Krysztofiak, A. Turchick, V. Jendrossek, P.M. Glazer, High-throughput Evaluation of Protein Migration and Localization after Laser Micro-Irradiation, Sci Rep, 9 (2019) 3148.

[66] C.A. Koczor, A.J. Haider, K.M. Saville, J. Li, J.F. Andrews, A.V. Beiser, R.W. Sobol, Live Cell Detection of Poly(ADP-Ribose) for Use in Genetic and Genotoxic Compound Screens, Cancers (Basel), 14 (2022).

[67] J.B. Tang, D. Svilar, R.N. Trivedi, X.H. Wang, E.M. Goellner, B. Moore, R.L. Hamilton, L.A. Banze, A.R. Brown, R.W. Sobol, N-methylpurine DNA glycosylase and DNA polymerase beta modulate BER inhibitor potentiation of glioma cells to temozolomide, Neuro-oncology, 13 (2011) 471–486.

[68] Q. Fang, B. Inanc, S. Schamus, X.H. Wang, L. Wei, A.R. Brown, D. Svilar, K.F. Sugrue, E.M. Goellner, X. Zeng, N.A. Yates, L. Lan, C. Vens, R.W. Sobol, HSP90 regulates DNA repair via the interaction between XRCC1 and DNA polymerase beta, Nature communications, 5 (2014) 5513.

[69] P. Desjardins, J.B. Hansen, M. Allen, Microvolume protein concentration determination using the NanoDrop 2000c spectrophotometer, Journal of visualized experiments : JoVE, (2009).

[70] W.A. Beard, S.H. Wilson, Purification and domain-mapping of mammalian DNA polymerase beta, Methods Enzymol, 262 (1995) 98–107.

[71] Z. Wang, G.A. Michaud, Z. Cheng, Y. Zhang, T.R. Hinds, E. Fan, F. Cong, W. Xu, Recognition of the iso-ADP-ribose moiety in poly(ADP-ribose) by WWE domains suggests a general mechanism for poly(ADP-ribosyl)ation-dependent ubiquitination, Genes Dev, 26 (2012) 235–240.

[72] F. He, K. Tsuda, M. Takahashi, K. Kuwasako, T. Terada, M. Shirouzu, S. Watanabe, T. Kigawa, N. Kobayashi, P. Guntert, S. Yokoyama, Y. Muto, Structural insight into the interaction of ADP-ribose with the PARP WWE domains, FEBS Lett, 586 (2012) 3858–3864.

[73] C. UniProt, UniProt: the universal protein knowledgebase in 2021, Nucleic Acids Res, 49 (2021) D480–D489.

[74] L.A. Aaltonen, P. Peltomaki, F.S. Leach, P. Sistonen, L. Pylkkanen, J.P. Mecklin, H. Jarvinen, S.M. Powell, J. Jen, S.R. Hamilton, et al., Clues to the pathogenesis of familial colorectal cancer, Science, 260 (1993) 812–816.

[75] M.D. Allen, A.M. Buckle, S.C. Cordell, J. Lowe, M. Bycroft, The crystal structure of AF1521 a protein from Archaeoglobus fulgidus with homology to the non-histone domain of macroH2A, J Mol Biol, 330 (2003) 503–511.

[76] C.K. Donawho, Y. Luo, Y. Luo, T.D. Penning, J.L. Bauch, J.J. Bouska, V.D. Bontcheva- Diaz, B.F. Cox, T.L. DeWeese, L.E. Dillehay, D.C. Ferguson, N.S. Ghoreishi-Haack, D.R. Grimm, R. Guan, E.K. Han, R.R. Holley-Shanks, B. Hristov, K.B. Idler, K. Jarvis, E.F. Johnson, L.R. Kleinberg, V. Klinghofer, L.M. Lasko, X. Liu, K.C. Marsh, T.P. McGonigal, J.A. Meulbroek, A.M. Olson, J.P. Palma, L.E. Rodriguez, Y. Shi, J.A. Stavropoulos, A.C. Tsurutani, G.D. Zhu, S.H. Rosenberg, V.L. Giranda, D.J. Frost, ABT-888, an orally active poly(ADP-ribose) polymerase inhibitor that potentiates DNA-damaging agents in preclinical tumor models, Clin Cancer Res, 13 (2007) 2728–2737.

[77] D.I. James, K.M. Smith, A.M. Jordan, E.E. Fairweather, L.A. Griffiths, N.S. Hamilton, J.R. Hitchin, C.P. Hutton, S. Jones, P. Kelly, A.E. McGonagle, H. Small, A.I. Stowell, J. Tucker, I.D. Waddell, B. Waszkowycz, D.J. Ogilvie, First-in-Class Chemical Probes against Poly(ADP-ribose) Glycohydrolase (PARG) Inhibit DNA Repair with Differential Pharmacology to Olaparib, ACS chemical biology, 11 (2016) 3179–3190.

[78] M. Mashimo, J. Kato, J. Moss, ADP-ribosyl-acceptor hydrolase 3 regulates poly (ADP-ribose) degradation and cell death during oxidative stress, Proc Natl Acad Sci U S A, 110 (2013) 18964–18969.

[79] F. Rosenthal, K.L. Feijs, E. Frugier, M. Bonalli, A.H. Forst, R. Imhof, H.C. Winkler, D. Fischer, A. Caflisch, P.O. Hassa, B. Luscher, M.O. Hottiger, Macrodomain-containing proteins are new mono-ADP-ribosylhydrolases, Nat Struct Mol Biol, 20 (2013) 502–507.

[80] N.V. Maluchenko, D.O. Koshkina, A.V. Feofanov, V.M. Studitsky, M.P. Kirpichnikov, Poly(ADP-Ribosyl) Code Functions, Acta Naturae, 13 (2021) 58–69.

[81] S.F. El-Khamisy, M. Masutani, H. Suzuki, K.W. Caldecott, A requirement for PARP-1 for the assembly or stability of XRCC1 nuclear foci at sites of oxidative DNA damage, Nucleic Acids Res, 31 (2003) 5526–5533.

[82] J. Fahrer, R. Kranaster, M. Altmeyer, A. Marx, A. Burkle, Quantitative analysis of the binding affinity of poly(ADP-ribose) to specific binding proteins as a function of chain length, Nucleic Acids Res, 35 (2007) e143.

[83] J.M. Reber, A. Mangerich, Why structure and chain length matter: on the biological significance underlying the structural heterogeneity of poly(ADP-ribose), Nucleic Acids Res, 49 (2021) 8432–8448.

[84] S. Challa, K.W. Ryu, A.L. Whitaker, J.C. Abshier, C.V. Camacho, W.L. Kraus, Development and characterization of new tools for detecting poly(ADP-ribose) in vitro and in vivo, Elife, 11 (2022).

[85] S. Challa, A.L. Whitaker, W.L. Kraus, Detecting Poly (ADP-Ribose) In Vitro and in Cells Using PAR Trackers, Methods Mol Biol, 2609 (2023) 75–90.

[86] H. Wang, Characterization and application of ubiquitin E3 ligase RNF146 WWE domain, in, John Hopkins University, John Hopkins University, 2016.

[87] H. Gaillard, T. Garcia-Muse, A. Aguilera, Replication stress and cancer, Nat Rev Cancer, 15 (2015) 276–289.

[88] M. Berti, D. Cortez, M. Lopes, The plasticity of DNA replication forks in response to clinically relevant genotoxic stress, Nat Rev Mol Cell Biol, 21 (2020) 633–651.

[89] C.K. Anders, E.P. Winer, J.M. Ford, R. Dent, D.P. Silver, G.W. Sledge, L.A. Carey, Poly(ADP-Ribose) polymerase inhibition: "targeted" therapy for triple-negative breast cancer, Clin Cancer Res, 16 (2010) 4702–4710.

[90] Z.D. Zhou, C.H. Chan, Z.C. Xiao, E.K. Tan, Ring finger protein 146/Iduna is a poly(ADP-ribose) polymer binding and PARsylation dependent E3 ubiquitin ligase, Cell Adh Migr, 5 (2011) 463–471.

[91] H.C. Kang, Y.I. Lee, J.H. Shin, S.A. Andrabi, Z. Chi, J.P. Gagne, Y. Lee, H.S. Ko, B.D. Lee, G.G. Poirier, V.L. Dawson, T.M. Dawson, Iduna is a poly(ADP-ribose) (PAR)-dependent E3 ubiquitin ligase that regulates DNA damage, Proc Natl Acad Sci U S A, 108 (2011) 14103–14108.

