## Supplement and Full blots for "Overexpression of the WWE domain of RNF146 modulates poly-(ADP)-ribose dynamics at sites of DNA damage"

**Running title**: RNF146 modulation of PAR dynamics

**Keywords**: poly-ADP-ribose (PAR), RNF146, WWE domain, Macrodomain, PAR binding domains

***Address correspondence to**:

Robert W. Sobol, Ph.D.
Department of Pathology and Laboratory Medicine
Warrant Alpert Medical School & Legorreta Cancer Center
Brown University, Providence, RI 02912


The following supporting information is available:

**Figure S1.** Recruitment dynamics of the WWE domains of RNF146(92-168), TRIP12(749-836), DTX(9-97), DTX(98-174) and DTX(9-174) to sites of laser micro-irradiation.

**Figure S2.** Recruitment dynamics of the RNF146(100-182) WWE domain and the engineered macrodomain Af1521(K35E/Y145R) to sites of laser micro-irradiation.

**Figure S3.** *In vitro* PAR binding assay of RNF146(100-182), Af1521(WT) and Af1521(K35E/Y145R).

**Figure S4.** Overexpression of the RNF146(100-182) WWE domain modulates PAR levels and PAR dynamics at sites of laser micro-irradiation in ES-2 cells.

**Figure S5.** Overexpression of the engineered macrodomain Af1521(K35E/Y145R) does not modulate PAR levels or PAR dynamics at sites of laser micro-irradiation in ES-2 cells.

**Figure S6.** WWE domain binding to sites of laser micro-irradiation is not influenced by the overexpression of the RNF146(100-182) WWE domain in ES-2 cells.

**Table S1:** Reagent List.

**Table S2:** Proteins encoding WWE domains.

#### Figure S1

**A.**

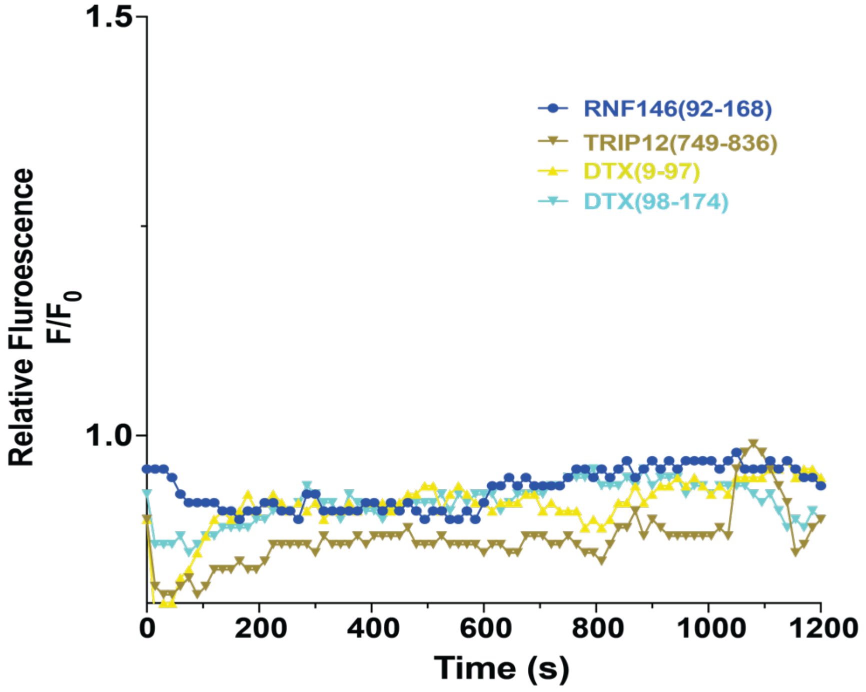

1.5
1.0
Relative Fluroescence F/F0
RNF146(92-168)
TRIP12(749-836)
DTX(9-97)
DTX(98-174)
0
200
400
600
800
1000
1200
Time (s)

**B.**

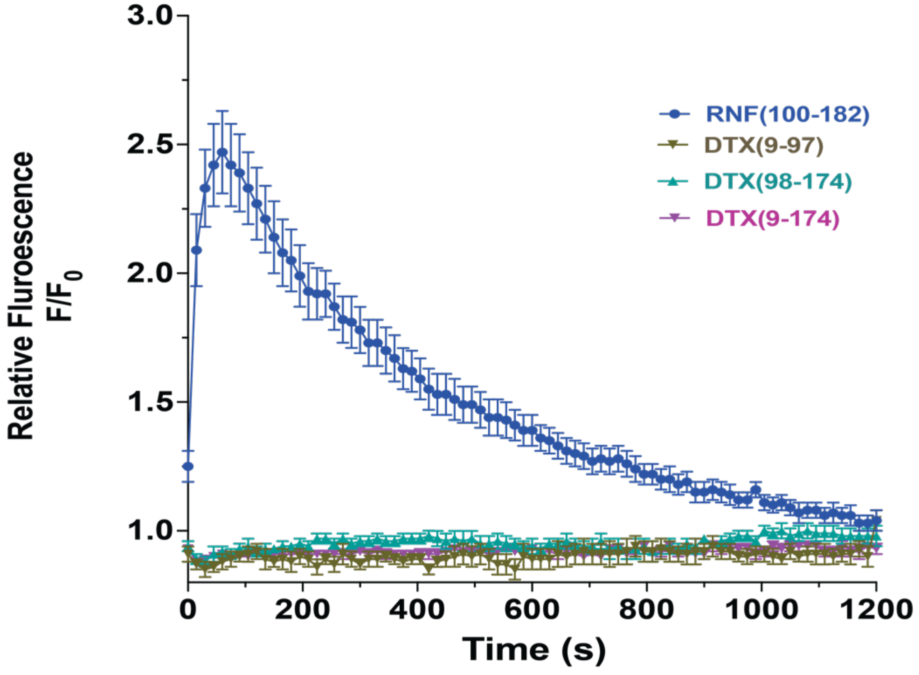

3.0
2.5
2.0
1.5
1.0
Relative Fluroescence F/F0
RNF(100-182)
DTX(9-97)
DTX(98-174)
DTX(9-174)
0
200
400
600
800
1000
1200
Time (s)

**Figure S1. Recruitment dynamics of the WWE domains of RNF146(92-168), TRIP12(749-836), DTX(9-97), DTX(98-174) and DTX(9-174) to sites of laser micro-irradiation.** (**A**) Plot of the recruitment of the WWE domains of RNF146(92-168), TRIP12(749-836), DTX(9-97), and DTX(98-174) to sites of laser micro-irradiation (405nm) without BrdU sensitization, N≥12 cells, ($F/F_0$: Maximum Fluorescence intensity / Fluorescence intensity at $t_0$); (**B**) Plot of the recruitment of the WWE domains of RNF146(92-168), DTX(9-97), DTX(98-174), and the tandem WWE domains of DTX(9-174) to sites of laser micro-irradiation (405nm) after BrdU sensitization (10μM, 24 hours), N≥12 cells, ($F/F_0$: Maximum Fluorescence intensity / Fluorescence intensity at $t_0$).

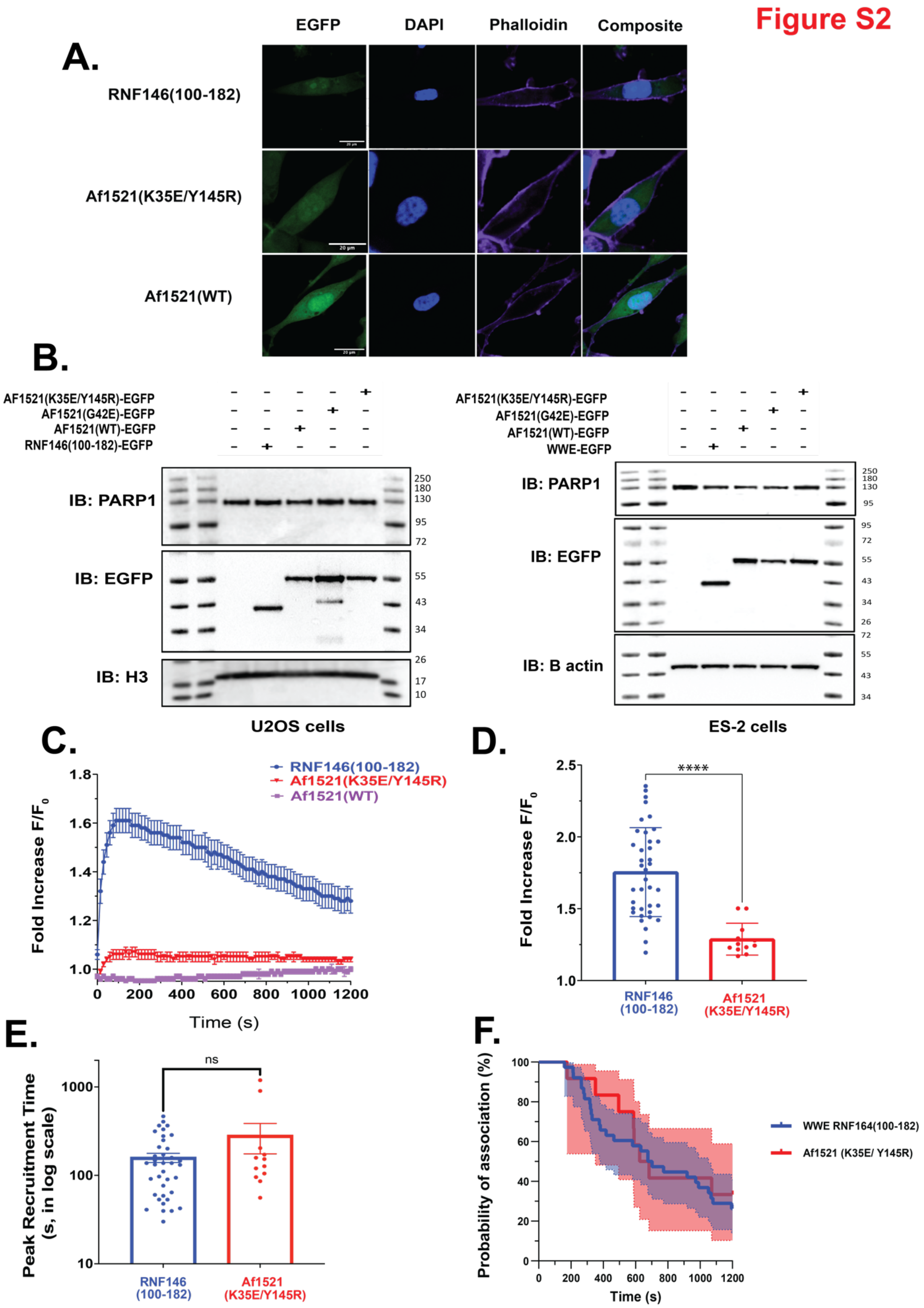

Figure S2
A.
EGFP
DAPI
Phalloidin
Composite
RNF146(100-182)
Af1521(K35E/Y145R)
Af1521(WT)
B.
AF1521(K35E/Y145R)-EGFP
AF1521(G42E)-EGFP
AF1521(WT)-EGFP
RNF146(100-182)-EGFP
IB: PARP1
IB: EGFP
IB: H3
U2OS cells
AF1521(K35E/Y145R)-EGFP
AF1521(G42E)-EGFP
AF1521(WT)-EGFP
WWE-EGFP
IB: PARP1
IB: EGFP
IB: B actin
ES-2 cells
C.
RNF146(100-182)
Af1521(K35E/Y145R)
Af1521(WT)
Fold Increase F/F0
Time (s)
D.
Fold Increase F/F0
****
RNF146 (100-182)
Af1521 (K35E/Y145R)
E.
Peak Recruitment Time (s, in log scale)
ns
RNF146 (100-182)
Af1521 (K35E/Y145R)
F.
Probability of association (%)
Time (s)
WWE RNF164(100-182)
Af1521 (K35E/ Y145R)

**Figure S2. Recruitment dynamics of the RNF146(100-182) WWE domain and the engineered macrodomain Af1521(K35E/Y145R) to sites of laser micro-irradiation.** (**A**) Confocal micrograph images of EGFP-fused RNF146(100-182), Af1521(WT), or Af1521(K35E/Y145R), expressed in ES-2 cells, white scale bar denotes 20μm; (**B**) Immunoblots of EGFP-fused RNF146(100-182), Af1521(WT), AF1521(G42E), or Af1521(K35E/Y145R), PARP1, H3 and actin using from whole cell protein lysates prepared from U2OS cells (Left) and ES-2 cells (Right). Molecular weight markers (kDa) are labeled on the right side; (**C**) Recruitment of the RNF146(100-182) WWE domain, the macrodomain Af1521(WT), and the engineered macrodomain Af1521(K35E/Y145R), to sites of laser micro-irradiation (405nm) without BrdU sensitization, N≥12 cells, ($F/F_0$: Maximum Fluorescence intensity / Fluorescence intensity at $t_0$); (**D**) Relative peak intensity of recruitment for the RNF146(100-182) WWE domain and the engineered macrodomain Af1521(K35E/Y145R), in U2OS cells. Each point represents a single cell recruitment focus, graph shows mean ± SEM. ($F/F_0$: Maximum Fluorescence intensity / Fluorescence intensity at $t_0$); (**E**) Peak recruitment time for the RNF146(100-182) WWE domain and the engineered macrodomain Af1521(K35E/Y145R), in U2OS cells. Each point represents a single cell recruitment focus, graph shows mean ± SEM; (**F**) Plot depicting the dissociation dynamics of RNF146(100-182), Af1521(K35E/Y145R) and Af1521(WT) foci in U2OS cells over 20 minutes following laser-induced DNA damage; N ≥ 40 cells. Recruitment foci with a relative peak intensity below 1.15 for the first frame were excluded from the experiment and from statistical analysis in graphs **D-F**. Exclusion percentages were 70% (28 foci) for Af1521(K35E/Y145R) foci and 5% (2 foci) for RNF146(100-182) foci. NS, no significance; *p<0.05, **p<0.01, ***p<0.001, ****p<0.0001; a Mann-Whitney test was used for panels **D** and **E**, and a Kaplan-Meier test for panel **F**.

#### Figure S3_Panels AB

**A.** **Protein purification scheme**

BL-21 cells expressing GST-tagged proteins.

↓

Protein expression induced by 0.4mM IPTG and incubated overnight at 16°C.

↓

Cells were sonicated and cell supernatant was collected.

↓

Cell lysates were mixed with Pierce glutathione superflow agarose overnight at 4°C with rotation.

↓

Glutathione agarose was washed and GST-tagged protein was eluted with glutathione.

↓

The mixture was passed through the Pierce spin column and flow-through was collected.

↓

The purity of the proteins was examined by SDS-PAGE.

**B.**

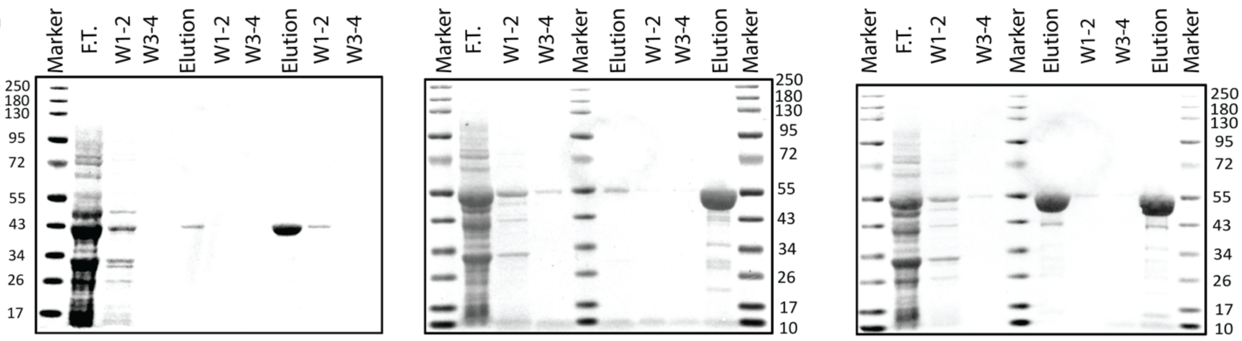

**WWE-GST** Coomassie stain

**AF1521(WT)-GST** Coomassie stain

**AF1521(K35E/Y145R)-GST** Coomassie stain

Figure S3_Panels CD

**C.** **PAR binding sandwich ELISA assay scheme**

Serial dilutions of GST-tagged proteins were added to Glutathione-coated plates (100μl/well) in PBS-Tween

↓

Plates incubated at room temperature for 1 hour

↓

Wells were washed 4 times with PBS-Tween

↓

100nM of PAR was added to each well (50 μl/well) and incubated overnight at 4C

↓

Wells were washed 4 times with PBS-Tween

↓

PAR primary antibody was added (1:200, 50 μl/well) and incubated for 2 hours at room temperature

↓

Wells were washed 4 times with PBS-Tween

↓

Secondary antibody was added (1:200, 50 μl/well) and incubated for 1 hour at room temperature

↓

Wells were washed 4 times with PBS-Tween

↓

HRP substrate was added and luminescence was measured

**D.**

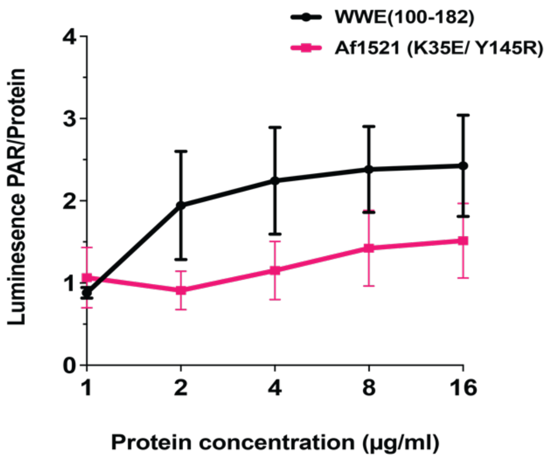

Figure S3_Panel E

E.

#### Scheme for *in vitro* PAR binding assay

PAR formation was induced by treating ES-2 cells with MNNG and PARGi then cell lysate was collected.

↓

PAR-containing cell lysate was serially diluted and placed into assigned blots on a nitrocellulose membrane using a slot blot device.

↓

The membrane was blocked with B-TBST (TBS buffer with 0.05% Tween-20 and supplemented with 5% blotting grade non-fat dry milk).

↓

Membrane was incubated with the assigned protein overnight at 4°C with rocking.

↓

Membrane was blotted with the GST antibodies in B-TBST for 2 hours at RT.

↓

Membranes were incubated with secondary antibodies in B-TBST for 1 hour (room temperature)

↓

Membranes were imaged using a Bio-Rad Chemi-Doc MP imaging system.

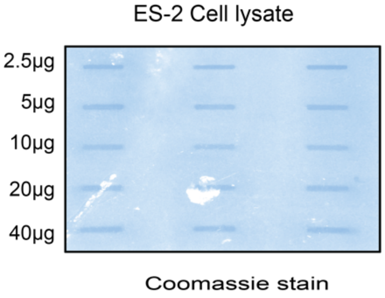

#### Figure S3_Panels F-M

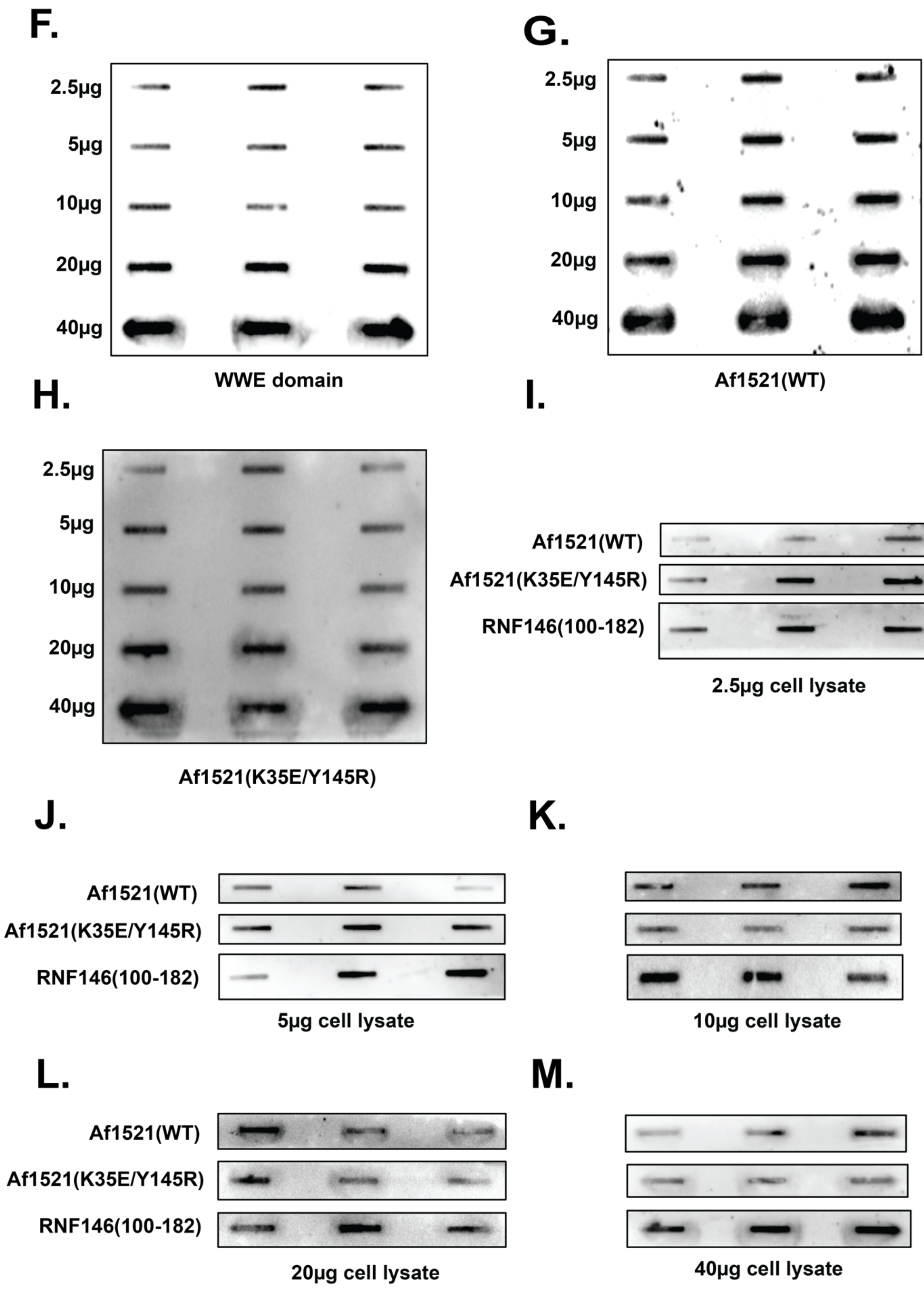

**Figure S3.** ***In vitro*** **PAR binding assay of RNF146(100-182), Af1521(WT), and Af1521(K35E/Y145R).** (**A**)The scheme used for recombinant protein expression and purification of the GST-tagged PAR binding domains in *E. coli*; (**B**) Purified PAR binding domains examined by Coomassie Blue staining following SDS-PAGE gel electrophoresis for GST-tagged RNF146(100-182) WWE domain (left), Af1521(WT) macrodomain (middle) and Af1521(K35E/Y145R) macrodomain (right); Molecular weights, as determined by marker proteins, are indicated; (**C**) The scheme used for *in vitro* PAR binding sandwich ELISA assay of RNF146(100-182) and Af1521(K35E/Y145R); (**D**) Plot showing the binding of PAR (100nM) to an increased concentration of RNF146(100-182) or Af1521(K35E/Y145R), PAR luminescence was normalized to luminescence of 1 μg/ml protein concentration; (**E**) The scheme used for the *in vitro* PAR binding assay of RNF146(100-182), Af1521(WT) and Af1521(K35E/Y145R) (Left) and PAR-containing lysate loading in three replicates as shown by Coomassie Blue staining (Right) of the membrane following transfer into the designated slot; (**F**) Immunoblots of purified GST-tagged RNF146(100-182) (100μg) binding to an increased concentration of PAR-containing ES-2 cell lysate (4°C, overnight) in three replicates as probed by GST primary antibody; (**G**) Immunoblots of purified GST-tagged Af1521(WT) (100μg) binding to an increased concentration of PAR-containing ES-2 cell lysate (4°C, overnight) in three replicates as probed by GST primary antibody; (**H**) Immunoblots of purified GST-tagged Af1521(K35E/Y145R) (100μg) binding to an increased concentration of PAR-containing ES-2 cell lysate (4°C, overnight) in three replicates as probed by GST primary antibody; (**I**) Immunoblots of RNF146(100-182), Af1521(WT), Af1521(K35E/Y145R) binding to 5μg PAR-containing cell lysate; (**J**) Immunoblots of RNF146(100-182), Af1521(WT), Af1521(K35E/Y145R) binding to 2.5μg PAR-containing cell lysate; (**K**) Immunoblots of RNF146(100-182), Af1521(WT), Af1521(K35E/Y145R) binding to 10μg PAR-containing cell lysate; (**L**) Immunoblots of RNF146(100-182), Af1521(WT), Af1521(K35E/Y145R) binding to 20μg PAR-containing cell lysate; (**M**) Immunoblots of RNF146(100-182), Af1521(WT), Af1521(K35E/Y145R) binding to 40μg PAR-containing cell lysate.

Figure S4

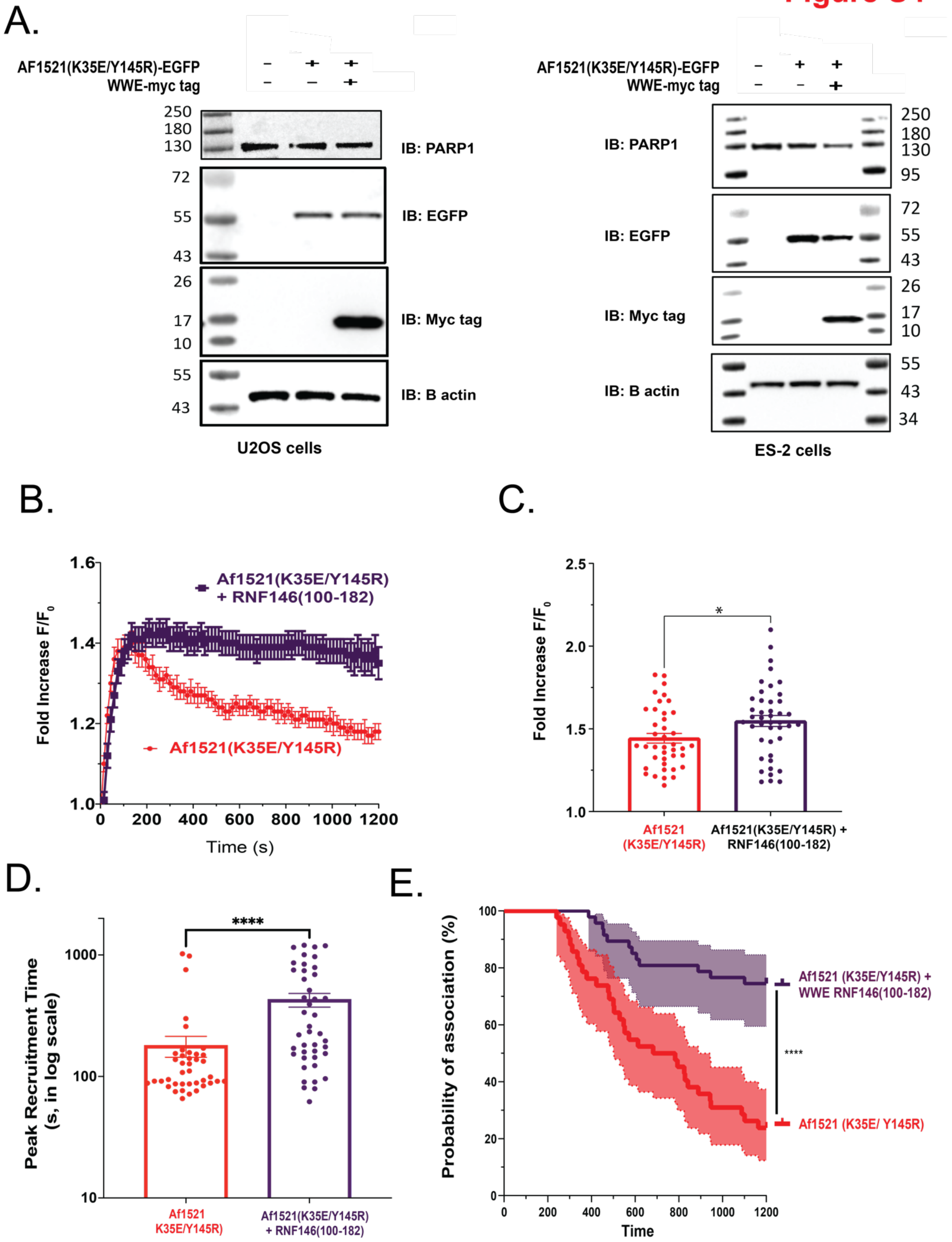
A.
AF1521(K35E/Y145R)-EGFP
WWE-myc tag
250
180
130
IB: PARP1
72
55
43
IB: EGFP
26
17
10
IB: Myc tag
55
43
IB: B actin
U2OS cells
AF1521(K35E/Y145R)-EGFP
WWE-myc tag
IB: PARP1
250
180
130
95
IB: EGFP
72
55
43
IB: Myc tag
26
17
10
IB: B actin
55
43
34
ES-2 cells
B.
Af1521(K35E/Y145R)
+ RNF146(100-182)
Af1521(K35E/Y145R)
Fold Increase F/F0
Time (s)
C.
Fold Increase F/F0
*
Af1521
(K35E/Y145R)
Af1521(K35E/Y145R) +
RNF146(100-182)
D.
****
Peak Recruitment Time
(s, in log scale)
Af1521
K35E/Y145R)
Af1521(K35E/Y145R)
+ RNF146(100-182)
E.
Probability of association (%)
Time
Af1521 (K35E/Y145R) +
WWE RNF146(100-182)
****
Af1521 (K35E/ Y145R)

**Figure S4. Overexpression of the RNF146(100-182) WWE domain modulates PAR levels and dynamics at sites of laser micro-irradiation in ES-2 cells.** (**A**) Immunoblots of RNF146(100-182) (fused to a myc tag), Af1521(K35E/Y145R) (fused to EGFP), PARP1, and β-actin using whole cell protein lysates prepared from U2OS cells (Left) and ES-2 cells (Right). Molecular weight markers (kDa) are labeled as indicated; (**B**) Recruitment of the engineered macrodomain Af1521(K35E/Y145R) in ES-2 cells, after overexpression of the myc-tagged RNF146(100-182) WWE domain, to sites of laser micro-irradiation (405nm), N≥40 cells, ($F/F_0$: Maximum Fluorescence intensity / Fluorescence intensity at $t_0$); (**C**) Relative peak intensity of recruitment for the engineered macrodomain Af1521(K35E/Y145R), after overexpression of the RNF146(100-182) WWE domain, in ES-2 cells. Each point represents a single cell recruitment focus. Graph shows mean ± SEM. ($F/F_0$: Maximum Fluorescence intensity/ Fluorescence intensity at $t_0$); (**D**) Peak recruitment time for the engineered macrodomain Af1521(K35E/Y145R), after overexpression of the myc-tagged RNF146(100-182) WWE domain, in ES-2 cells. Each point represents a single cell recruitment focus, graph shows mean ± SEM; (**E**) Plot depicting the dissociation dynamics of Af1521(K35E/Y145R) foci after overexpression of RNF146(100-182) in ES-2 cells over 20 minutes following laser-induced DNA damage; N ≥ 40 cells. Af1521(K35E/Y145R) EGFP foci with a relative peak intensity below 1.15 for the first frame were excluded from the experiment and from statistical analysis in graphs **C-E**. Exclusion percentages were 24% (12 foci) for Af1521(K35E/Y145R) foci and 10% (5 foci) for Af1521(K35E/Y145R) foci after overexpression of RNF146(100-182). NS, no significance; *p<0.05, **p<0.01, ***p<0.001, ****p<0.0001; a two-sample t-test was used for panel **C**, Mann-Whitney test for panel **D** and a Kaplan-Meier test for panel **E**.

### Figure S5

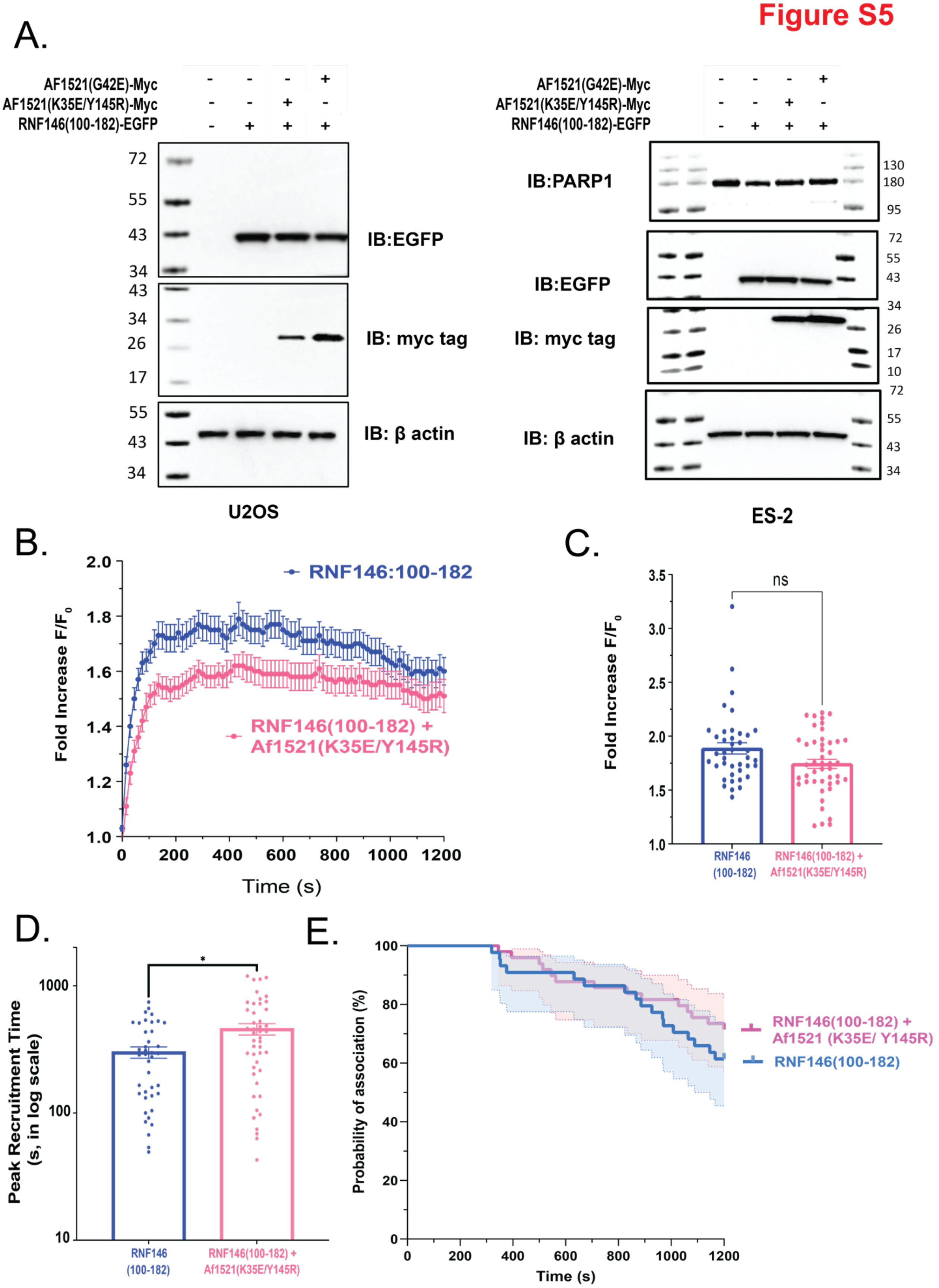

A.
AF1521(G42E)-Myc
AF1521(K35E/Y145R)-Myc
RNF146(100-182)-EGFP
IB:EGFP
IB: myc tag
IB: β actin
U2OS
IB:PARP1
ES-2
B.
RNF146:100-182
RNF146(100-182) + Af1521(K35E/Y145R)
Fold Increase F/F0
Time (s)
C.
ns
RNF146 (100-182)
RNF146(100-182) + Af1521(K35E/Y145R)
D.
Peak Recruitment Time (s, in log scale)
E.
Probability of association (%)
RNF146(100-182) + Af1521 (K35E/ Y145R)
RNF146(100-182)

**Figure S5. Overexpression of the engineered macrodomain Af1521(K35E/Y145R) does not modulate PAR levels or PAR dynamics at sites of laser micro-irradiation in ES-2 cells.** (**A**) Immunoblots of RNF146(100-182) (fused to EGFP), Af1521(G42E) (fused to a myc tag), Af1521(K35E/Y145R) (fused to a myc tag), PARP1, and β-actin using whole cell protein lysates prepared from U2OS cells (Left) and ES-2 cells (Right). Molecular weight markers (kDa) are labeled as indicated; (**B**) Recruitment of the RNF146(100-182) WWE domain in ES-2 cells, after overexpression of the engineered macrodomain Af1521(K35E/Y145R), to sites of laser micro-irradiation (405nm), N≥40 cells. ($F/F_0$: Maximum Fluorescence intensity / Fluorescence intensity at $t_0$); (**C**) Relative peak intensity of recruitment for the RNF146(100-182) WWE domain, after overexpression of the engineered myc-tagged macrodomain Af1521(K35E/Y145R), in ES-2 cells at sites of laser micro-irradiation (405nm). Each point represents a single cell recruitment focus, graph shows mean ± SEM. ($F/F_0$: Maximum Fluorescence intensity / Fluorescence intensity at $t_0$); (**D**) Peak recruitment time for the RNF146(100-182) WWE domain, after overexpression of the engineered macrodomain Af1521(K35E/Y145R), in ES-2 cells at sites of laser micro-irradiation (405nm). Each point represents a single cell recruitment focus, graph shows mean ± SEM; (**E**) Plot depicting the dissociation dynamics of RNF146(100-182) foci after overexpression of Af1521(K35E/Y145R) in ES-2 cells over 20 minutes following laser-induced DNA damage; N ≥ 38 cells. RNF146(100-182)-EGFP foci with a relative peak intensity below 1.15 for the first frame were excluded from the experiment and from statistical analysis in graphs **C**-**E**. Exclusion percentages were 0% for RNF146(100-182) foci and 5% (2 foci) for RNF146(100-182) foci after overexpression of Af1521(K35E/Y145R). NS, no significance; *p<0.05, **p<0.01, ***p<0.001, ****p<0.0001; Mann-Whitney test for panels **C** and **D** and a Kaplan-Meier test for panel **E**.

#### Figure S6

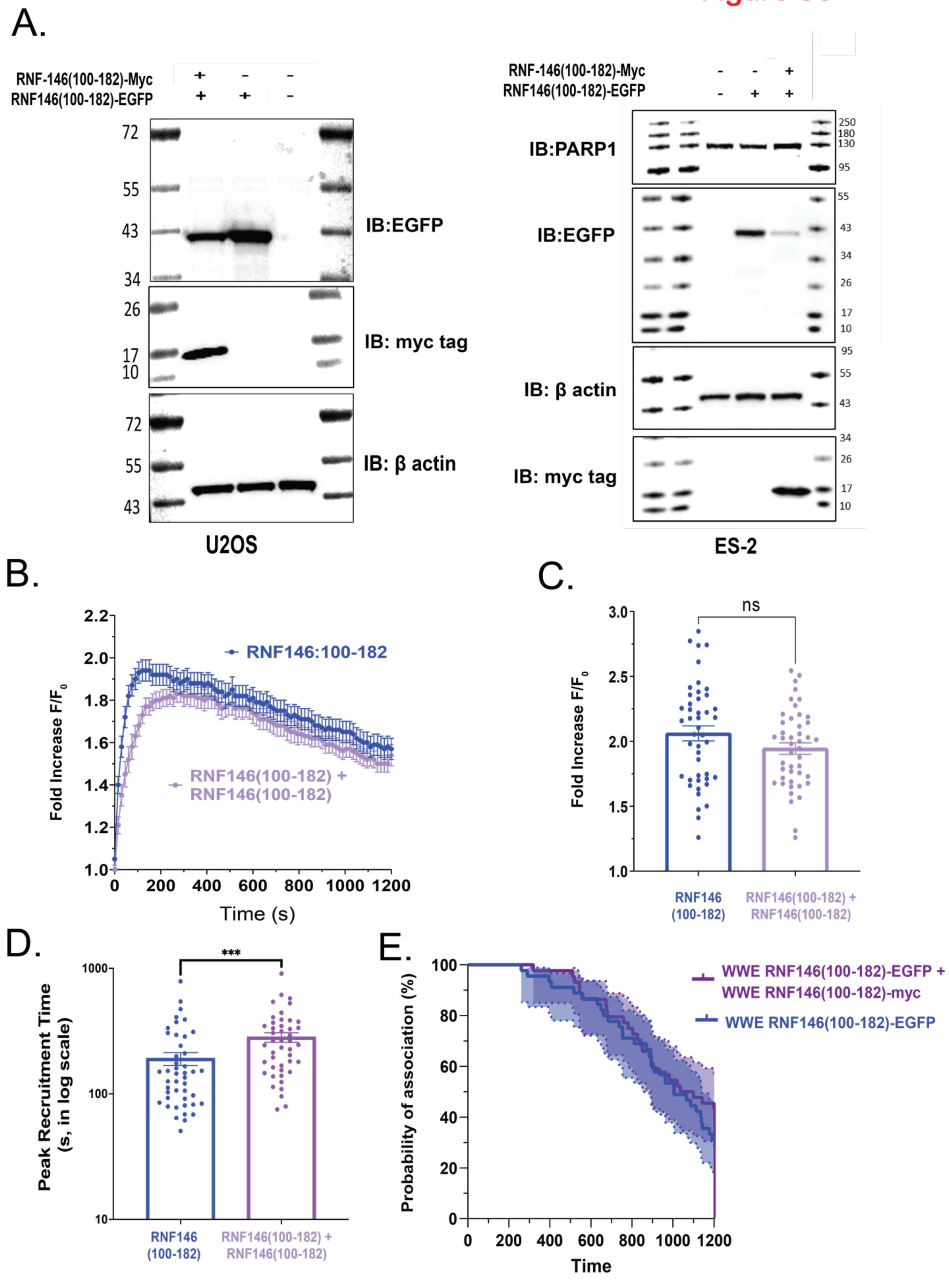
A.
RNF-146(100-182)-Myc
RNF146(100-182)-EGFP
IB:EGFP
IB: myc tag
IB: β actin
U2OS
IB:PARP1
ES-2
B.
RNF146:100-182
RNF146(100-182) + RNF146(100-182)
Fold Increase F/F0
Time (s)
C.
ns
RNF146 (100-182)
RNF146(100-182) + RNF146(100-182)
D.
***
Peak Recruitment Time (s, in log scale)
E.
WWE RNF146(100-182)-EGFP + WWE RNF146(100-182)-myc
WWE RNF146(100-182)-EGFP
Probability of association (%)
Time

**Figure S6. WWE domain binding to sites of laser micro-irradiation is not saturated by the overexpression of the RNF146(100-182) WWE domain in ES-2 cells.** (**A**) Immunoblots of RNF146-(100-182) (fused to EGFP), RNF146-(100-182) (fused to a myc tag), PARP1, and β-actin using whole cell protein lysates prepared from U2OS cells (Left) and ES-2 cells (Right). Molecular weight markers (kDa) are labeled as indicated; (**B**) Recruitment of the RNF146(100-182) WWE domain in ES-2 cells, after overexpression of a competing RNF146(100-182) WWE domain (fused to a myc tag), to sites of laser micro-irradiation (405nm, N≥45 cells), ($F/F_0$: Maximum Fluorescence intensity / Fluorescence intensity at $t_0$); (**C**) Relative peak intensity of recruitment for the RNF146(100-182) WWE domain, after overexpression of a competing RNF146(100-182) WWE domain (fused to a myc tag), in ES-2 cells at sites of laser micro-irradiation (405nm). Each point represents a single cell recruitment focus, graph shows mean ± SEM. ($F/F_0$: Maximum Fluorescence intensity / Fluorescence intensity at $t_0$); (**D**) Peak recruitment time for the RNF146(100-182) WWE domain, after overexpression of a competing RNF146(100-182) WWE domain (fused to a myc tag), in ES-2 cells at sites of laser micro-irradiation (405nm). Each point represents a single cell recruitment focus, graph shows mean ± SEM; (**E**) Plot depicting the dissociation dynamics of RNF146(100-182) foci after overexpression of RNF146(100-182)-myc in ES-2 cells over 20 minutes following laser-induced DNA damage. N ≥ 45 cells. RNF146(100-182)-EGFP foci with a relative peak intensity below 1.15 for the first frame were excluded from the experiment and from statistical analysis in graphs **C-E**. Exclusion percentages were 6.7% (3 foci) for RNF146(100-182) foci and 11.1% (5 foci) for RNF146(100-182) foci after overexpression of RNF146(100-182)-myc. NS, no significance; *p<0.05, **p<0.01, ***p<0.001, ****p<0.0001; a t-test was used for panel **C,** Mann-Whitney test for panel **D** and a Kaplan-Meier test for panel **E**.

| Table S1: Reagent List | SOURCE | IDENTIFIER |
|---|---|---|
| **Antibodies** | | |
| Rabit anti-PARP1 (Immunoblot; 1:1000) | Abcam | Cat# 191217 |
| Rabbit anti-GFP (Immunoblot; 1:1000) | Chromotek | Cat# pabg1-100 |
| Rabbit anti-H3 (Immunoblot; 1:2000) | Abcam | Cat# ab1791 |
| Rabbit anti-$\beta$-actin (Immunoblot; 1:2000) | CST | Cat# 8457S |
| Rabbit anti-myc (Immunoblot; 1:2000) | CST | Cat# 71D10 |
| Mouse anti-PAR (10H) (Immunoblot; 1:1000) | Generous gift from Mathias Ziegler (University of Bergen, Norway) | N/A |
| Goat anti-mouse-HRP conjugate (Immunoblot; 1:2000) | Bio-Rad | Cat# 1706516 |
| Goat anti-rabbit-HRP conjugate (Immunoblot; 1:2000) | Bio-Rad | Cat# 1706515 |
| HRP-conjugated streptavidin (Immunoblot; 1:2000) | Thermo Fisher Scientific | Cat# N100 |
| GST-Biotin (Immunoblot; 1:1000) | Invitrogen | Cat# MA4-004-Btin |
| **Chemicals, peptides, and recombinant proteins** | | |
| Heat-inactivated fetal bovine serum | Bio-Techne | Cat# S11150H |
| Penicillin/streptomycin | Thermo Fisher Scientific | Cat# 15140-122 |
| DMEM | Corning | Cat# 15-017-CV |
| McCoy's 5A medium (1X) | Thermo Fisher Scientific | Cat# 16600-082 |
| MEM Alpha (1X) | Thermo Fisher Scientific | Cat# 12571-063 |
| L-glutamine | Thermo Fisher Scientific | Cat# 25030-081 |
| Dimethyl Sulfoxide | Thermo Fisher Scientific | Cat# BP231-1 |
| Hygromycin | Thermo Fisher Scientific | Cat# 10687010 |
| Trypsin-EDTA | Thermo Fisher Scientific | Cat# 25200-056 |
| 0.45μM nitrocellulose | Bio-Rad | Cat# 162-0115 |
| 0.45μM Durapore Steriflip Filters | Sigma-Aldrich | Cat# SE1M003M00 |
| Polybrene | Sigma-Aldrich | Cat# 107689 |
| Protease inhibitor cocktail | Sigma-Aldrich | Cat# P8340 |
| Blotting grade non-fat dry milk | Bio-Rad | Cat# 170-6404 |
| Nupage 4-12% Bis-Tris gel | Invitrogen | Cat# NP0323BOX |
| Clarity Western ECL Substrate | Bio-Rad | Cat# 1705060 |
| SuperSignal West Femto Maximum Sensitivity Substrate | Thermo Fisher Scientific | Cat# 34095 |
| DC protein assay kit | Bio-Rad | Cat# 5000112 |

| | | |
|---|---|---|
| Formaldehyde solution (37%) | Thermo Fisher Scientific | Cat# BP531-500 |
| NucBlue Live Cell ReadyProbes (Hoechst 33342) | Thermo Fisher Scientific | Cat# R37605 |
| NucBlue Fixed Cell ReadyProbes (DAPI) | Thermo Fisher Scientific | Cat# R37606 |
| Alexa Fluor 647 Phalloidin | Thermo Fisher Scientific | Cat# A22287 |
| TransIT-X2 Transfection Reagent | Mirus Bio | Cat# MIR 6005 |
| ABT-888 (Veliparib) | Selleckchem | Cat# S1004 |
| PDD00017273 | Sigma-Aldrich | Cat# SML1781 |
| MNNG | Sigma-Aldrich | Cat# 129941 |
| Bromodeoxyuridine (BrdU) | Sigma-Aldrich | Cat# B5002 |
| Glutathione | Takara | Cat# 635619 |
| QIAprep Spin Miniprep Kit | Qiagen | Cat# 27106 |
| Formaldehyde solution (37%) | Thermo Fisher Scientific | Cat# BP531-500 |
| FastDigest MluI | Thermo Fisher Scientific | Cat# FD0564 |
| FastDigest BamHI | Thermo Fisher Scientific | Cat# FD0054 |
| FastDigest XpaI | Thermo Fisher Scientific | Cat# FD1383 |
| T7 DNA Ligase | New England Biolabs | Cat# M0318 |
| 10X TBE Electrophoresis Buffer | Thermo Fisher Scientific | Cat# B52 |
| OneTaq® Quick-Load® 2X Master Mix with Standard Buffer | New England Biolabs | Cat# M0486S |
| IPTG | Sigma-Aldrich | Cat# 16758-5G |
| NaCL | Fisher Bioreagents | Cat# BP358-212 |
| EDTA | Promega | Cat# V4231 |
| GST sepharose 4B resin | Sigma-Aldrich | Cat# GE17-0756-01 |
| Gelcode Blue Safe Protein Stain | Thermo Fisher Scientific | Cat# 18605971 |
| One Shot™ STBL 3 Chemically Competent E. coli | Thermo Fisher Scientific | Cat# C737303 |
| Glutathione coated plates | Thermo Fisher Scientific | Cat#15240 |
| Pierce™ GST Protein Interaction Pull-Down Kit | Thermo Fisher Scientific | Cat# 21516 |
| **Cell Lines** | | |
| U2OS<br>(Human osteosarcoma tumor cell line) | ATCC | Cat# HTB-96 |

| | | |
|---|---|---|
| ES-2<br>(Human ovarian tumor cell line) | ATCC | Cat# CRL-1987 |
| LN428<br>(Human glioblastoma tumor cell line) | Generous gift from Dr. Ian Pollack (University of Pittsburgh) | N/A |
| 293FT<br>(A human embryonal kidney cell line transformed with SV40 large T antigen) | Thermo Fisher Scientific | Cat# R70007 |
| U2OS/RNF146(100-182)-myc<br>(U2OS cells expressing RNF146(100-182)-myc) | This study | N/A |
| U2OS/Af1521(K35E/Y145R)-myc<br>(U2OS cells expressing Af1521(K35E/Y145R)-myc) | This study | N/A |
| U2OS/Af1521(G42E)-myc<br>(U2OS cells expressing Af1521(G42E)-myc) | This study | N/A |
| ES-2/RNF146(100-182)-myc<br>(ES-2 cells expressing RNF146(100-182)-myc) | This study | N/A |
| ES-2/Af1521(K35E/Y145R)-myc<br>(ES-2 cells expressing Af1521(K35E/Y145R)-myc) | This study | N/A |
| ES-2/Af1521(G42E)-myc<br>(ES-2 cells expressing Af1521(G42E)-myc) | This study | N/A |
| **Recombinant DNA (plasmids)** | | |
| pMDLg/pRRE (packaging vector for lentiviral production) | Lab stock 253, [1] | Addgene,<br>Cat# 1225 |
| pRSV-Rev (packaging vector for lentiviral production) | Lab stock 254, [1] | Addgene,<br>Cat# 12253 |
| pMD2.G (packaging vector for lentiviral production) | Lab stock 252, [1] | Addgene,<br>Cat# 12259 |
| pLV-EF1A-LivePAR-Hygro<br>(RNF146(100-182) WWE domain with EGFP tag & a hygromycin resistance cassette) | Lab stock 1727, [2] | Addgene,<br>Cat# 176063 |
| pLV-Hygro-EF1A-RealPARBackbone<br>(Expression vector with a BamHI site in frame with a Gly-Ser linker fused to EGFP; serves as the backbone for PAR binding domain incorporation for LivePAR) | Lab stock 1749 | [2] |
| pLV-Hygro-EF1A-WWE-Linker-EGFP<br>(EGFP fused to the C-terminus of a WWE domain & a hygromycin resistance cassette) | Lab stock 1770 | [2] |
| pLV-Hygro-EF1A-RNF146(92-168)-EGFP<br>(EGFP fused to the C-terminus of a WWE domain within RNF146 & a hygromycin resistance cassette) | This study;<br>Lab stock 2361 | Addgene,<br>Cat# 196228 |
| pLV-Hygro-EF1A-TRIP12(749-836)-EGFP<br>(EGFP fused to the C-terminus of a WWE domain within TRIP12 & a hygromycin resistance cassette) | This study;<br>Lab stock 2362 | Addgene,<br>Cat# 196229 |
| pLV-Hygro-EF1A-DTX2(8-97)-EGFP | This study; | Addgene, |

| | | |
|---|---|---|
| (EGFP fused to the C-terminus of the first WWE domain within Deltex2 & a hygromycin resistance cassette) | Lab stock 2366 | Cat# 196230 |
| pLV-Hygro-EF1A-DTX2(98-174)-EGFP (EGFP fused to the C-terminus of the second WWE domain within Deltex2 & a hygromycin resistance cassette) | This study; Lab stock 2367 | Addgene, Cat# 196231 |
| pLV-Hygro-EF1A-DTX2(8-174)-EGFP (EGFP fused to the C-terminus of tandem WWE domains within Deltex2 & a hygromycin resistance cassette) | This study; Lab stock 2368 | Addgene, Cat# 196232 |
| pLV-Hygro-EF1A-Af1521(WT)-EGFP (EGFP fused to the C-terminus of Af1521 (WT) macrodomain & a hygromycin resistance cassette) | This study; Lab stock 2179 | Addgene, Cat# 196233 |
| pLV-Hygro-EF1A-Af1521(G42E)-EGFP (EGFP fused to the C-terminus of Af1521(G42E) macrodomain & a hygromycin resistance cassette) | This study; Lab stock 2180 | Addgene, Cat# 196234 |
| pLV-Hygro-EF1A-Af1521(K35E/Y145R)-EGFP (EGFP fused to the C-terminus of Af1521(K35E/Y145R) macrodomain & a hygromycin resistance cassette) | This study; Lab stock 2181 | Addgene, Cat# 196235 |
| pLV-Hygro-EF1A-WWE(RNF146)-myc (RNF146(100-182) WWE domain with a myc tag fused to the C terminus & a hygromycin resistance cassette) | This study; Lab stock 2285 | Addgene, Cat# 196236) |
| pLV-Hygro-EF1A-Af1521(K35E/Y145R)-myc (Af1521(K35E/Y145R) macrodomain with a myc tag fused to the C terminus & a hygromycin resistance cassette) | This study; Lab stock 2286 | Addgene, Cat# 196237 |
| pLV-Hygro-EF1A-Af1521(G42E)-myc (Af1521(G42E) macrodomain with a myc tag fused to the C terminus & a hygromycin resistance cassette) | This study; Lab stock 2287 | Addgene, Cat# 196238 |
| pGEX-4T-3-GST-Tev-Af1521-myc (N-terminal GST fusion of Af1521(WT) with a TEV protease site located between the GST tag and Af1521 and a myc tag on the C terminus) | This study; Lab stock 2458 | Addgene, Cat# 196239 |
| pGEX-4T-3-GST-Tev-WWE(RNF146)-myc (N-terminal GST fusion of RNF146(100-182) WWE domain with a TEV protease site located between the GST tag and Af1521 and a myc tag on the C terminus) | This study; Lab stock 2459 | Addgene, Cat# 196240 |
| pGEX-4T-3-GST-Tev-Af1521(K35E Y145R)-myc (N-terminal GST fusion of Af1521(K35E/Y145R) with a TEV protease site located between the GST tag and Af1521 and a myc tag on the C terminus) | This study; Lab stock 2460 | Addgene, Cat# 196241 |
| **Software and Algorithms** | | |
| Image J | Image J 1.48v | https://imagej.nih.gov/ij/ 1.6.0_65 |
| FIJI | http://fiji.sc/ | [3] |

| | | |
|---|---|---|
| Adobe Illustrator (for preparation of figures) | Adobe Systems | Version 2024 |
| GraphPad Prism | GraphPad | Version 9, (Mac OS X) |
| MIDAS | [2] | https://doi.org/10.5281/zenodo.5534950 |
| NIS-Elements | Nikon Instruments | Versions 4.51 and 5.11 |
| Image lab | BioRad | Version 6.0.0 build 26, 2017 |
| BioRender (for graphics development) | BioRender | Online version |

**Table S2: Proteins encoding WWE domains**

| Gene | Uniprot ID | Full length protein | Amino acids of WWE | WWE Amino acid Sequence from Uniprot |
|---|---|---|---|---|
| RNF146 | Q9NTX7 | 359 aa | 92-168 | EELKAASRGNGEYAWYYEGRNGWWQYDERTSRELEDAFSKGKKNTEMLIAGFLYVADLENMVQYRRNEHGRRRKIKR |
| DTX1 | Q86Y01 | 620 aa | 14-94 | GLGFPPQNVARVVVWEWLNEHSRWRPYTATVCHHIENVLKEDARGSVVLGQVDAQLVPYIIDLQSMHQFRQDTGTMRPVRR |
| DTX1 | Q86Y01 | 620 aa | 95-171 | NFYDPSSAPGKGIVWEWENDGGAWTAYDMDICITIQNAYEKQHPWLDLSSLGFCYLIYFNSMSQMNRQTRRRRRLRR |
| DTX2 | Q86UW9 | 622 aa | 8 to 97 | SLVQVYTSPAAVAVWEWQDGLGTWHPYSATVCSFIEQQFVQQKGQRFGLGSLAHSIPLGQADPSLAPYIIDLPSWTQFRQDTGTMRAVRR |
| DTX2 | Q86UW9 | 622 aa | 98-174 | HLFPQHSAPGRGVVWEWLSDDGSWTAYEASVCDYLEQQVARGNQLVDLAPLGYNYTVNYTTHTQTNKTSSFCRSVRR |
| DTX4 | Q9Y2E6 | 619 aa | 1 to 78 | MLLASAVVVWEWLNEHGRWRPYSPAVSHHIEAVVRAGPRAGGSVVLGQVDSRLAPYIIDLQSMNQFRQDTGTLRPVRR |
| DTX4 | Q9Y2E6 | 619 aa | 79-155 | NYYDPSSAPGKGVVWEWENDNGSWTPYDMEVGITIQHAYEKQHPWIDLTSIGFSYVIDFNTMGQINRQTQRQRRVRR |
| TRIP12 | Q14669 | 1992 aa | 749-836 (Note, in isoform a mRNA, the domain runs from 798-884) | MLKKGNAQNTDGAIWQWRDDRGLWHPYNRIDSRIIEQINEDTGTARAIQRKPNPLANSNTSGYSESKKDDARAQLMKEDPELAKSFIK |

| | | | | |
|---|---|---|---|---|
| HUWE1 | Q7Z6Z7 | 4374 aa | 1603-1680 | RAQMTKYLQSNSNNWRWFDDRSGR WCSYSASNNSTIDSAWKSGETSVRFT AGRRRYTVQFTTMVQVNEETGNRRPV ML |
| PARP11 | Q9NR21 | 338 aa | 22-106 | NEVDDMDTSDTQWGWFYLAECGKWH MFQPDTNSQCSVSSEDIEKSFKTNPCG SISFTTSKFSYKIDFAEMKQMNLTTGKQ RLIKR |
| PARP12 | Q9H0J9 | 701 aa | 298-361 | PYRWQFLDRGKWEDLDNMELIEEAYC NPKIERILCSESASTFHSHCLNFNAMTY GATQARRLST |
| PARP12 | Q9H0J9 | 701 aa | 364-458 | SVTKPPHFILTTDWIWYWSDEFGSWQ EYGRQGTVHPVTTVSSSDVEKAYLAY CTPGSDGQAATLKFQAGKHNYELDFK AFVQKNLVYGTTKKVCR |
| PARPT | Q7Z3E1 | 657 aa | 332-410 | STPPSSNVNSIYHTVWKFFCRDHFGW REYPESVIRLIEEANSRGLKEVRFMMW NNHYILHNSFFRREIKRRPLFRSCFI |
| PARP14 | Q460N5 | 1801 aa | 1523-1601 | EQESRADCISEFIEWQYNDNNTSHCFN KMTNLKLEDARREKKKTVDVKINHRHY TVNLNTYTATDTKGHSLSVQRLTKS |
| DDHD2 | O94830 | 711 aa | 30-112 | DMDAGSLYEPVSPHWFYCKIIDSKETW IPFNSEDSQQLEEAYSSGKGCNGRVV PTDGGRYDVHLGERMRYAVYWDELA SEVRR |
| ZCCHV | Q7Z2W4 | 902 aa | 594-681 | SVTKPANSVFTTKWIWYWKNESGTWI QYGEEKDKRKNSNVDSSYLESLYQSC PRGVVPFQAGSRNYELSFQGMIQTNIA SKTQKDVIR |

**References**

[1] D.A. Rubinson, C.P. Dillon, A.V. Kwiatkowski, C. Sievers, L. Yang, J. Kopinja, D.L. Rooney, M. Zhang, M.M. Ihrig, M.T. McManus, F.B. Gertler, M.L. Scott, L. Van Parijs, A lentivirus-based system to functionally silence genes in primary mammalian cells, stem cells and transgenic mice by RNA interference, Nat Genet, 33 (2003) 401-406.

[2] C.A. Koczor, K.M. Saville, J.F. Andrews, J. Clark, Q. Fang, J. Li, R.Q. Al-Rahahleh, M. Ibrahim, S. McClellan, M.V. Makarov, M.E. Migaud, R.W. Sobol, Temporal dynamics of base excision/single-strand break repair protein complex assembly/disassembly are modulated by the PARP/NAD(+)/SIRT6 axis, Cell reports, 37 (2021) 109917.

[3] J. Schindelin, I. Arganda-Carreras, E. Frise, V. Kaynig, M. Longair, T. Pietzsch, S. Preibisch, C. Rueden, S. Saalfeld, B. Schmid, J.Y. Tinevez, D.J. White, V. Hartenstein, K. Eliceiri, P. Tomancak, A. Cardona, Fiji: an open-source platform for biological-image analysis, Nat Methods, 9 (2012) 676-682.

### Full Blot Images for:

#### Overexpression of the WWE domain of RNF146 modulates poly-(ADP)-ribose dynamics at sites of DNA damage

Rasha Q. Al-Rahahleh[1,2], Kate M. Saville[2], Joel F. Andrews[2],

Zhijin Wu[3], Christopher A. Koczor[2], and Robert W. Sobol[1,2]*

[1]Department of Pathology and Laboratory Medicine, Warren Alpert Medical School & Legorreta Cancer Center, Brown University, Providence, RI 02912

[2]Department of Pharmacology & Mitchell Cancer Institute, College of Medicine, University of South Alabama, Mobile, AL 36604, USA

[3]Department of Biostatistics, Brown University, Providence, RI 02912

**Running title**: RNF146 modulation of PAR dynamics

**Keywords**: poly-ADP-ribose (PAR), RNF146, WWE domain, Macrodomain, PAR binding domains

***Address correspondence to**:

Robert W. Sobol, Ph.D.
Department of Pathology and Laboratory Medicine
Warrant Alpert Medical School & Legorreta Cancer Center
Brown University, Providence, RI 02912


**Uncropped immunoblot images.** Each panel shows the uncropped images of the immunoblots (or Coomassie stained gels) following SDS-PAGE or slot-blot, as indicated in the corresponding figures. Gray lines denote the edge of the membrane analyzed when membrane borders are not clear; black lines denote the borders of the cropped image displayed in the main figure. Molecular weight markers are indicted on the left for the immunoblots following SDS-PAGE. The antigen probed with the corresponding antibody is indicated on the right.

(**A**) Raw data of immunoblots following slot-blot analysis from Figure 3F.

(**B**) Raw data of immunoblots following separation by SDS-PAGE from Figure 4F.

(**C**) Raw data of immunoblots following separation by SDS-PAGE from Figure S2B.

(**D**) Raw data of Coomassie stained gels following separation by SDS-PAGE from Figure S3B.

(**E**) Raw data of immunoblots following slot-blot analysis from Figure S3F-H.

(**F**) Raw data of immunoblots following slot-blot analysis from Figure S3I-M-.

(**G**) Raw data of immunoblots following separation by SDS-PAGE from Figure S4A.

(**H**) Raw data of immunoblots following separation by SDS-PAGE from Figure S5A.

(**I**) Raw data of immunoblots following separation by SDS-PAGE from Figure S6A.

**A.** Immunoblot from Figure 3F

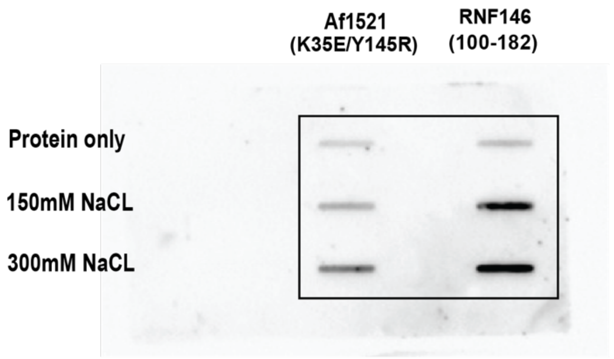

**B.** Immunoblot from Figure 4F (SDS-PAGE)

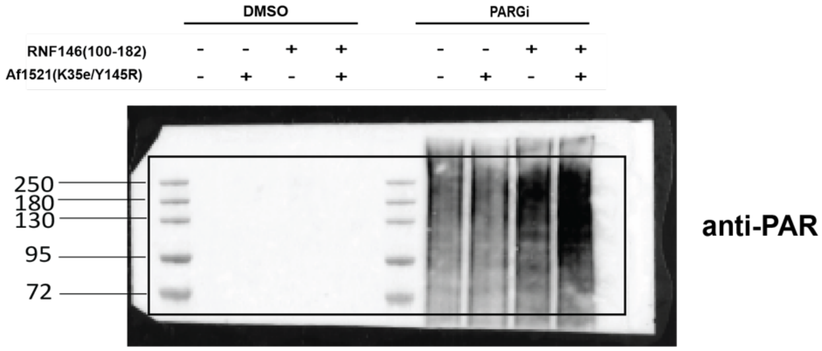

**C.** Immunoblot from Figure S2B (SDS-PAGE)

**U2OS cells**

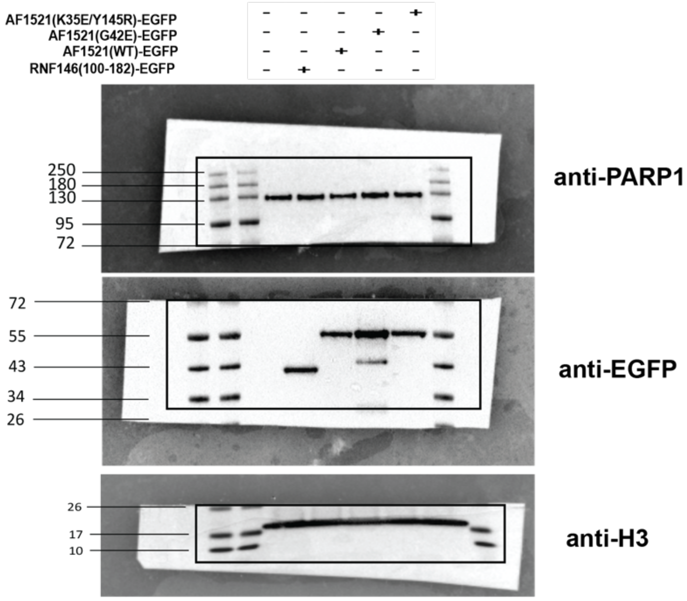

**C.** Immunoblot from Figure S2B (SDS-PAGE) (continued)

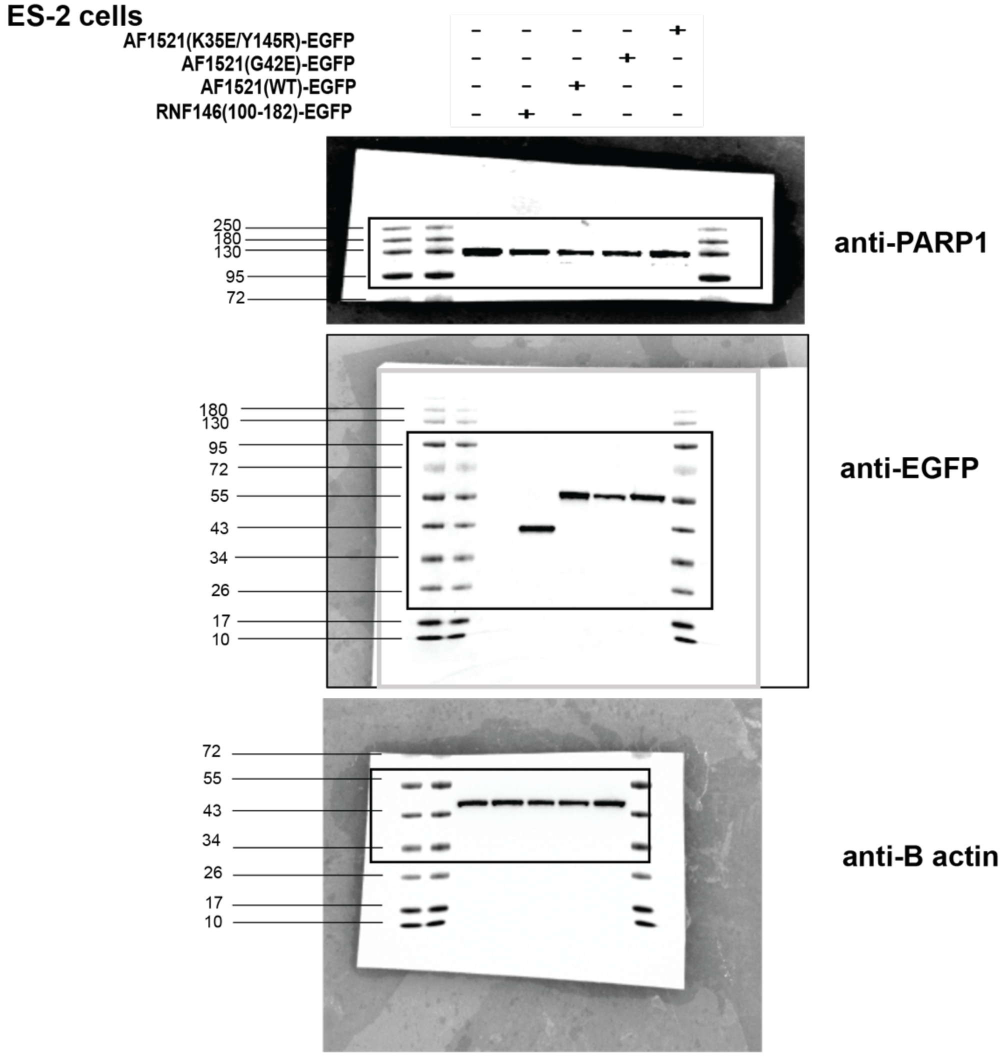

**D.** Coomasie Blue stained gels from Figure S3B (SDS-PAGE)

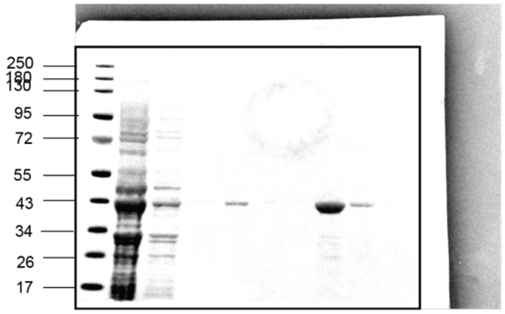

**Purification of GST-RNF146(100-182)**

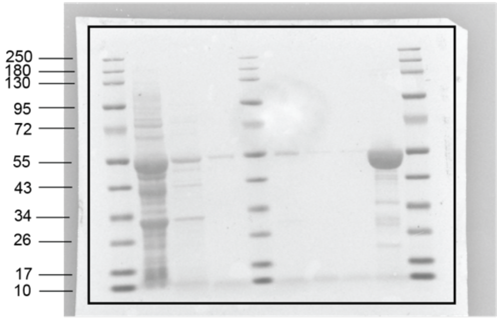

**Purification of GST-Af1521(WT)**

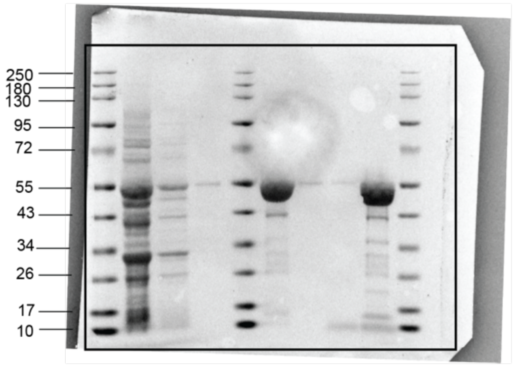

**Purification of GST-Af1521(K35E/Y145R)**

**E.** Immunoblots from Figure S3F-H (slot-blot)

**GST-WWE**

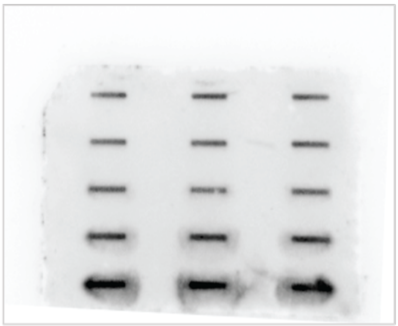

**Anti-GST tag**

**GST-Af1521**

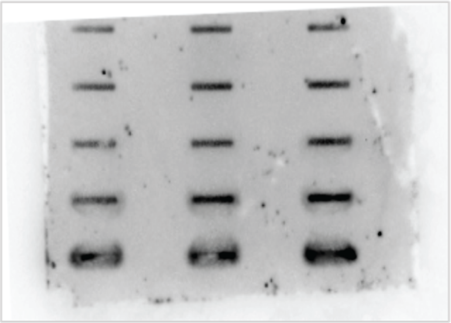

**Anti-GST tag**

**GST-Af1521(K35E/Y145R)**

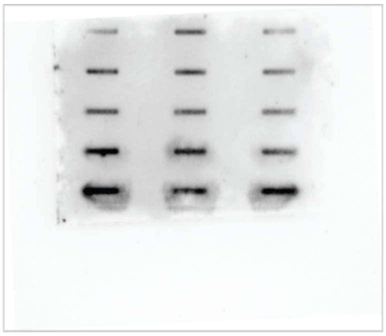

**Anti-GST tag**

**F.** Immunoblots from Figure S3I-M (slot-blot)

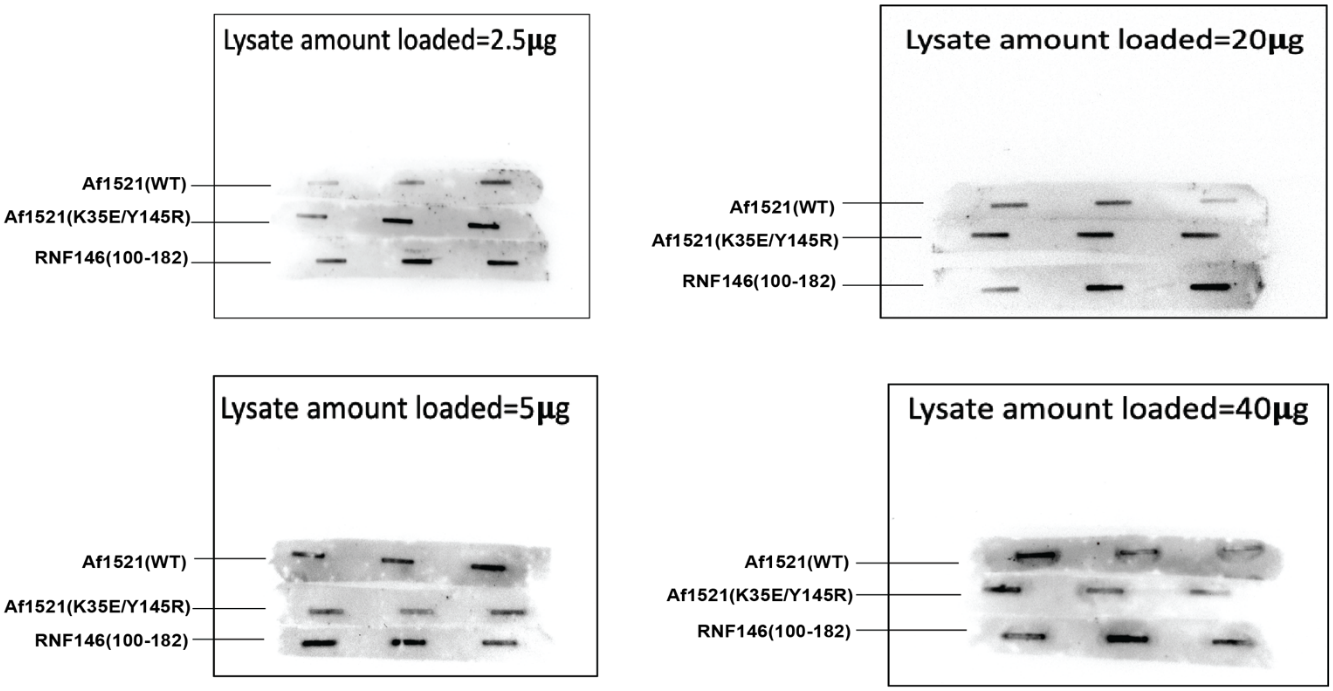

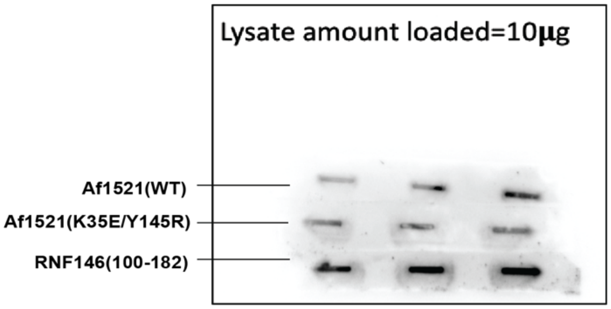

**Each filter is incubated with the protein indicated on the left of each image above. Each filter was then washed and finally probed with the anti-GST tag AB.**

**G.** Immunoblots from Figure S4A (SDS-PAGE)

**U2OS cells**

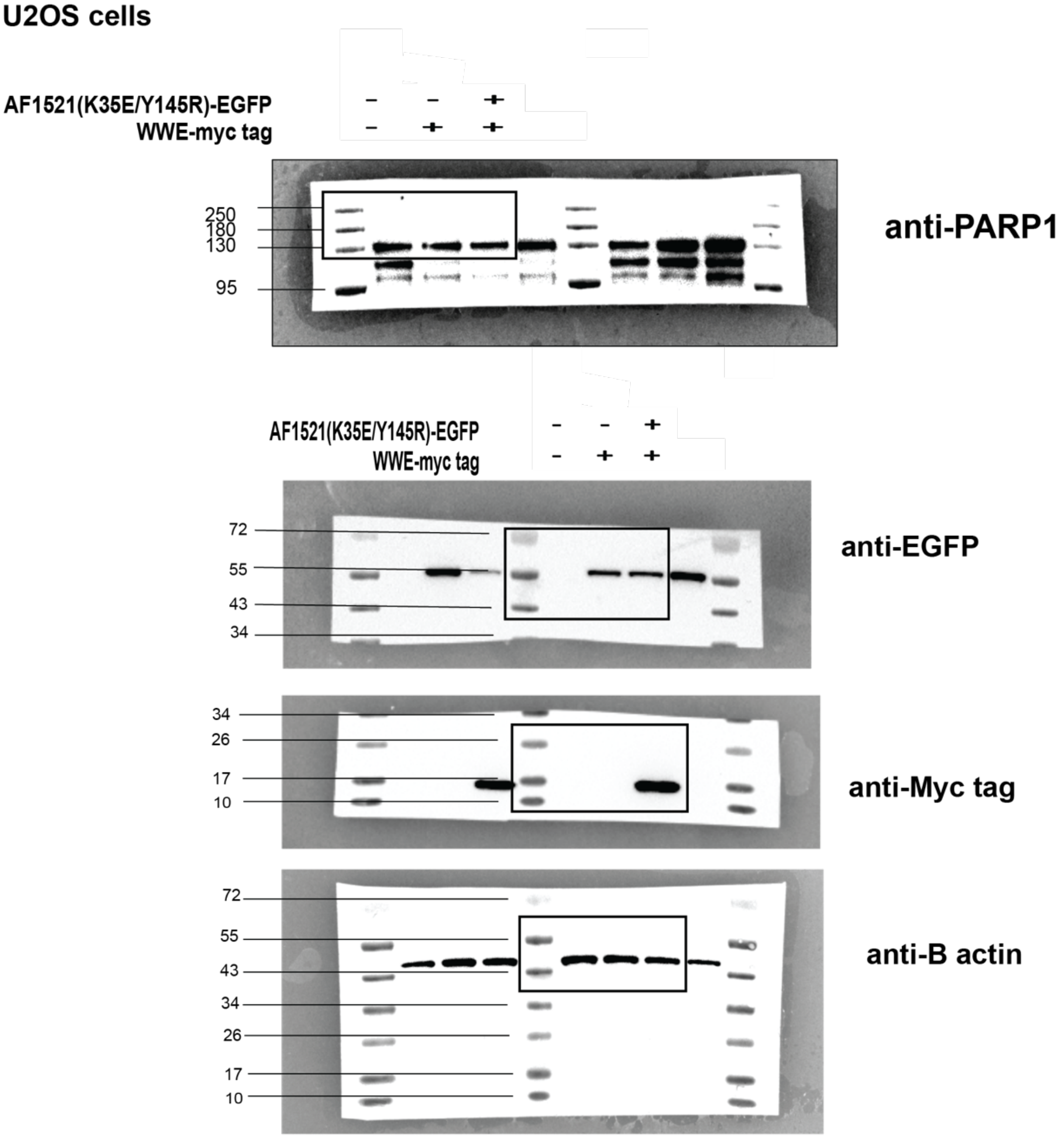

**G.** Immunoblots from Figure S4A (SDS-PAGE) (continued)

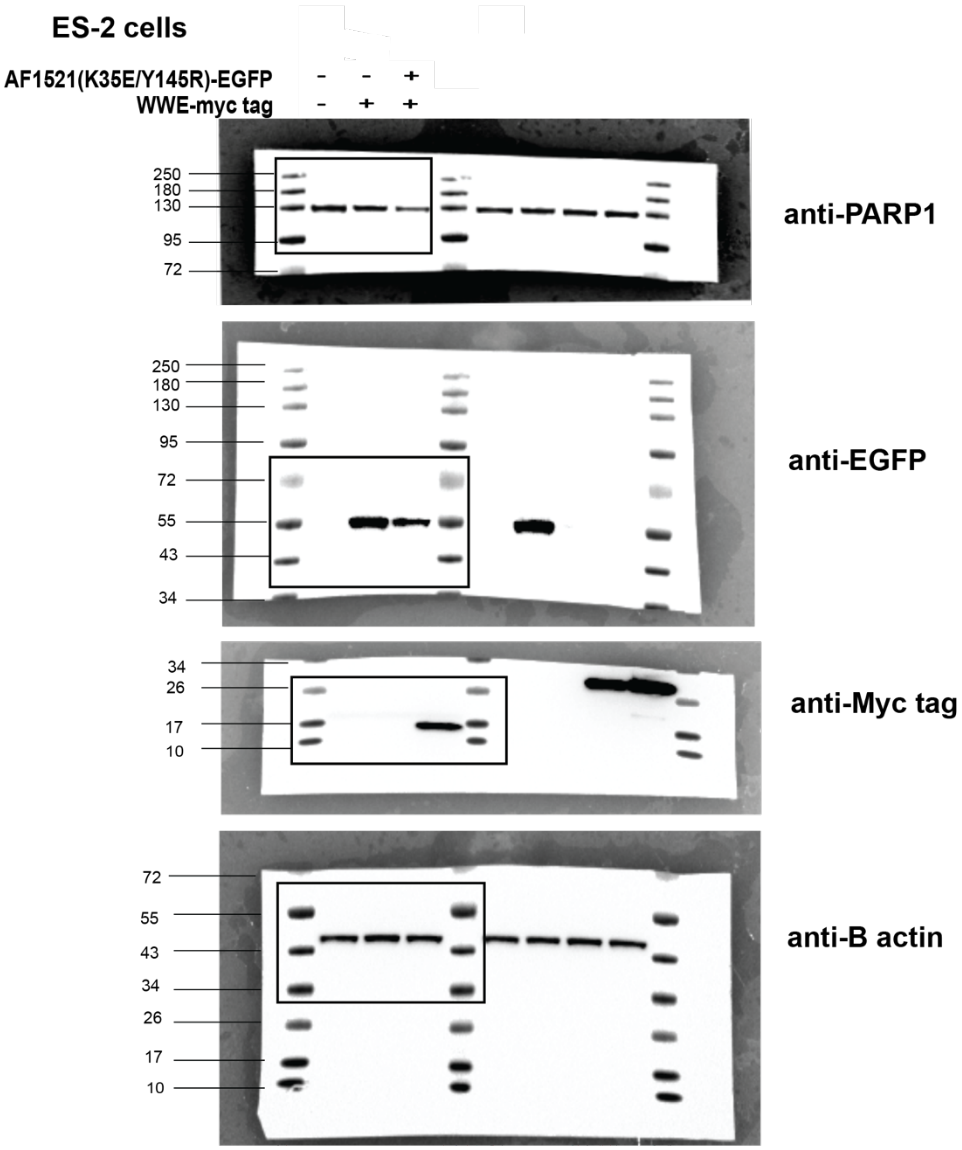

**H.** Immunoblots from Figure S5A (SDS-PAGE)

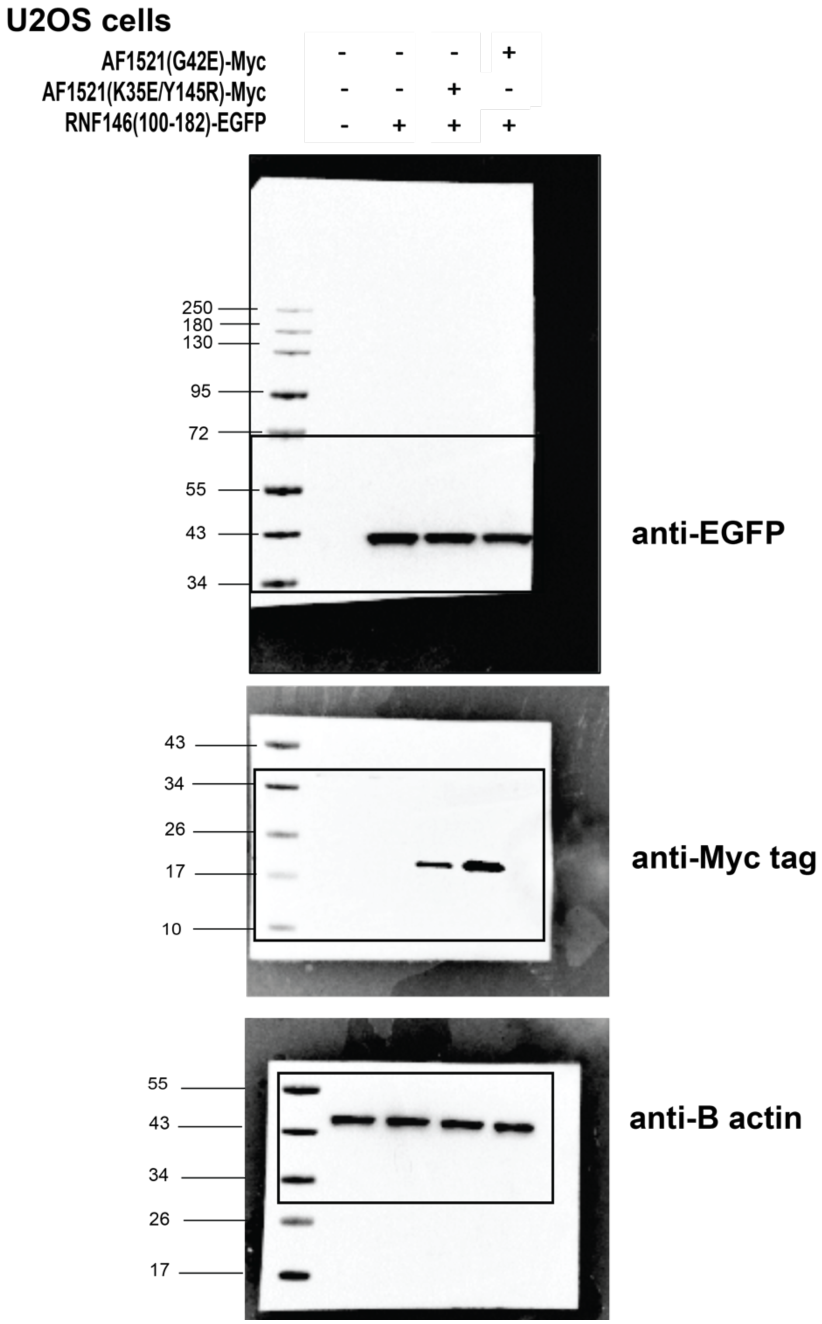

**H.** Immunoblots from Figure S5A (SDS-PAGE) (continued)

**ES-2 cells**

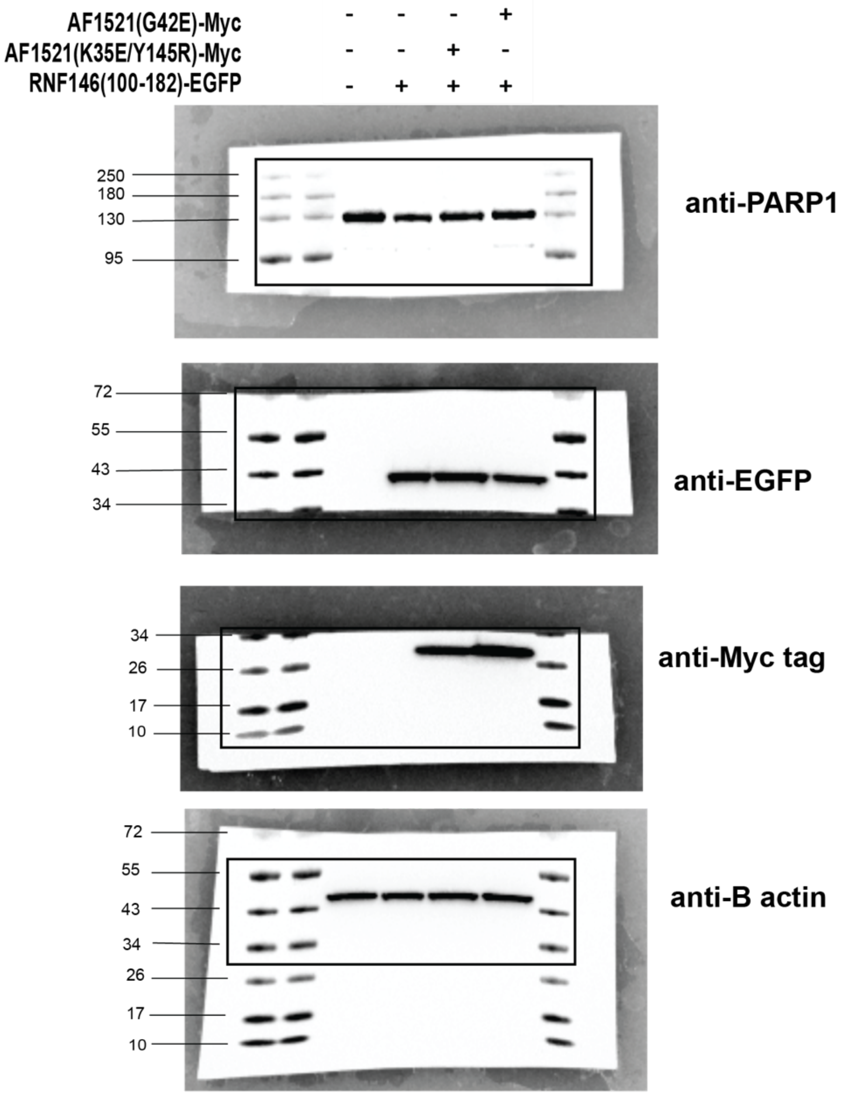

### I. Immunoblots from Figure S6A (SDS-PAGE)

**U2OS cells**

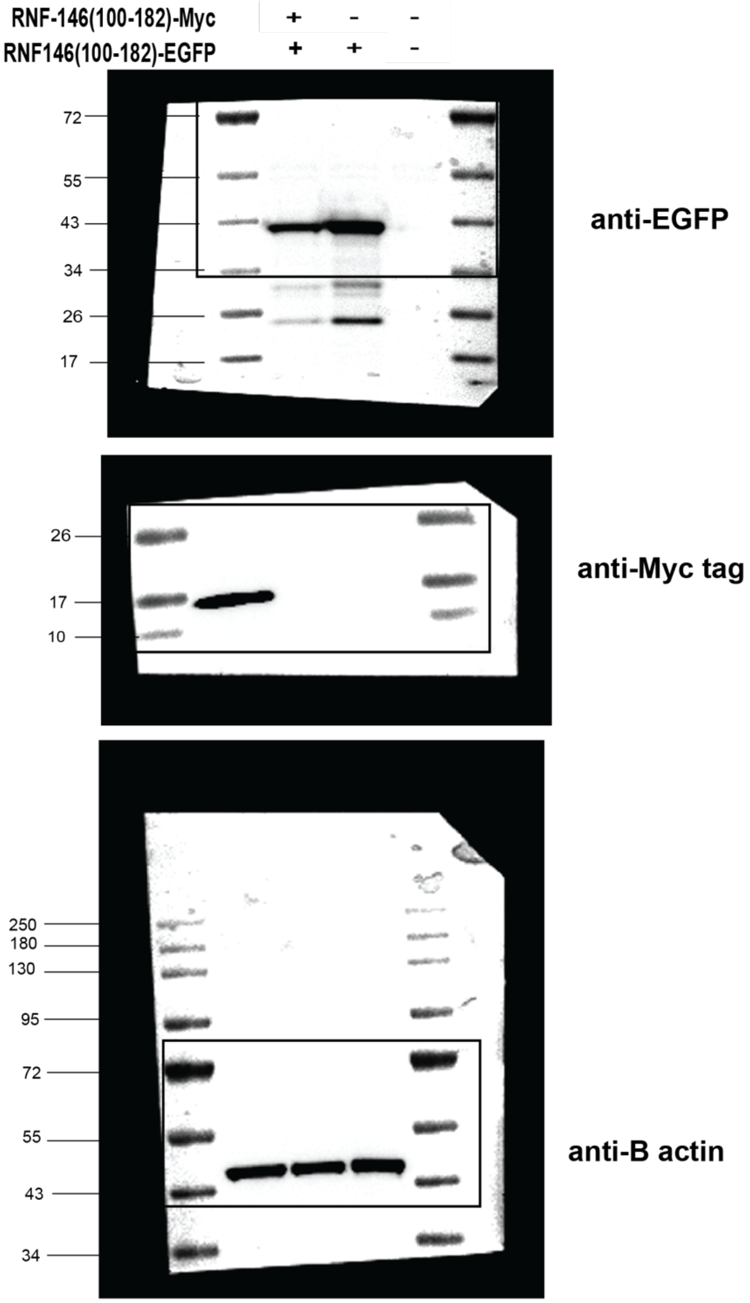

**I.** Immunoblots from Figure S6A (SDS-PAGE) (continued)

**ES-2 cells**

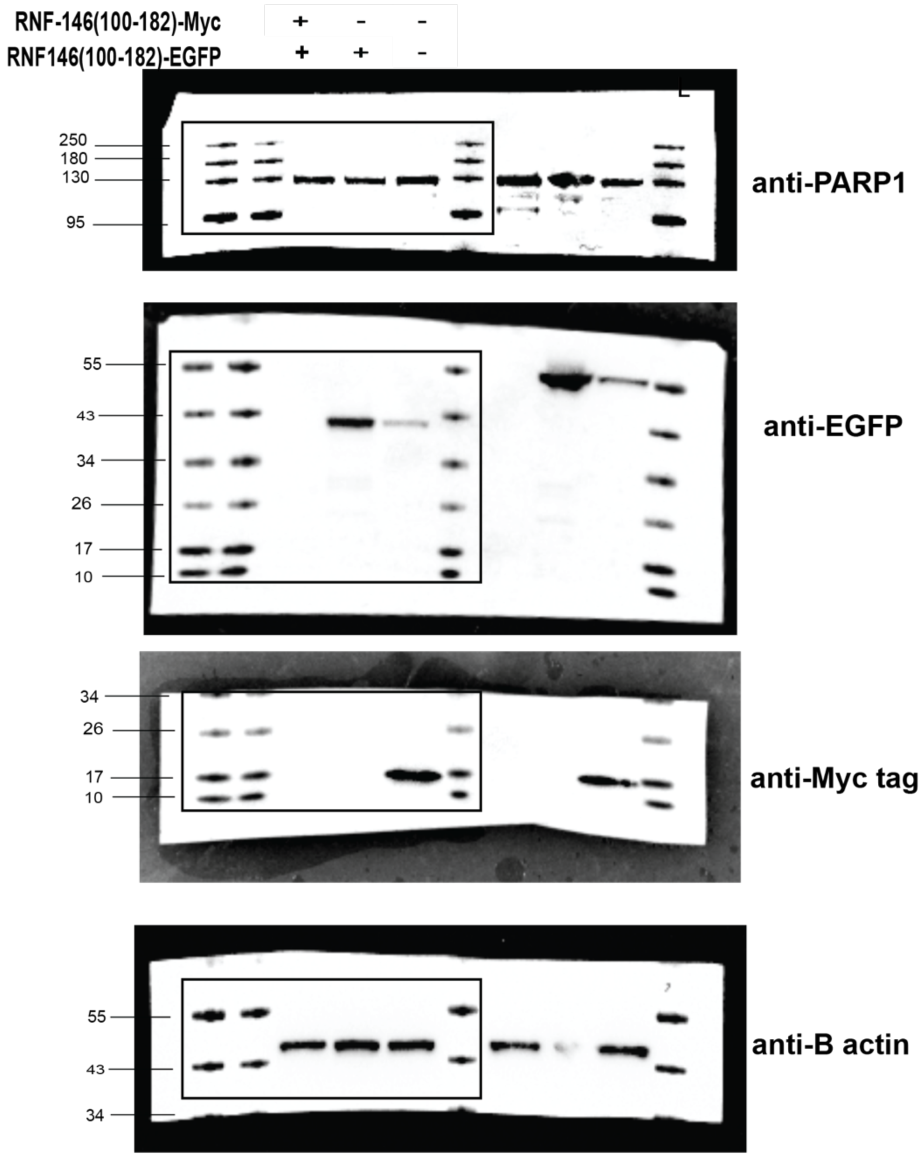
